# The genetic structure of the European black pine (*Pinus nigra* Arnold) is shaped by its recent Holocene demographic history

**DOI:** 10.1101/535591

**Authors:** Guia Giovannelli, Caroline Scotti-Saintagne, Ivan Scotti, Anne Roig, Ilaria Spanu, Giovanni Giuseppe Vendramin, Frédéric Guibal, Bruno Fady

## Abstract

Fragmentation acting over geological times confers wide, biogeographical scale, genetic diversity patterns to species, through demographic and natural selection processes. To test the effects of historical fragmentation on the genetic diversity and differentiation of a major European forest tree and to resolve its demographic history, we describe and model its spatial genetic structure and gene genealogy. We then test which Pleistocene event, whether recent or ancient, could explain its widespread but patchy geographic distribution using population genetic data, environmental data and realistic demographic timed scenarios.

The taxon of interest is a conifer forest tree, *Pinus nigra* (Arnold), the European black pine, whose populations are located in the mountains of southern Europe and North Africa, most frequently at mid-elevation. We used a set of different genetic markers, both neutral and potentially adaptive, and either bi-parentally or paternally inherited, and we sampled natural populations across the entire range of the species. We analysed the data using frequentist population genetic methods as well as Bayesian inference methods to calibrate realistic, demographic timed scenarios.

Species with geographically fragmented distribution areas are expected to display strong among-population genetic differentiation and low within-population genetic diversity. Contrary to these expectations, we show that the current diversity of *Pinus nigra* and its weak genetic spatial structure are best explained as resulting from late Pleistocene or early Holocene fragmentation of one ancestral population into seven genetic lineages, which we found to be the main biogeographical contributors of the natural black pine forests of today. Gene flow among the different lineages is strong across forests and many current populations are admixed between lineages. We propose to modify the currently accepted international nomenclature made of five subspecies and name these seven lineages using regionally accepted subspecies-level names.

**Highlights:**

- The European black pine, *Pinus nigra* (Arnold), has a weak spatial genetic structure.
- Gene flow among populations is frequent and populations are often of admixed origin. Current genealogies result from recent, late Pleistocene or Holocene events.
- Seven modern genetic lineages emerged from divergence and demographic contractions.
- These seven lineages warrant a revision of subspecies taxonomic nomenclature.

## 1. Introduction

Species with geographically fragmented distribution areas are expected to display strong among-population genetic differentiation and low within-population genetic diversity, similarly to biogeographic islands (Mac Arthur & Wilson 1967). This is because gene flow will necessarily be low among long-separated fragments while isolated fragments, if small enough, will lose diversity because of genetic drift and inbreeding (Young et al., 1996). Such patterns are often found in the wild, both in animals and in plants and both when distribution areas are large or small (e.g. Riginos & Liggins 2013; Young et al., 1996). However, examples also abound in both animals and plants, where, despite fragmentation, among population genetic differentiation remains low or modest (e.g. the European mountain pine *Pinus mugo* (Heuertz et al., 2010) or the North American white-footed mouse *Peromyscus leucopus* (Mossman & Waser, 2001)), while within-population genetic diversity is kept at high levels (e.g. the short leaved cedar *Cedrus brevifolia* (Eliades et al., 2011) and see Andrén (1994) for a review).

The climate cycles of the Pleistocene have reshuffled species geographical distributions and genetic diversity patterns (Hewitt 1999; Petit et al., 2003). The imprint left on their current genetic diversity pattern is variable and depends greatly on their life history traits and their ecological preference. Species with ecological niches translating to mid-elevation (thus isolated or patchy) distributions on mountains during the Holocene (the interglacial period of today) were probably widespread at lower elevation repeatedly during the many glacial periods of the Pleistocene (Feurdean et al., 2012), offering several opportunities for gene flow to occur unrestrictedly. In such species, current day patchiness and isolation may not indicate strong genetic differentiation and, a fortiori, the onset of speciation events (see Futuyma (2010) for factors constraining genetic divergence).

*Pinus nigra* Arnold, the European black pine, belongs to the section *Pinus* and the subgenus *Pinus* of the Pinaceae family (Eckert & Hall 2006). It is characterized by a wide, discontinuous distribution area, which spreads from isolated occurrences in North Africa to the Northern Mediterranean and eastwards to the Black Sea and Crimea (Gaussen et al., 1964; Barbéro et al., 1998; Isajev et al., 2004). The European black pine can also grow and adapt to several different soil types and topographic conditions supporting a wide variety of climates across its geographic range. It grows at altitudes ranging from sea level to two thousand meters, most commonly between 800 and 1,500 m. The European black pine is one of the most economically important native conifers in southern and central Europe and one of the most used species in European reforestation programs since as early as the 19th century and throughout the 20th century (Isajev et al., 2004). Several geographic subspecies are described but its taxonomy is still considered as unresolved (Rubio-Moraga et al., 2012).

*Pinus nigra* has a large, but highly fragmented geographic distribution area (Figure 1). Yet, what is known of its genetic diversity is that of a typical temperate forest tree species, with high within- and low among-population genetic diversity (Fady & Conord 2010). The genetic structure data of the European black pine contradict what is expected from a habitat fragmentation perspective, where patchy distributions equate to high ecological and evolutionary divergence (Young et al., 2006). One of the favoured explanation for this *a priori* contradiction is a historically high gene flow (dispersal events over millennia of colonization) over long distances (Kremer et al., 2012). In forest trees, long generation time and late sexual maturity increase the effect of gene flow (Austerlitz et al., 2000). The population genetic pattern of the *Pinus nigra* biogeographical islands of today (particularly at the southern and western edges of the species) should thus be the result of relatively recent fragmentation.

**Figure 1:**
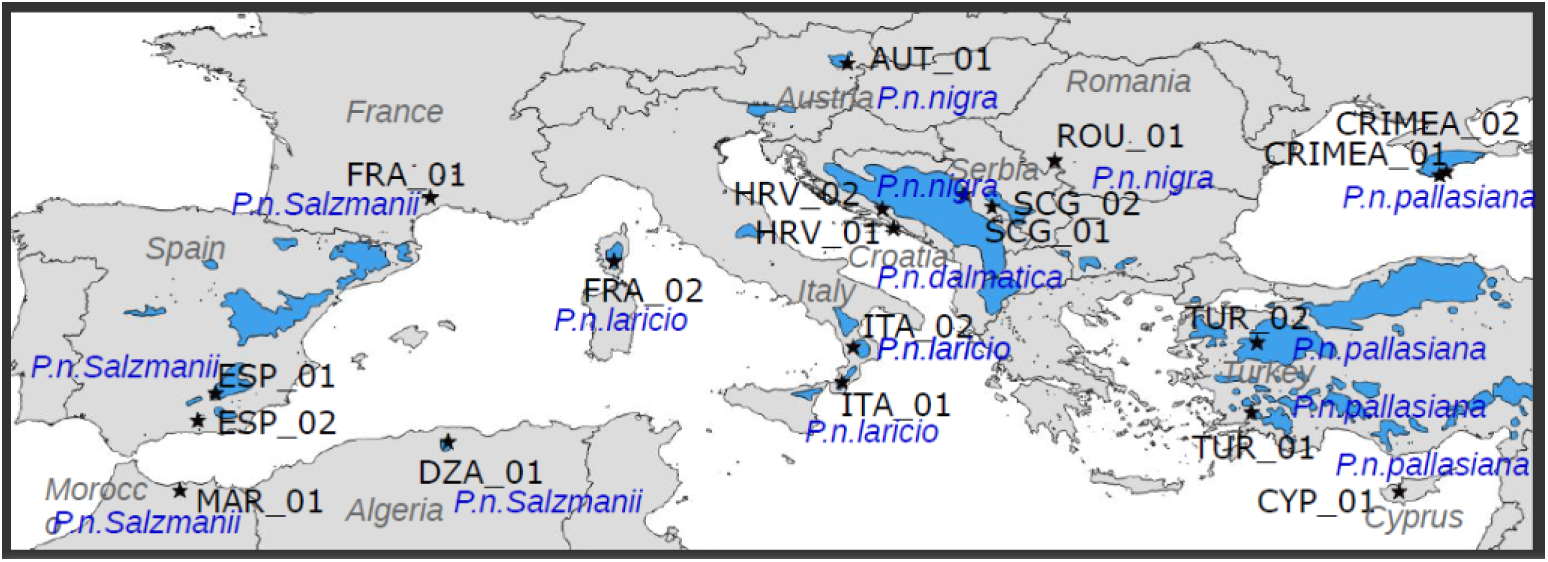
Native geographic distribution area of Pinus nigra (in blue, from Isajev et al., 2004) and location of sampled populations with their code names (black stars) and subspecies name according to the nomenclature of Catalogue of life (http://www.catalogueoflife.org/).

However, widely contradicting evolutionary scenarios have been proposed for explaining the genetic diversity and structure of the European black pine, from very recent Holocene or Quaternary divergence (Rafii & Dodd 2007) to pre-Pleistocene divergence (Naydenov et al., 2016; 2017).

The goal of this study is to test the effects of historical fragmentation on the genetic diversity and differentiation of the European black pine and to resolve its biogeographic history. For this, we describe and model its spatial genetic structure and we test whether its large and patchy distribution results from recent Holocene or earlier Pleistocene events using population genetic data and realistic demographic, timed scenarios. We use a set of different genetic markers, both neutral and putatively adaptive, and both bi-parentally and uni-parentally (cytoplasmic) inherited. Although population genetic studies traditionally use selectively neutral genetic makers, loci influenced by selection are also particularly helpful for assessing relative differences in levels of gene flow, especially in high gene flow species (Guichoux et al., 2013).

We sampled natural populations across the entire range of the species. We discuss why our results differ from, or concur with, the few previous molecular studies carried out at a similar scale on this species (Nikolić & Tucić 1983; Naydenov et al., 2016; 2017; Rafii & Dodd 2007). We also considered how habitat suitability might have affected demography by correlating climate variables at different Pleistocene ages with genetic diversity estimates under the assumption that harsher climate conditions and thus declining habitat suitability, would produce demographic contractions with observable signatures in the genetic data (Conord et al., 2012).

## 2. Materials and Methods

### 2.1 -Sampling and DNA extraction

Needles were collected from 19 different locations across the Mediterranean Basin and southern Europe where black is naturally occurring (Table 1). Populations were represented by 12 individuals each, with the exception of HRV-01 for which 16 individuals were collected. The 19 populations selected cover the full extent of the geographic distribution of *Pinus nigra* as well as its taxonomic diversity (Table 1). DNA was extracted from needles using the DNeasy 96 Plant Kit (QIAGEN, Germany) at the INRA molecular biology laboratory of Avignon, France. Microsatellites data analyses, both chloroplast (cpSSRs) and nuclear (nSSRs), were carried out on the 19 selected populations, while gene sequencing was performed on 18 populations

**Table 1:** Description of the sampled Pinus nigra populations

| Population code* | Population ID | Subspecies** | Country | Samples origin | Latitude | Longitude |
| --- | --- | --- | --- | --- | --- | --- |
| ITA-01 | Aspromonte | P.n. laricio | Italy | Common garden | 38°14' N | 16°00' E |
| TUR-01 | Mugla | P.n. pallasiana | Turkey | Common garden | 37°20' N | 28°24' E |
| HRV-01 | Peljesac | P.n. dalmatica | Croatia | Natural population | 42°52'07" N | 17°33'03" E |
| DZA-01 | Djurdjura | P.n. salzmannii | Algeria | Natural population | 36°27' N | 4°06' E |
| CYP-01 | Cyprus | P.n. pallasiana | Cyprus | Natural population | 34°57' N | 32°50' E |
| FRA-01 | St Guilhem | P.n. salzmannii | France | Natural population | 43°46' N | 3°34' E |
| SCG-01 | Studenica | P.n. nigra | Serbia | Common garden | 43°29' N | 20°32' E |
| FRA-02 | Valdoniello | P.n. laricio | France | Common garden | 41°51' N | 9°06' E |
| AUT-01 | Skie 07 | P.n. nigra | Austria | Natural population | 47°46'00" N | 16°11'00" E |
| SCG-02 | Skie 12 | P.n. nigra | Serbia | Natural population | 43°49'39" N | 19°35'22" E |
| HRV-02 | Skie 14 | P.n. dalmatica | Croatia | Natural population | 43°26'00" N | 17°13'00" E |
| ITA-02 | Skie 19 | P.n. laricio | Italy | Natural population | 39°18'08" N | 16°20'22" E |
| ESP-01 | Skie 33 | P.n. salzmannii | Spain | Natural population | 37°55'00" N | 02°55'00" W |
| TUR-02 | Skie 46 | P.n. pallasiana | Turkey | Natural population | 39°25'50" N | 28°34'10" E |
| ROU-01 | Skie 53 | P.n. nigra | Romania | Natural population | 44°52'57" N | 22°25'40" E |
| CRIMEA-01 | Skie 54 | P.n. pallasiana | Ukraine / Russia | Natural population | 44°25'39" N | 34°03'05" E |
| ESP-02 | 72 | P.n. salzmannii | Spain | Natural population | 37°5'35" N | 3°27'33" W |
| MAR-01 | TAT | P.n. salzmannii | Morocco | Natural population | 35°01'02" N | 4°00'36" W |
| CRIMEA-02 | CRIM | P.n. pallasiana | Ukraine / Russia | Common garden | 44°33' N | 34°17' E |

\* Population codes are based on the ISO 3166-1 standard, which identify countries and territories using a three-letter combination. Given the current geopolitical tensions in Crimea, we decided to code populations from there not using the ISO code (UKR for Ukraine or RUS for the Russian Federation).
\*\* The nomenclature follows Catalogue of life (http://www.catalogueoflife.org/) which recognizes five subspecies: *Pinus nigra* subsp. *salzmannii* (Dunal) Franco, *Pinus nigra* subsp. *laricio* (Poir.) Maire, *Pinus nigra* subsp. *pallasiana* (Lamb.) Holmboe, *Pinus nigra* subsp. *nigra* J.F. Arnold and *Pinus nigra* subsp. *dalmatica* (Vis.) Franco.

### 2.2 -Genotyping and sequencing

Four paternally-inherited cpSSRs, previously identified as polymorphic in black pine, were selected for this study (Appendix S1, Table S1.1 in Supporting Information, Vendramin et al., 1996). The fourteen bi-parentally inherited nSSRs used were those characterized for black pine by Giovannelli et al. (2017) (Table S1.2 in Appendix S1). The fourteen putatively adaptive nuclear genes (called candidate genes hereafter) selected for this study were those that could be transferred from *P. taeda* (Mosca et al., 2012) in a preliminary test (Table S1.3 in Appendix S1). Finally, we also used four organelle genes that can reveal sub-species level differences in widely distributed and taxonomically complex species (Kress & Erickson 2007, Table S1.4 in Appendix S1).

### 2.3 -Data analysis

#### 2.3.1 -Genetic diversity and differentiation estimates: cpSSR and nSSR

We used MICRO-CHECKER (Oosterhout et al., 2004) to estimate the presence of null alleles among the 14 selected nSSRs performing 1000 randomizations. We then used ML-NullFreq (Kalinowski & Taper, 2006) to calculate the frequencies of the null alleles. This two-step method minimizes the false negative rate of null allele detection (Dąbrowski et al., 2014). When the presence of a null allele was detected, genotypes were corrected using MICRO-CHECKER. GenAlEx v6.5 (Peakall & Smouse, 2012) was used to assess genetic diversity as the effective number of alleles (*A*), and observed (*H*_*O*_) and expected (*H*_*E*_) heterozygosity. Genetic differentiation between populations was estimated using *GST* (Nei 1973), the “standardized” measures *G’ST* (Hedrick 2005) and Jost’s D (Jost 2008) following the recommendations for correction of sampling bias of Meirmans & Hedrick (2011). FSTAT V2.9.3.2 (Goudet 2001) was used to calculate rarefied allelic richness. Genepop v4.2.1 was used to test for genotypic disequilibrium among the 14 nSSR loci using likelihood ratio statistics (Rousset, 2008) and default Markov chain parameters.

#### 2.3.2 -Genetic diversity and differentiation estimates: Genes

PHASE v.2.1.1 included in DnaSP v5 (Librado & Rozas, 2009) was used to infer haplotypes. Linkage disequilibrium between pair of SNPs within each gene and within each population was inferred from the exact test of linkage disequilibrium (Raymond & Rousset, 1995) available in Arlequin v3.5.2.2 (Excoffier & Lischer, 2010). The effective number of alleles (*A*), observed (*H*_*O*_) and expected (*H*_*E*_) heterozygosity and *F*_*ST*_ were calculated for each population also using Arlequin v3.5.2.2.

#### 2.3.3 -Phylogeographic patterns

SPAGeDi (Hardy & Vekemans, 2002) was used to calculate and compare *F*_*ST*_ and *R*_*ST*_ for nSSRs, and *F*_*ST*_ and *N*_*ST*_ for cpSSR haplotypes and genes. *R*_*ST*_ and *N*_*ST*_ are analogues of *F*_*ST*_ but take into account the similarities between alleles (Goldstein & Pollock, 1997, Pons & Petit, 1996). The tests were performed globally and within nineteen (eighteen for genes) classes of geographic distances equalling the number of populations analysed. For adaptive genes, the analysis was performed considering each gene separately whereas for nSSR and cpSSR the analysis considered all loci together. The presence of a spatial genetic structure was tested by permuting locations (10,000 permutations). The presence of phylogeographic signals was tested by permuting (10,000) microsatellites allele sizes for nSSRs and by permuting rows and columns of distance matrices between alleles for cpSSRs and genes (Hardy et al., 2003; Pons & Petit 1996).

#### 2.3.4 -Range wide structure

To test for the presence of population genetic structure without *a priori* on the geographic origin of the individuals, we performed a Bayesian clustering approach using the software STRUCTURE v2.3 (Pritchard et al., 2000; Falush et al., 2003). An admixture model for which individuals may have mixed ancestry and a correlated allele frequency model in which frequencies in the different populations are likely to be similar (due to migration or shared ancestry) were considered. The analysis was performed including together nSSRs, cpSSR and genes. Five independent runs for each K value ranging from 1 to 20 were performed after a burn-in period of 5×10^5^ steps followed by 1×10^6^ Markov chain Monte Carlo replicates. The most likely number of clusters (K) and the rate of change of L(K) between successive K values were estimated following Evanno et al. (2005) using the web application StructureHarvester (Earl & von Holdt, 2012). Results from five runs for the most likely K were averaged using the software CLUMPP (Jakobsson & Rosenberg, 2007) with Greedy algorithm and 1 x 10^4^ iterations and visualized using the DISTRUCT software (Rosenberg, 2004).

#### 2.3.5 -Demographic inference

DIYABC v2.1.0 (Cornuet et al., 2014) was used to infer the species’ demographic history within an approximate Bayesian computation (ABC) framework. Among the 19 sampled populations of *P. nigra*, only a subset was used for the demographic inference. The selection of the populations was performed based on the results of STRUCTURE, which identified seven evolutionary lineages. The distribution of the seven genetic lineages was strongly structured: nine populations were homogeneous while the other ten were strongly admixed, suggesting the presence of past or continuous gene flow. Since the latter is not modelled by DIYABC, we focused on the past history reconstruction of both divergence and admixture events. To simplify the historical reconstruction, we first focused on the divergence between the seven lineages.

For this first analysis, seven *P. nigra* populations (12 individuals each) were selected which best represented the identified lineages and contained the lowest level (between 9% and 19%) of admixed individuals: ITA-01 (*P. n. laricio*, Italy), HRV-01 (*P. n. dalmatica*, Croatia), DZA-01 (*P. n. salzmannii*, Algeria), CYP-01 (*P. n. pallasiana*, Cyprus), FRA-01 (*P.n. salzmannii*, France), SCG-02 (*P.n. nigra*, Serbia) and CRIMEA-01 (*P.n. pallasiana*, Crimea) (Table 1). The choice of the tested scenarios is described in details in Table S2.1 of Appendix S2 and shown in Figure 2.

**Figure 2:**
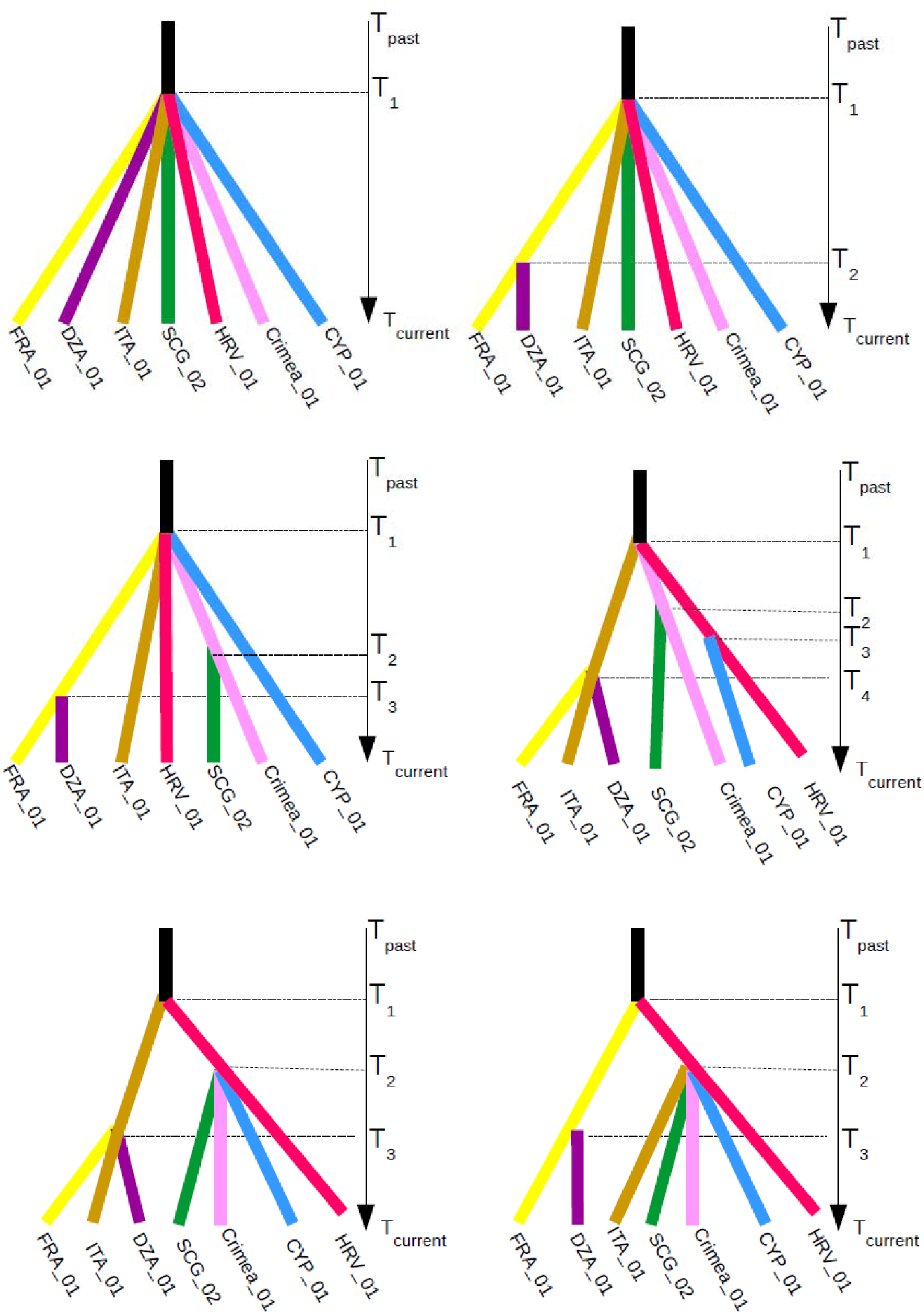
Demographic scenarios tested to infer past range-wide divergence events between seven populations of Pinus nigra.

In a second step, the historical reconstruction focused on the admixed populations. Rather than inferring past demographic parameters from a single tree composed of seven branches and later events of admixtures, we ran separately five different scenarios of admixture events which best explained current population genetic composition. The details of the DIYABC analysis (prior distributions, evaluation of the simulations, selection of the most likely scenario, estimation of the demographic parameters, estimation of the bias and the precision of parameters estimation) are given in Table S2.1 and Figure S2.2 of Appendix S2.

All scenario analyses were performed using the R environment (R Core Team, 2015).

#### 2.3.6 -Correlations between Genetic Diversity and Environmental features

We computed correlations to test whether environmental characteristics could influence neutral genetic diversity patterns through demographic changes. Latitude, longitude and 19 standard bioclimatic variables were downloaded from the WorldClim database (version 1.4, Hijmans et al., 2005) in January 2016 for the present time, the Mid Holocene, the Last Glacial Maximum and the last inter-glacial (Table S3.1 and S3.2 in Appendix S3). We also added three custom-made variables representative of the population isolation (Table S3.1 and S3.2 in Appendix S3).We conducted all analyses in Rstudio (RStudio Team version 1.0.143, 2016).

## 3. Results

### 3.1 -Within population diversity

At nSSR, no significant linkage disequilibrium was detected among loci after the application of the Bonferroni correction at the α = 0.05 confidence level. Significant tests for the presence of null alleles were scattered among loci and populations without a clear mark indicating that a locus or an entire population should be excluded from the analysis (Table S1.5 in Appendix S1). At candidate genes, 173 Single Nucleotide Polymorphisms (SNPs) were detected and significant within-population linkage disequilibrium was found for each gene (Table S1.6 Appendix S1).

All within-population diversity results are described in Table S1.7 of Appendix S1. Organelle DNA gene did not show any diversity at all in any of the preliminary samples we tested. We considered them as monomorphic in all populations throughout our dataset. For candidate genes, the genetic diversity at each locus averaged over eighteen populations varied between 2 and 40 haplotypes (Table S1.6 in Appendix S1). When SSR genetic diversity was estimated per population averaging all loci (Table S1.7 in Appendix S1), gene diversity H_E_ was high both at nuclear genome (between 0.53 and 0.75) and cp genome (between 0.39 and 0.69). A longitudinal cline was observed at both nSSR and candidate genes, with H_E_ and N_A_ values increasing from west to east (Pearson correlation between 0.48 and 0.54, P.value<0.05, Figure S1.1 in Appendix S1).

### 3.2 -Among population structure

Genetic differentiation among populations was significant for all types of genetic markers with strongest values at cpSSR whatever the indices used: Nei’s G_ST_, Hedrick’s G_ST_ or the Jost’s D (Table 2). Nei’s G_ST_ was 0.127, 0.064 and 0.11 for cpSSR, nSSR and candidate genes, respectively. Results from SPAGeDi (Hardy & Vekemans, 2002) showed the presence of an isolation-by-distance pattern at nSSRs and five out of fourteen candidate genes as shown by the positive and significant regression slope of pairwise F_ST_ on geographical distances (Figure S4.1, S4.2 and Table S4.1 in Appendix S4). However, the linear regression slopes of pairwise N_ST_ and R_ST_ on geographical distances were never significant for any markers, indicating that allelic relatedness did not increase with distance, and thus indicating a lack of phylogeographic signal among populations (Figure S4.1, S4.2 and Table S4.1 in Appendix S4).

**Table 2:** Overall estimates of genetic differentiation (Nei’s (1973) GST, Hedrick’s (2005) G’ST and Jost’s (2008) D) at 4 cpSSRs, 14 nSSRs (19 populations) and 14 candidate genes (18 populations).

|  | Gst | G'st | D |
| --- | --- | --- | --- |
| nSSR | 0.064 | 0.262 | 0.209 |
| cpSSR | 0.127 | 0.372 | 0.276 |
| Candidate genes | 0.108 | - | - |

### 3.3 -Range wide population structure

Results derived from STRUCTURE using combined nSSR, cpSSR and nuclear gene data, showed that the most likely number of clusters (K) was 7 (Figure S4.3 and S4.4 in Appendix S4). The first cluster included populations from Algeria (DZA-01) and Morocco (MAR-01) and the second one the French (FRA-01) population. Populations (ESP-01, ESP-02) from Spain where admixed between the two. The third group included populations from Italy (ITA-01) and France (Corsica, FRA-02) and a somewhat admixed Italian population (ITA-02). The Serbian population (SCG-02) and Croatian (HRV-01) populations constituted two different and separate lineages and populations AUT-01 from Austria and ROU-01 from Romania were admixed between the two. The population from Cyprus (CYP-01) constituted a highly differentiated lineage. Populations from Crimea (CRIMEA-01 and CRIMEA-02) and Turkey (TUR-02) constituted the final lineage. Turkish population (TUR-01) was admixed between these last two groups while populations from Serbia (SCG-01) and Croatia (HRV-02) were highly admixed with multiple contributions from different groups (Figure 3).

**Figure 3:**
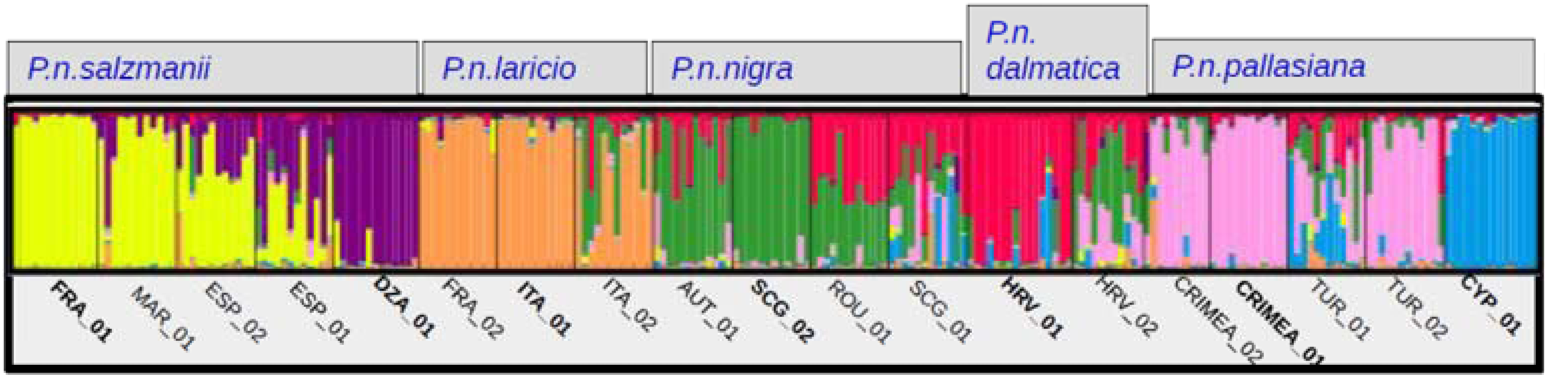
Barplots of ancestry proportions for genetic clusters averaged over 5 runs from K = 2 to K = 7. Each individual is represented by a vertical bar divided into colour segments representing the gene pools identified by the STRUCTURE analysis. Bold types indicate the seven most homogeneous populations (lineages) used in the ABC analysis.

### 3.4 -Demographic inference

#### 3.4.1 -Divergence patterns

The evaluations of the ABC simulations are given in the Figure S2.3 and Table S2.2 in Appendix S2, for both the PCA and a test of rank. Priors represented well the observed summary statistics and empirical data were located within the 95 % of the distribution of the simulated data. The most probable scenario of divergence was scenario 1 with a probability of 0.80 (credible interval between 0.7615 and 0.8311) (Figure S2.4 in Appendix S2). According to this scenario, all populations derived independently from a single common ancestor. The posterior error rate (also called “posterior predictive error”), given as a proportion of wrongly identified scenarios over the 1000 test datasets for both the direct and the logistic approaches, was equal to 0.291 (Table S2.3 in Appendix S2). Estimates of the original parameters (demographic size *N*, divergence time *t*, mutation rate *mu*) are given in Table 3.

**Table 3:**
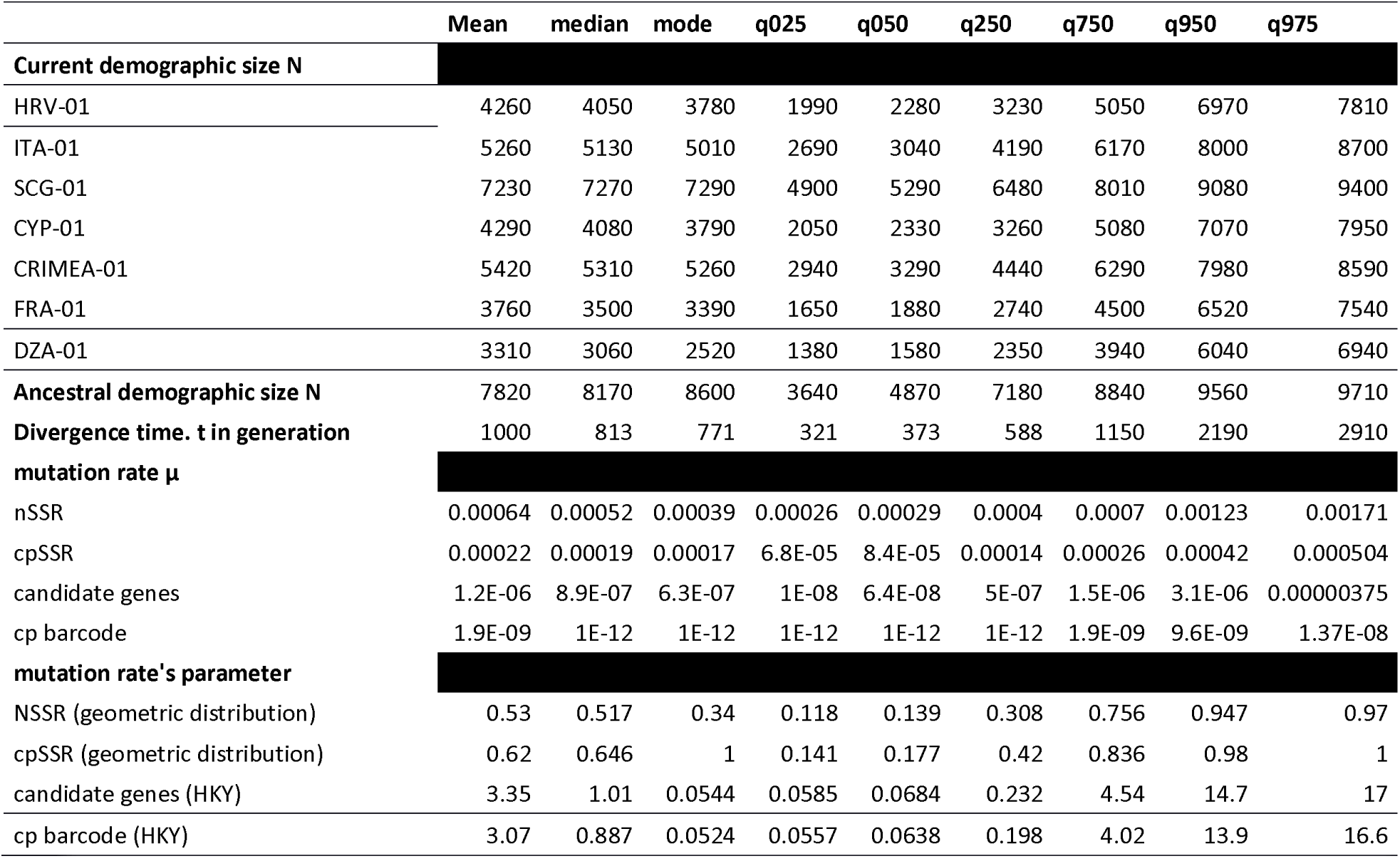
Estimates of the demographic parameters from their posterior distribution (mean, median, mode and quantiles)

The median for the mutation rate was 0.00052 for nSSR, 0.00019 for cpSSR, 8.9.10^-7^ for nuclear genes and 1.10^-12^ for chloroplast DNA sequences. The latter corresponds to the maximum limit of the prior. The absence of polymorphism did not allow estimation of the mutation rate. Median divergence time among the seven lineages was estimated to be 813 generations ago, with a 95% credible interval of 321 – 2910 generations (Table 3). The divergence between the seven lineages occurred approximately 12,195 (4,815 – 43,650) years before present for a generation time fixed at 15 years (i.e. the onset of Holocene) and 24,390 (9,630 – 87,300) years before present for a generation time fixed at 30 years (i.e. thus at the time of Late Glacial Maximum).

Median effective population sizes for scenario 1 were 8,170 for the ancestral population, and between 3,060 (Algeria) and 7,270 (Serbia) for current populations. When comparing the mode of the ratio between current and ancestral population sizes, the ratio N_current_/N_past_ was systematically lower than 1 (between 0.12 and 0.75) (Figure 4) and significantly so for six populations at credible interval of 90% according to the method proposed by Barthe et al. (2016) (Table S2.4 in Appendix S2). Those six populations underwent a contraction at the same time as the split between populations. Only population SCG-02 from Serbia displayed demographic stability over time.

**Figure 4:**
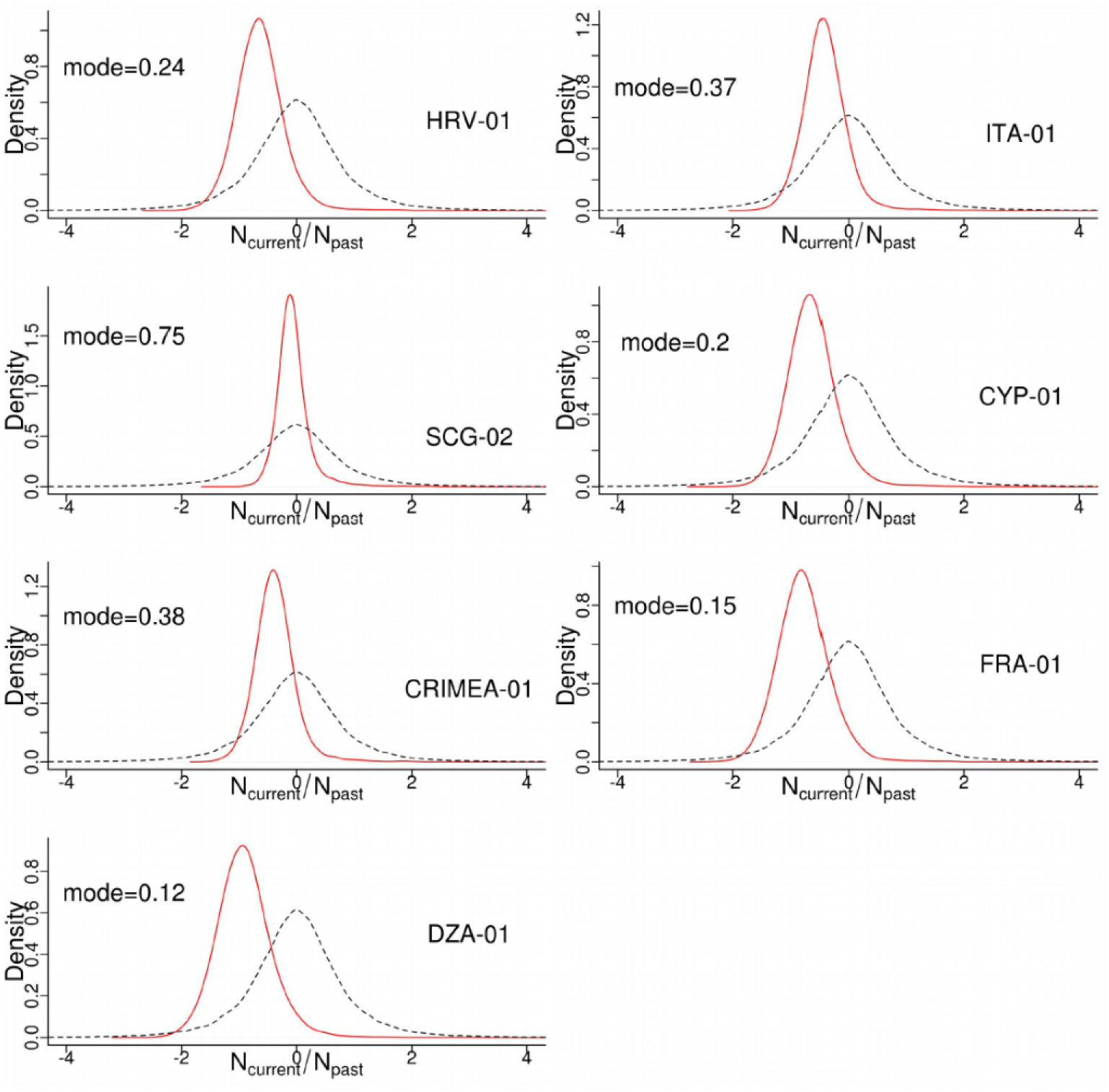
comparison of the mode of the ratio between current and ancestral population sizes (N_current_/N_past_) for seven populations. In black the prior distribution and in red the posterior distribution. This ratio is systematically lower than 1, indicating population contraction.

Results from the accuracy test of parameters estimation (as measured by mean relative bias (MRB), relative root mean square (RMSE) and factor 2) and precision (as shown by coverage values) displayed, globally, reasonable values (Table S2.5 in Appendix S2). The mean relative bias was generally smaller than 1, while factor 2 values ranged from small to moderate according to the estimated parameters. The width of the 50% - 95% credible interval was small, showing that the parameters were reasonably well estimated.

Summary statistics computed after having simulated new data sets from the posterior distribution of parameters obtained under scenario 1 fitted well with the observed summary statistics (Table S2.6 and Figure 2.5 in Appendix S2). We found that none of the eight summary statistics had low tail probability values when applying the model checking option to the scenario 1. The most probable scenario remained the same (S1) with a probability of 0.999 (credible interval between 0.9998 and 1.0000) when it was compared to a scenario allowing demographic resizing along the branches after divergence from the common ancestor (Figure S2.6 in Appendix S2).

#### 3.4.2 -Admixture patterns

The PCA and the test of rank are given in Table S2.7 and Figure S2.6 in Appendix S2. Median time of divergence between all five pairs of populations was between 304 and 1520 generations (Table 4 and Table S2.8 in Appendix S2 for all parameter estimates) overlapping the estimates obtained from the seven populations taken together (Table 3, median was 813 generations). Median time of admixture was between 130 and 503 generations (Table 4) with a confidence interval between 28.9 and 1940 generations. Assuming a generation time between 15 and 30 years, the admixture events between the pairs of lineages occurred between 448 and 58,200 years before present, from the Late Glacial Maximum to after the onset of Holocene. The proportion of test data sets for which the point estimate was at least half and at most twice the true value (factor 2) was rather high (mode of factor 2 > 0.70 for 44 parameters) (Table S2.9 in Appendix S2).

**Table 4:**
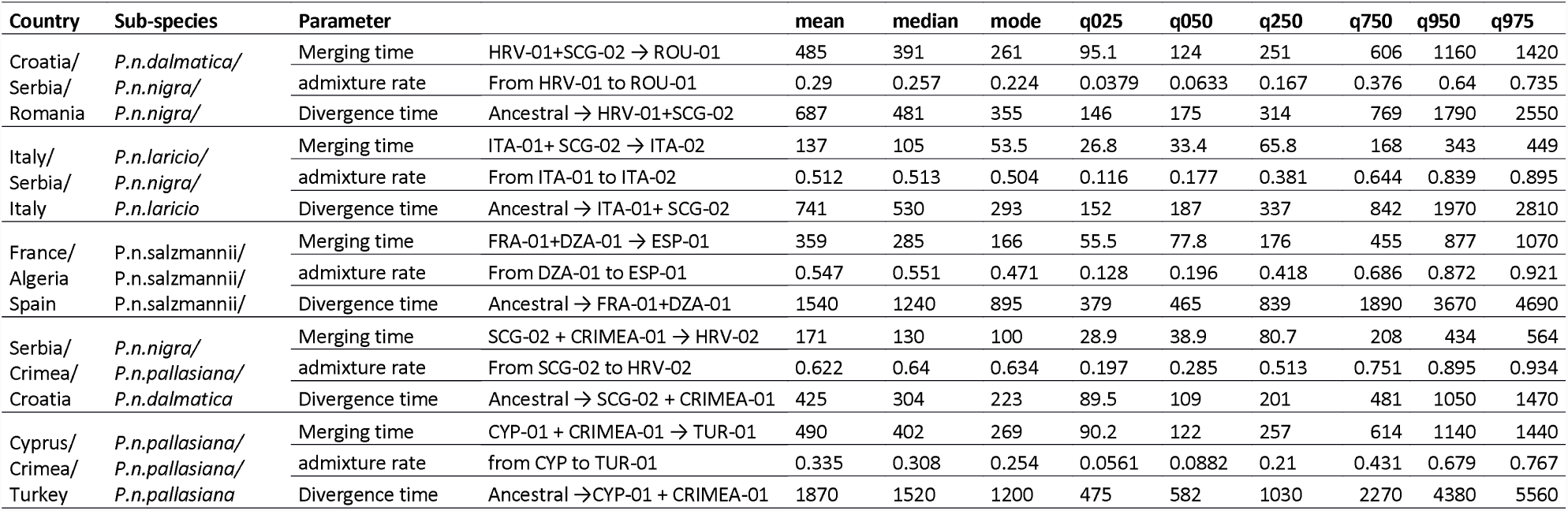
Estimates of the demographic parameters from their posterior distribution (mean, median, mode and quantiles) in the three population models ith admixture

Summary statistics computed after having simulated new data sets from the posterior distribution of parameters fitted well with the observed summary statistics (Table S2.10 and Figure S2.8 in Appendix S2). We found that none of the seven summary statistics had low tail probability values.

#### 3.4.3 -Correlations between genetic diversity estimates and environmental variables

Partial Pearson correlations were not significant. Pearson correlations with a false discovery rate (FDR) lower than 0.1 (i.e. 10% chance that the retained relationships are false positives) were observed between N_A_ and H_E_ at candidate genes and several temperature variables of the Last Glacial Maximum (Table S3.3 in Appendix S3). In addition we observed a FDR <10% between the ratio of demographic size (N_current_/N_past_) and the bioclimatic variable bio10 (mean temperature of the warmest quarter) at Present, Mid Holocene and Last Inter-Glacial. The rank of the explicative environmental variables obtained from the random analyses and the conditional inference tree obtained from the five most important environmental variables (Figure S3.1 of Appendix S3) confirmed the importance of the two LGM bioclimatic variables bio4 (temperature seasonality) and bio10 for candidate genes. The smaller the minimum temperature of coldest month at LGM, the higher was the number of nuclear gene alleles. In the same way, the smaller the temperature seasonality within population, the smaller was the expected heterozygosity (H_E_).

## 4. Discussion and conclusions

Our Bayesian clustering and demographic scenario testing study demonstrates that *Pinus nigra* is composed of seven genetic lineages dating from relatively recent Holocene divergence. A simple, seven-toothed rake-like demographic scenario best explains the biogeographic distribution of present-day black pine populations. The seven lineages diverged from their most recent common ancestor at a very recent time, estimated between the Last Glacial Maximum and the onset of the Holocene.

This scenario agrees with the results of Rafii & Dodd (2007) who also estimated the fragmentation of European black pine lineages to be recent. Focusing on Western Europe, they were able to identify five barriers to gene flow including, similarly to ours, between the Alps and the Calabria - Corsica group, between Corsica and Southern France and between Southern Spain and the Pyrenees. However, our results strongly depart from those of Naydenov et al. (2016, 2017) who identified three genetic groups from three different geographical areas: Western Mediterranean, Balkan Peninsula and Eastern Mediterranean. They estimated their most recent common ancestor to be older than 10 million years and dated the most recent splits between groups in the late Pliocene. These patterns and dates were obtained using cpDNA sequence and cpSSR length polymorphism data, assuming an average mutation rate per generation of 5.6 × 10^−5^. This low mutation rate, identical for both length polymorphism and sequence variation in cpDNA, may be the main reason why our results are so different. As our study uses a suite of different molecular markers and because an old Pliocene divergence time would certainly have left a mutational signature in organelle DNA genes, we are confident that Late Glacial Maximum and/or Holocene demographic events are the most likely explanations for the genetic patterns we observed.

No signature of demographic events older than the last Pleistocene glacial cycle can be found in our data. Environmental correlations with climate data demonstrate the importance of LGM temperature on the genetic diversity and structure of black pine. However, proofs of the presence of black pine ancestors throughout Europe date back to the lower Cretaceous and to the Neogene, between 23 and 2.6 million years ago (Gaussen, 1949). The molecular and fossil calibrated phylogeny of Eckert & Hall (2006) dates the origin of black pine at the Miocene-Pliocene boundary. Following its migration from Polar to Central and Southern European locations during the global cooling of the late Tertiary, black pine was then often found throughout the Pleistocene in charcoal and fossil records in the Mediterranean. From the end of the Pliocene (2.6 million years ago) onwards and during the Pleistocene, it is associated with the sub-Mediterranean flora of Europe (Vernet et al., 1983). *P. nigra* forests played an important ecological role during the Quaternary glacial and interglacial climatic fluctuations in the Western Mediterranean region (Vernet et al., 1983).

Fossil macro-remains indicate that between the Late Pleistocene and the Holocene, *P. nigra* forests had a large distribution (altitude and latitude) in the north-western Mediterranean Basin (e.g. García-Amorena et al., 2011 for the Iberian peninsula). Although *Pinus nigra* is a highland pine, fossil and subfossil remains suggest that it also grew at very low elevation, in coastal areas during cold and dry periods (Postigo-Mijarra et al., 2010). During the warmer, inter-glacial periods of the Pleistocene, it colonized more continental, higher altitudes, as is the case for its current Holocene distribution in the supra-Mediterranean and medio-European vegetation belts (Isajev et al., 2004). It is thus likely that, whatever genetic signature left from fragmentation during the warm intervals of the Pleistocene, it was erased by downward migration during the longer cold climate episodes. Vegetation turnover estimates using fossil records suggest that mid-elevations were the most sensitive altitudinal belts to climate change during the late-glacial and early Holocene (Feurdean et al., 2012).

Human impact is known to have played a role in the past dynamics of *P. nigra* (Vernet, 1986). The decrease of the *Pinus nigra subsp. salzmannii* from the Iron Age onwards and the concomitant development of *Quercus ilex* and *Pinus halepensis*, for example, may be related to fire events, logging and agro-pastoral activities (Rodrigo et al., 2004). The effective size of today’s black pine populations is generally significantly smaller than that of their ancestral population. As resizing after the emergence of current lineages was not sustained by our data, the most likely explanation for this smaller effective population size is not an effect of human impact, rather that of a contraction at the time of lineage splitting.

Admixture among the seven lineages we identified, was found to be frequent in natural populations of black pine, suggesting recent gene flow despite habitat fragmentation. There was no signal of a phylogeographic structure, only that of a weak isolation by distance, confirming the hypothesis of recent differentiation and emergence of distinct lineages. This pattern is reminiscent of that of *P. sylvestris*, a European pine with broadly similar ecological requirements as black pine (Robledo-Arnucio et al., 2005; Pyhäjärvi et al., 2007; Tóth et al., 2017).

The moderately high within-population genetic diversity of black pine is between that of low elevation and high elevation Mediterranean pines (Robledo-Arnuncio et al., 2005; Rafii & Dodd 2007; Fady & Conord 2010). The highest genetic diversity within populations was found in populations originating from the Balkans and Turkey, the geographic area where the most extensive forests of *P. nigra* are found today. Within population genetic diversity shows an east-west decreasing trend, indicating that eastern Mediterranean black pine forests probably found more favourable habitats at the time of lineage splitting (Conord et al., 2012).

All but one populations corresponding to one of the seven lineages identified are located on islands, at isolated or rear-edge locations. It is likely that these locations correspond to glacial refugia. Most of them appear as refugia in the synthesis of Médail & Diadema (2009). To this list of 52 putative refugia of importance for Mediterranean plant species, our study adds the southern Serbian mountains and Crimea.

Finally, the seven lineages we identified can help resolve the debated taxonomy of the European black pine (Rubio-Moraga et al., 2012). Both widely and only regionally accepted subspecies or variety names can identify these lineages (Nyman, 1879; Fukarek, 1958; Debazac, 1964; Vidaković, 1974; Farjon, 2010). The westernmost, North African origin lineage corresponds clearly to the taxonomic entity recognized as *Pinus nigra subsp. mauretanica*. The second westernmost European lineage located in continental France, corresponds to the taxon *Pinus nigra subsp. salzmannii*. These two taxa are often grouped together in several nomenclatures. Moving eastward, the third lineage is located in Corsica (France) and Calabria (Italy) and corresponds to the taxon *Pinus nigra subsp. laricio*. It is worth noting that *Pinus nigra subsp. laricio* from Calabria and from Corsica are often distinguished in the forestry literature (e.g. Roman-Amat & Arbez, 1986), which we could not do here despite our use of uni- and bi-parentally inherited, neutral and putatively selected markers. The fourth lineage corresponds to the taxon *Pinus nigra subsp. dalmatica* on the Dalmatian coast in Croatia. The fifth lineage is found in Central Europe (Serbia) and corresponds to the widely distributed taxon *Pinus nigra subsp. nigra*. The sixth lineage corresponds to the widespread *Pinus nigra subsp. pallasiana* and originates from the Crimean peninsula. Lastly, the easternmost seventh lineage is located in Cyprus and corresponds to a taxon generally included in *Pinus nigra subsp. pallasiana* but sometimes recognized as *Pinus nigra subsp. caramanica*. A list of synonyms of *Pinus nigra* subspecies and varieties is available from the IUCN red list of threatened species website, at: http://dx.doi.org/10.2305/IUCN.UK.2013-1.RLTS.T42386A2976817.en

## 5. Acknowledgements

We thank: O. Gilg, F. Rei, N. Turion and D. Vauthier (INRA UEFM, Avignon, France), J. Rousselet (INRA Orléans, France), E. Kakouris (Cyprus Forest Department, Nicosia, Cyprus), F. Krouchi (University of Tizi Ouzou, Algeria), G. Huber (Bavarian Office for Forest Seeding and Planting, Teisendorf, Germany), S.C. Gonzalez-Martinez (formerly at National Institute for Agronomy, Madrid, Spain) and H. Sbay (Forest Research Centre, Rabat, Morocco) for sample collection. Collection of material was made before the Nagoya Protocol on Access to Genetic Resources and the Fair and Equitable Sharing of Benefits Arising from their Utilization to the Convention on Biological Diversity, was legally implement in Europe and by signatory countries.

## 7. Author contribution

GG, BF and FG conceived the research, GG, AR and ISp produced the data, GG, IS and CSS analysed the data and GG, BF and CSS wrote the text. All authors revised and approved the final version of the manuscript.

