## Supplementary files for "The genetic structure of the European black pine (*Pinus nigra* Arnold) is shaped by its recent Holocene demographic history"

Short title: the European black pine has a recent Holocene history

**Appendix S1: list of genetic markers and within population diversity**

**Table S1.1:** List of the cpSSR primer sequences and PCR and genotyping conditions.

**Table S1.2:** Genetic diversity estimates of nSSR loci and genotyping conditions.

**Table S1.3:** List of the nuclear genes used, and their associated putative function and sequencing conditions.

**Table S1.4:** Sequencing conditions of organelle genes

**Table S1.5:** Frequency of null alleles within population at 14 nSSR loci

**Table S1.6:** Within population haplotypic diversity at nuclear genes. Values were averaged over the eighteen populations

**Table S1.7:** Within population genetic diversity at organelle gene, cpSSRs, nSSRs and nuclear genes

**Figure S1.1:** Linear regression between genetic diversity and longitude of black pine populations.

**Table S1.1:** List of the cpSSR primer sequences and PCR and genotyping conditions. Codes are from Vendramin et al. (1996). Tm is the annealing temperature in °C.

| **cpSSR Code** | **Forward and reverse primer sequences (5' - 3')** | **Tm** | **PCR product size** | **Number of repeat units** |
| --- | --- | --- | --- | --- |
| Pt1254 | CAATTGGAATGAGAACAGATAGG | 57.8 | 74 | (T)17 |
|  | TGCGTTGCACTTCGTTATAG | 57.6 | 72 | (T)15 |
| Pt30204 | TCATAGCGGAAGATCCTCTTT | 58.0 | 145 | (A)12(G)10 |
|  | CGGATTGATCCTAACCATACC | 58.3 | 140 | (A)9(G)8 |
| Pt71936 | TTCATTGGAAATACACTAGCCC | 58.1 | 148 | (T)16 |
|  | AAAACCGTACATGAGATTCCC | 57.9 | 146 | (T)14 |
| Pt87268 | GCCAGGGAAAATCGTAGG | 58.1 | 165 | (T)14 |
|  | AGACGATTAGACATCCAACCC | 58.0 | 167 | (T)10C(T)5 |

The four cpSSRs were combined into one multiplex reaction. PCRs were carried out using the Multiplex PCR Kit (QIAGEN, Germany) with the following amplification conditions: the final volume was optimized to 10 μl and the PCR mix was: 5 μl of the QIAGEN Multiplex PCR Master Mix, 2 μl of Q solution, 2 μl of DNA (10 ng⁄μl) and 0.2 μM primers. The PCR thermal profile was: an initial step at 95 °C for 15 min, followed by 35 cycles at 94 °C for 30 sec, 57 °C for 90 sec and 72 °C for 90 sec, with a final 10 min extension step at 72 °C. PCR products were run on an AB 3730 XL automatic sequencer (Applied Biosystems, USA) with LIZ-600 as internal size standard at the INRA molecular biology laboratory of Avignon, France. Chromatograms were analysed using GeneMapper v4.1 (Applied Biosystems, USA) and two different readers carried out, separately, the cpSSRs scoring in order to validate the data.

Vendramin GG, Lelli L, Rossi P, Morgante M (1996) A set of primers for the amplification of 20 chloroplast microsatellites in Pinaceae. Molecular Ecology 5: 595–598

**Table S1.2:** Genetic diversity estimates of nSSR loci and genotyping conditions.

| **Locus code name** | **A** | **H0** | **He** | **F_IS_** | **F_IS_(p-value)** | **F_ST_** | **F_ST_(P-value)** | **D** | **D (p-value)** |
| --- | --- | --- | --- | --- | --- | --- | --- | --- | --- |
| pn6360 | 9.789 | 0.817 | 0.798 | -0.024 | 1 | 0.106 | 0.001 | 0.379 | 0.001 |
| pn7754 | 9.579 | 0.86 | 0.833 | -0.033 | 1 | 0.084 | 0.001 | 0.325 | 0.001 |
| pn2153 | 3.684 | 0.431 | 0.503 | 0.15 | 1 | 0.224 | 0.001 | 0.261 | 0.001 |
| SPAG_7.14 | 9.737 | 0.862 | 0.829 | 0.168 | 0.023 | 0.099 | 0.001 | 0.417 | 0.001 |
| pn4379 | 9 | 0.802 | 0.775 | 0.029 | 1 | 0.111 | 0.001 | 0.35 | 0.001 |
| pn2246 | 11.105 | 0.766 | 0.85 | 0.134 | 0.559 | 0.102 | 0.001 | 0.47 | 0.001 |
| pn6175 | 11.105 | 0.832 | 0.865 | 0.063 | 1 | 0.086 | 0.001 | 0.426 | 0.001 |
| pn1403 | 8.632 | 0.851 | 0.818 | -0.041 | 1 | 0.091 | 0.001 | 0.339 | 0.001 |
| PHA_6062 | 6.421 | 0.791 | 0.739 | -0.044 | 1 | 0.085 | 0.001 | 0.166 | 0.001 |
| PHA_4783 | 2 | 0.449 | 0.388 | -0.157 | 1 | 0.195 | 0.001 | 0.14 | 0.001 |
| PtTX4001 | 4.842 | 0.657 | 0.705 | 0.072 | 1 | 0.108 | 0.001 | 0.203 | 0.001 |
| pn8747 | 6.895 | 0.763 | 0.769 | 0.013 | 1 | 0.075 | 0.001 | 0.138 | 0.001 |
| PtTX3107 | 2.842 | 0.3 | 0.305 | 0.017 | 1 | 0.161 | 0.001 | 0.068 | 0.001 |
| pn6266 | 5.526 | 0.696 | 0.64 | -0.074 | 1 | 0.126 | 0.001 | 0.202 | 0.001 |

A: number of allele, Ho and He: observed and expected heterozygosities, F_IS_: inbreeding coefficient and its p-value, F_ST_ and D: differentiation indices (with their respective p-values), A description of the loci and amplification conditions can be found in Giovannelli et al. (2017).

For details on multiplex composition, amplification conditions and thermal profile selection, see Giovannelli et al. (2017). PCR products were run on an AB 3730 XL automatic sequencer (Applied Biosystems, USA) with LIZ-600 as internal size standard at the INRA molecular biology laboratory of Avignon, France. The match between simplex and multiplex profiles of samples was checked to control for allele amplification competition and possible allelic drop-out. Chromatograms were analysed using GeneMapper v4.1 (Applied Biosystems, USA) and scoring was done twice, by two different persons, to validate the data.

Giovannelli, G., Roig, A., Spanu, I., Vendramin, G.G. & Fady B. (2017). A New Set of Nuclear Microsatellites for an Ecologically and Economically Important Conifer: the European Black Pine (Pinus nigra Arn.). Plant Molecular Biology Reporter 35, 379–388

**Table S1.3:** List of the nuclear genes used, their associated putative function and sequencing conditions.

| **nuclear gene code name** | **GenBank Accession Number** | **Number of PCR cycles** | **Blastx (A.thalaiana peptides)** |
| --- | --- | --- | --- |
| 0_10162_01 | MG841334 - MG841499 | 35 | pectato liasi |
| 0_10384_02 | MG843122 - MG843336 | 25 | phosphatase2C_PP2C family protein |
| 0_10667_02 | MG841500 - MG841710 | 25 | unknown protein |
| 0_13484_01 | MG843337 - MG843451 | 25 | LBD40 (LOB Domain-Containing Protein 40) |
| 0_13957_02 | MG841711 - MG841903 | 35 | receptor-like protein kinase HSL1 |
| 0_14221_01 | MG841904 - MG842114 | 35 | SRS (Seryl-TRNA Synthetase) |
| 0_16810_02 | MG842115 - MG842318 | 35 | Protein Suppressor of npr1-1, Constitutive 1 |
| 0_18101_02 | MG843452 - MG843627 | 25 | AtRABG3d (Arabidopsis Rab GTPase homolog G3d) |
| 0_2078_01 | MG841159 - MG841333 | 25 | Rho GDP-dissociation inhibitor family protein |
| 0_6293_01 | MG842511 - MG842721 | 25 | unknown protein |
| 0_7916_01 | MG842722 - MG842931 | 25 | EXGT-A4 (Endoxyloglucan Transferase A4) |
| 0_8479_01 | MG842932 - MG843121 | 35 | INT2 (Inositol Transporter 2) |
| 2_1405_01 | MG842319 - MG842510 | 25 | methionine synthase |
| CL4470Ct1 | MG843628 - MG843837 | 25 | PRR1 (Pinoresinol Reductase 1) |

Fourteen putatively adaptive nuclear genes (called candidate genes hereafter) deriving from *P. taeda* (Mosca et al., 2012) could be amplified in *P. nigra* in a preliminary test (not shown) and were selected for this study (Table S1.3 in Appendix S1). Sequencing was carried out at the Institute of Biosciences and BioResources, National Research Council (IBBR-CNR) of Florence, Italy. Due to poor sequence quality, population CRI-02 was excluded from the analysis. PCR amplifications were carried out in an optimized final volume of 14 μl and the PCR mix was: 2.8 μl of 1X GoTaq® Reaction Buffer, 0.2 mM of each dNTP, 1.25 U of GoTaq® G2 DNA Polymerase (Promega, USA), 1 μl of DNA (20 ng⁄μl) and 0.2 μM primers.

The thermal profile for the reaction was: initial denaturation at 94°C for 3 min, followed by 10 cycles (decreasing the annealing temperature 1°C/cycle) at 94°C for 30 s, 60°C for 30 s and 72°C for 40 s, followed by 25 or 35 cycles (depending on the candidate gene analysed) at 94°C for 30 s, 50°C for 30 s and 72°C for 40 s, with a final 10 min extension step at 72°C.

PCR products were quality-checked on 2% agarose gel stained with GelRed (Biotium, USA) and then purified using Multiscreen^®^ filter plates (Merckmillipore) and a vacuum manifold in order to carry out the sequencing reaction. The sequencing mix composition was: 4 µl of purified PCR product, 0.32 µl of Forward or Reverse Primer, 1.5X Sequencing Buffer, 0.5 µl of BigDye® Terminator v3.1 Ready Reaction Mix (ThermoFisher SCIENTIFIC, USA) and RNase-free water for a total volume of 6 µl. The sequencing thermal profile was characterized by a first step of denaturation at 94°C for 1 min followed by 25 cycle at 96°C for 10 sec, 50°C for 5 sec, 60°C for 3 min, with a final step at 10°C for-ever.

Sequencing reactions were then purified by membrane filtration using MultiScreenHTS^®^ HV, 0.45 µm Plates (Merckmillipore) and Sephadex G50 (GE Healthcare) and analysed on an AB 3500 automatic sequencer (Applied Biosystems, USA) using a traditional Sanger DNA sequencing method. Sequences were finally manually aligned, quality checked and edited on CodonCode Aligner 3.7.1 (CodonCode Co., MA, USA); low quality sequences were trimmed. We obtained both the forward and the reverse sequences for each sampled individual, for all the 14 genes with the exception of locus 0_6293_01 for which only forward sequences were of high enough quality to be analysed. Electrophoregrams were visually inspected, low quality sequences were trimmed and insertions and deletions were excluded from the analysis. Forward and reverse sequences were joined into a single sequence using an in-house R script available upon request. The program automatically checks for read inconsistencies between the forward and reverse sequences by providing a summary table of inconsistencies. If after reading verification, no decision was possible, the base was treated as missing data. The sequences were then merged into a single sequence whose heterozygous nucleotide sites are coded using the IUPAC ambiguity codes.

**Table S1.4:** Sequencing conditions of organelle genes

Four organelle genes were selected for this study: matK, rbcL, trnH-psbA (chloroplast DNA genes, paternally inherited in the Pinaceae) and nad5-4 (mitochondrial DNA gene, maternally inherited in the Pinaceae). These organelle genes are often used for barcoding purposes and can reveal sub-species level differences in widely distributed and taxonomically complex species (Kress & Erickson 2007; Ziegenhagen et al., 2005). To check for the presence of potential mutations within these organelle gene regions, a preliminary test was carried out with a reduced set of 25 herbarium samples covering the entire geographic range of the species. DNA was extracted from leaf tissue using the DNeasy 96 Plant Kit (QIAGEN, Germany) at the INRA molecular biology laboratory of Avignon, France. Primer sequences were obtained from Kress & Erickson (2007) (matK and rbcL), from Kress et al. (2005) (trnH-psbA) and from Ziegenhagen et al. (2005) (nad5-4). Polymerase chain reactions (PCR) were performed as follows: denaturation at 95°C for 5 min, followed by 35 cycles at 94°C for 30 sec, 48°C or 53°C or 58°C for 30 sec (matK, or rbcL and trnH-psbA, or nad5-4) and 72°C for 45 sec with a final 10 min extension step at 72°C. PCR final volume was optimized to 30 μl. The PCR mix contained: 0.2 mM of each dNTP, 2.5 mM of MgCl2, 0.3 μM of each primers, 1X GoTaq® Flexi Buffer, 1.25 U of GoTaq® DNA Polymerase (Promega, USA) and 2 μl of 15 ng⁄μl DNA. PCR products were quality-checked on a 1.5% agarose gel stained with Ethidium Bromide (EtBr) and successfully amplified samples were Sanger sequenced. Electrophoregrams were visually inspected, low quality sequences were trimmed and insertions and deletions were excluded from the analysis. The consensus sequence of each organelle DNA region was obtained using CodonCode Aligner 3.7.1.

Kress, W.J., Erickson D.L. (2007). A two-locus global DNA barcode for land plants: the coding rbcL gene complements the non-coding trnHpsbA spacer region. PLOS ONE 2(6): e508.

Kress, W.J., Wurdack, K.J., Zimmer E.A., Weigt L.A., Janzen, D.H. (2005). Use of DNA barcodes to identify flowering plants. PNAS 102(23): 8369–8374.

Ziegenhagen B., Fady B., Kuhlenkamp V., Liepelt S., 2005. Differentiating groups of Abies species with a simple molecular marker. Silvae Genetica, 54(3), 123-126.

**Table S1.5:** Frequency of null alleles within population at 14 nSSR loci

| Population / nSSR locus | PHA_4783 | PHA_6062 | pn1403 | pn2153 | pn2246 | pn4379 | pn6175 | pn6266 | pn6360 | pn7754 | pn8747 | PtTX3107 | PtTX4001 | SPAG_7.14 |
| --- | --- | --- | --- | --- | --- | --- | --- | --- | --- | --- | --- | --- | --- | --- |
| DZA-01 | 0.525 | 0.923 | 0.579 | 0.253 | 0.080 | 0.320 | 0.329 | 0.773 | 0.548 | 0.785 | 0.828 | 0.319 | 0.775 | 0 |
| MAR-01 | 0.069 | 0.016 | 0.712 | 0.064 | 0 | 0.028 | 0.185 | 0.952 | 0.067 | 0.186 | 0.410 | 0.593 | 0.451 | 0 |
| ESP-01 | 0.344 | 0.110 | 0.002 | 0.056 | 0.152 | 0.320 | 0.044 | 0.565 | 0.004 | 0.182 | 0.242 | 0.257 | 0.351 | 0 |
| ESP-02 | 0.560 | 0.009 | 0.112 | 0.049 | 0.000 | 0 | 0.001 | 0.690 | 0.180 | 0.066 | 0.056 | 0.752 | 0.049 | 0.004 |
| FRA-01 | 0.197 | 0.142 | 0.119 | 0.128 | 0.099 | 0.298 | 0.524 | 0.026 | 0.128 | 0.629 | 0.001 | 0.830 | 0.469 | 0 |
| FRA-02 | 0.563 | 0.422 | 0.003 | 0.150 | 0.007 | 0.018 | 0.388 | 0.786 | 0.973 | 0.420 | 0.378 | 0.313 | 0.797 | 0 |
| ITA-01 | 0.129 | 0.174 | 0.107 | 0.217 | 0.010 | 0.697 | 0.277 | 0.279 | 0.289 | 0.341 | 0.803 | 0.048 | 0.011 | 0.006 |
| ITA-02 | 0.254 | 0.917 | 0.305 | 0 | 0.297 | 0.012 | 0 | 0.944 | 0.295 | 0.187 | 0.679 | 0.561 | 0.023 | 0.066 |
| AUT-01 | 0.561 | 0.569 | 0.760 | 0 | 0 | 0.183 | 0.003 | 0.233 | 0.567 | 0.249 | 0.794 | 0.006 | 0.264 | 0.022 |
| SCG-01 | 0.193 | 0.731 | 0.283 | 0.994 | 0.001 | 0.577 | 0.162 | 0.467 | 0.442 | 0.844 | 0.149 | 0.000 | 0.460 | 0.007 |
| SCG-02 | 0.237 | 0.744 | 0.899 | 0.309 | 0.044 | 0.746 | 0.033 | 0.602 | 0.557 | 0.464 | 0.026 | 0.146 | 0.002 | 0.544 |
| ROU-01 | 0.707 | 0.542 | 0.797 | 0.047 | 0.222 | 0.966 | 0.191 | 0.705 | 0.783 | 0.534 | 0.074 | 0.015 | 0.166 | 0.919 |
| HRV-01 | 0.033 | 0.938 | 0.397 | 0.087 | 0.321 | 0.356 | 0.166 | 0.223 | 0.042 | 0.272 | 0.503 | 0.032 | 0.309 | 0.034 |
| HRV-02 | 0.526 | 0.454 | 0.687 | 0.071 | 0.041 | 0.004 | 0.114 | 0.018 | 0.045 | 0.305 | 0.057 | 0.176 | 0.488 | 0.069 |
| CRIMEA-01 | 0.529 | 0.376 | 0.438 | 0.030 | 0.193 | 0.014 | 0.001 | 0.244 | 0.270 | 0.051 | 0.384 | 0.000 | 0.392 | 0.269 |
| CRIMEA-02 | 0.530 | 0.504 | 0.018 | 0.107 | 0.039 | 0.349 | 0.115 | 0.242 | 0.052 | 0.040 | 0.114 | 0.052 | 0.051 | 0.001 |
| TUR-01 | 0.032 | 0.099 | 0.719 | 0.259 | 0.348 | 0.358 | 0.631 | 0.561 | 0.419 | 0.413 | 0.001 | 0.030 | 0.045 | 0 |
| TUR-02 | 0.829 | 0.235 | 0.636 | 0.526 | 0.001 | 0.439 | 0.048 | 0.245 | 0.267 | 0.606 | 0.565 | 0.044 | 0.006 | 0.545 |
| CYP-01 | 0.031 | 0.956 | 0.664 | 0.119 | 0.011 | 0.156 | 0.051 | 0.020 | 0.217 | 0.577 | 0.496 | 0.000 | 0.408 | 0.013 |

**Table S1.6:** Within population haplotypic diversity at nuclear genes. Values were averaged over the eighteen populations

| **Gene** | **Size** | **P** | **H** | **Hd** | **pi** |
| --- | --- | --- | --- | --- | --- |
| 2_1405_01 | 323 | 1 | 2 | 0.246 | 0.0008 |
| 0_2078_01 | 326 | 6 | 7 | 0.521 | 0.0020 |
| 0_6293_01 | 343 | 4 | 7 | 0.444 | 0.0016 |
| 0_7916_01 | 317 | 9 | 21 | 0.723 | 0.0052 |
| 0_8479_01 | 432 | 8 | 15 | 0.758 | 0.0039 |
| 0_10162_01 | 329 | 6 | 12 | 0.588 | 0.0025 |
| 0_10384_02 | 437 | 8 | 10 | 0.571 | 0.0057 |
| 0_10667_02 | 242 | 10 | 28 | 0.888 | 0.0107 |
| 0_13484_01 | 431 | 7 | 6 | 0.558 | 0.0026 |
| 0_13957_02 | 303 | 5 | 7 | 0.62 | 0.0037 |
| 0_14221_01 | 468 | 11 | 29 | 0.805 | 0.0057 |
| 0_16810_02 | 247 | 10 | 14 | 0.61 | 0.0122 |
| 0_18101_02 | 539 | 14 | 40 | 0.848 | 0.0055 |
| CL4470Ct1 | 468 | 10 | 13 | 0.263 | 0.0009 |

Size: length in base pair of the sequence, P: number of polymorphic sites, H: number of haplotypes, Hd: Haplotype Diversity, pi: Nucleotide Diversity

**Table S1.7:** Within population genetic diversity at organelle gene, cpSSRs, nSSRs and nuclear genes

|  | Organelle gene | cpSSR | cpSSR | nSSR | nSSR | nSSR | nSSR | Nuclear genes | Nuclear genes | Nuclear genes | Nuclear genes |
| --- | --- | --- | --- | --- | --- | --- | --- | --- | --- | --- | --- |
| Population | He | He | Na | F | He | Ho | Na | He | Ho | Na | LD* |
| DZA-01 | 0 | 0.389 | 2.75 | -0.15 | 0.693 | 0.797 | 6.429 | 0.535 | 0.369 | 4.071 | 9.429 |
| MAR-01 | 0 | 0.49 | 2.25 | -0.052 | 0.64 | 0.667 | 5.214 | 0.619 | 0.454 | 4.786 | 7.000 |
| ESP-01 | 0 | 0.59 | 3.75 | -0.069 | 0.67 | 0.67 | 7.571 | 0.63 | 0.528 | 4.786 | 5.429 |
| FRA-01 | 0 | 0.681 | 4.25 | -0.008 | 0.697 | 0.692 | 6.429 | 0.619 | 0.363 | 4.571 | 6.143 |
| AUT-01 | 0 | 0.556 | 3.25 | 0.004 | 0.692 | 0.685 | 6.929 | 0.744 | 0.624 | 6.071 | 9.143 |
| ITA-01 | 0 | 0.688 | 4.75 | 0.036 | 0.738 | 0.707 | 8.357 | 0.563 | 0.373 | 4.643 | 5.286 |
| ITA-02 | 0 | 0.601 | 3.75 | 0.021 | 0.753 | 0.733 | 9.071 | 0.651 | 0.503 | 5.143 | 9.286 |
| FRA-02 | 0 | 0.632 | 3.75 | -0.082 | 0.692 | 0.74 | 6.571 | 0.583 | 0.391 | 4.714 | 5.429 |
| HRV-01 | 0 | 0.502 | 2.75 | 0.058 | 0.705 | 0.677 | 6.429 | 0.668 | 0.476 | 5.5 | 12.500 |
| HRV-02 | 0 | 0.611 | 3.75 | 0.02 | 0.721 | 0.707 | 7.143 | 0.664 | 0.509 | 5.286 | 7.214 |
| SCG-02 | 0 | 0.681 | 4.25 | -0.032 | 0.722 | 0.74 | 7.786 | 0.745 | 0.603 | 6.214 | 8.286 |
| SCG-01 | 0 | 0.576 | 3.5 | -0.03 | 0.717 | 0.732 | 8.214 | 0.702 | 0.452 | 5.429 | 5.357 |
| ROU-01 | 0 | 0.653 | 4.25 | -0.038 | 0.662 | 0.708 | 7.429 | 0.669 | 0.56 | 5.571 | 12.571 |
| CRIMEA-01 | 0 | 0.552 | 3.5 | 0.035 | 0.716 | 0.685 | 7.357 | 0.705 | 0.515 | 5.429 | 11.714 |
| TUR-01 | 0 | 0.667 | 4.25 | 0.111 | 0.723 | 0.694 | 8.143 | 0.66 | 0.377 | 5.071 | 9.500 |
| TUR-02 | 0 | 0.528 | 3.75 | -0.016 | 0.7 | 0.706 | 8.143 | 0.679 | 0.537 | 5.786 | 10.643 |
| CYP-01 | 0 | 0.649 | 4 | 0.024 | 0.718 | 0.703 | 7.429 | 0.666 | 0.329 | 4.929 | 8.643 |
| CRIMEA-02 | 0 | 0.537 | 3.25 | 0.046 | 0.705 | 0.655 | 7.143 | NA | NA | NA | NA |

LD*: number of pairs of SNPs within gene (averaged over a total of 14 genes) displaying a significant linkage disequilibrium within population

**Figure S1.1:** Linear regression between genetic diversity and longitude of black pine populations. Diversity is measured by (a) expected heterozygosity (*He*) at nuclear genes (b) *He* at nSSR.

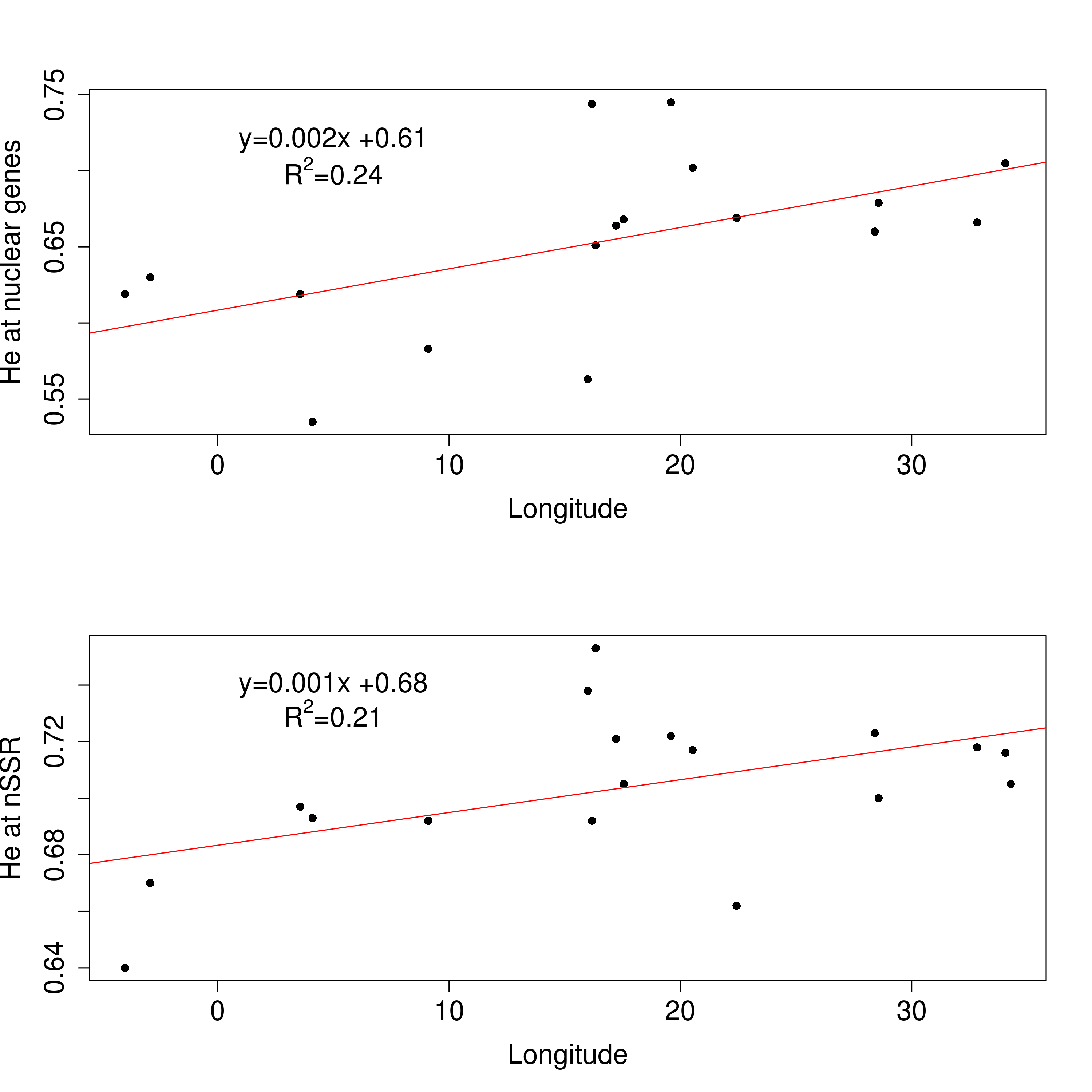

**Appendix S2:** details of ABC analyses

**Figure S2.1:** Alternative scenario of divergence

**Figure S2.2:** Scenarios of admixture

**Figure S2.3:** Evaluation of the simulated datasets to test for the scenarios of divergence: Principal component analysis

**Figure S2.4:** Posterior probability of scenarios: logistic regression.

**Figure S2.5:** Model checking for the scenario of divergence

**Figure S2.6:** comparison of the best scenario of divergence with an alternative scenario allowing demographic changes along the branches

**Figure S2.7:** Evaluation of the simulated datasets to infer admixture parameters: Principal component analysis

**Figure S2.8:** Model checking for the scenarios of admixture

**Table S2.1:** Prior distribution of the mutation parameters used in DIYABC and detailed description of the divergence and admixture demographic scenarios.

**Table S2.2:** Evaluation of the simulated data set to test for the scenarios of divergence: test of rank.

**Table S2.3:** Evaluation of the simulated data set to test for the scenarios of divergence: Posterior error rate.

**Table S2.4:** Estimates of the ratio N_current_/N_past_

**Table S2.5:** Accuracy test of parameters estimation for the scenarios of divergence

**Table S2.6:** Model checking for the best scenario of divergence

**Table S2.7:** Evaluation of the simulated datasets to infer admixture parameters: test of rank.

**Table S2.8:** Inference of demographic parameters in the scenarios with admixture

**Table S2.9:** Accuracy test of parameters estimation for the scenarios of admixture

**Table S2.10:** Model checking for the scenarios of admixture

**Figure S2.1:** Alternative scenario of divergence

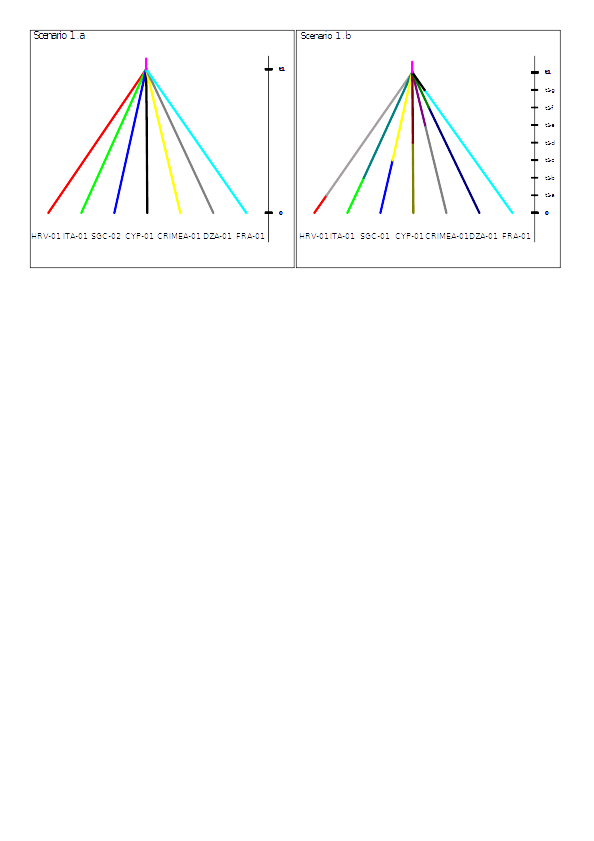

Scenario 1.a corresponds to Scenario 1 which is the best scenario of divergence (among six tested) without taking into account demographic changes along the branches. Scenario 1.b corresponds to scenario 1 when each of the seven populations could have had their effective population resized at independent times.

**Figure S2.2:** Scenarios of admixture - Five ABC analyses were performed to infer past events of migration between five pairs of populations

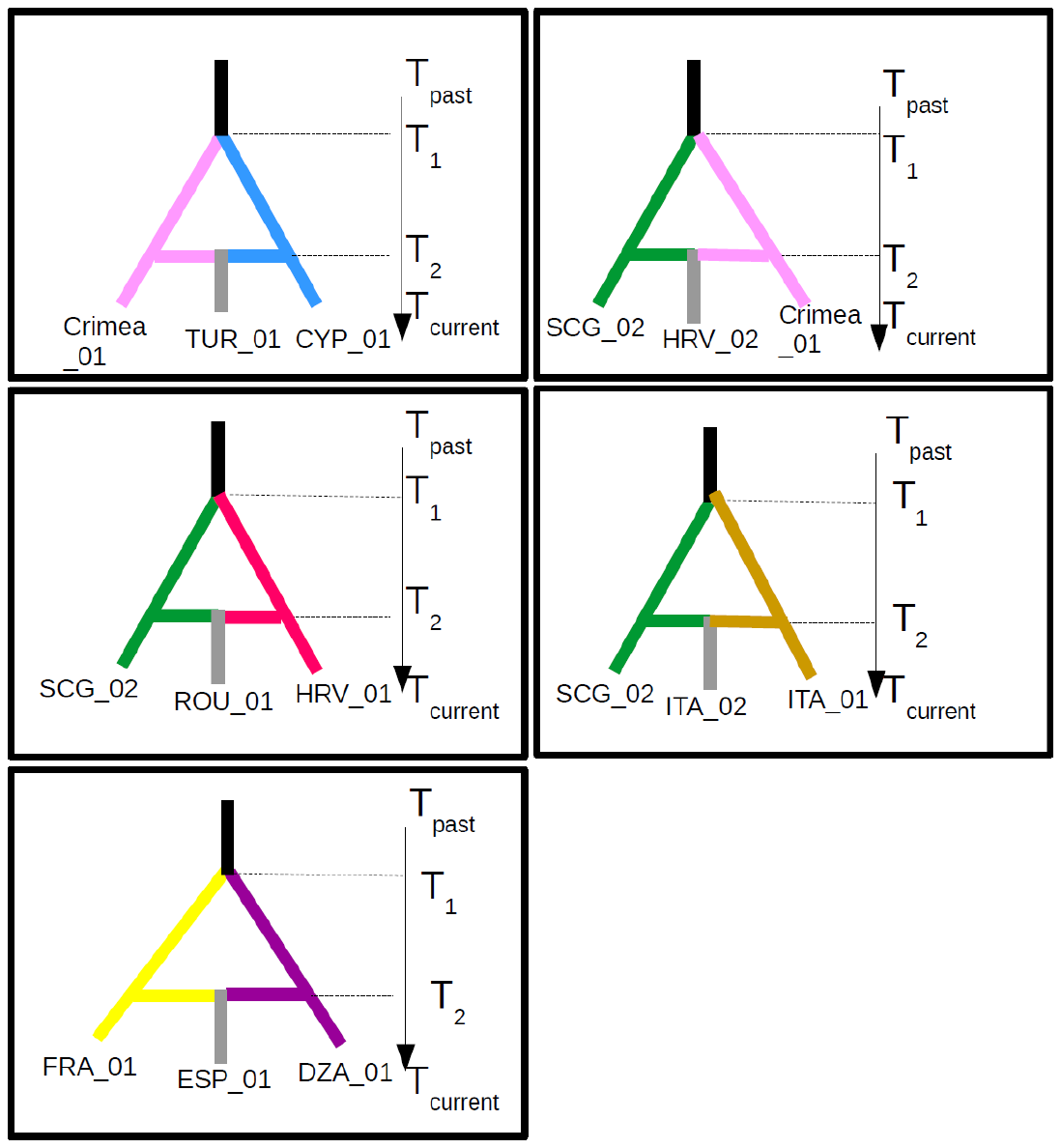

**Figure S2.3:** Evaluation of the simulated datasets to test for the scenarios of divergence: Principal component analysis

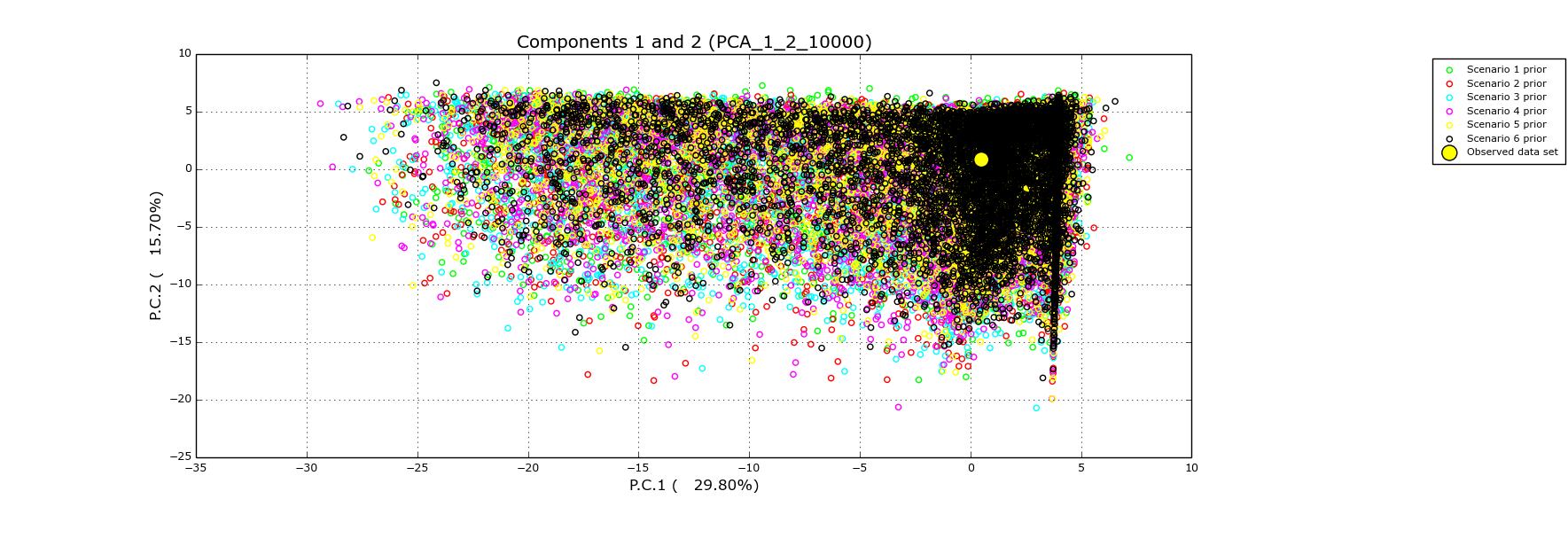

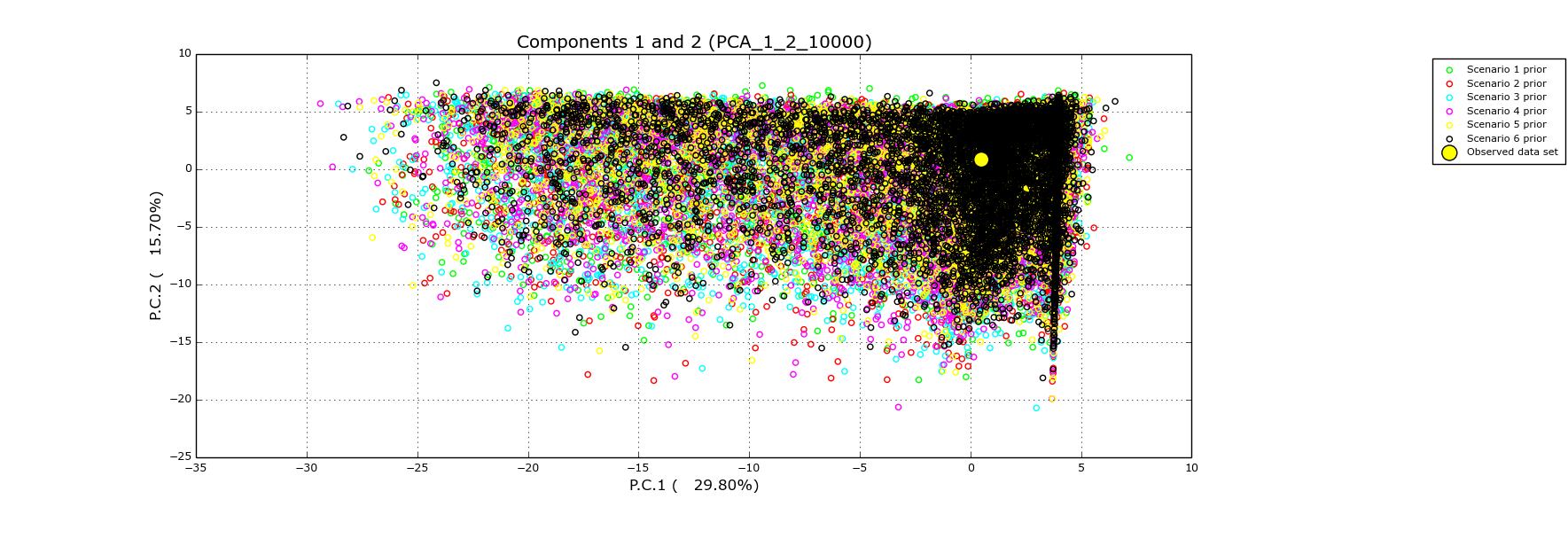

The purpose is to check that at least one combination of scenarios and priors can produce simulated data sets that are close enough to the observed data set. Principal component analysis is performed in the space of summary statistics on at most 100,000 simulated data set and the observed data is added on each plane (biggest yellow point) of the analysis in order to evaluate how the latter is surrounded by simulated data sets (6 colours for 6 scenario of divergence).

**Figure S2.4:** Posterior probability of scenarios of divergence: logistic regression.

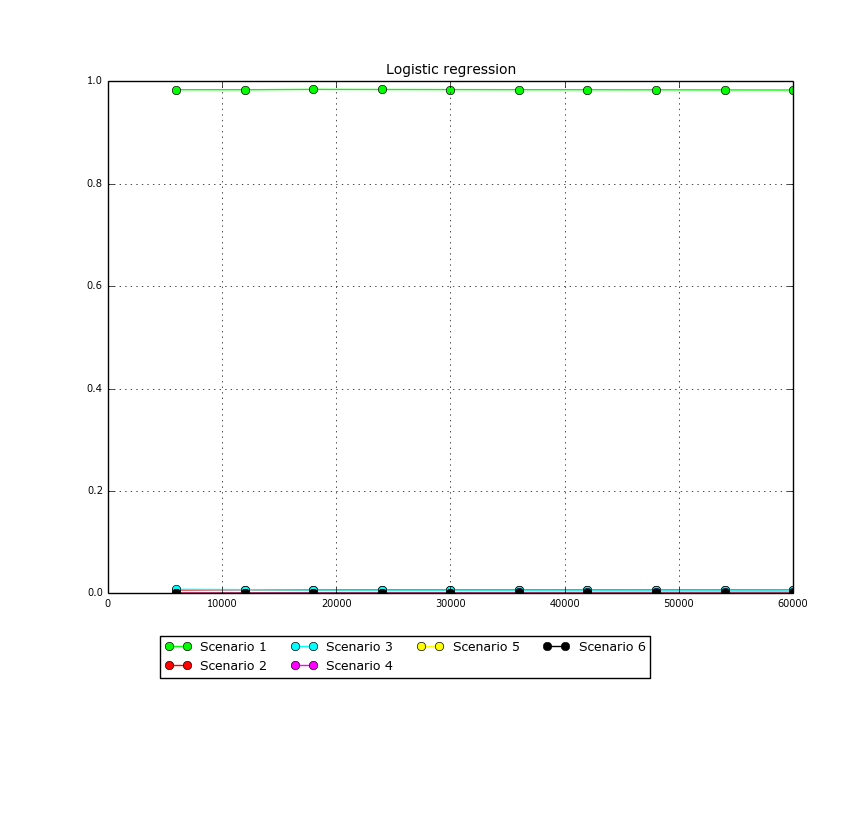

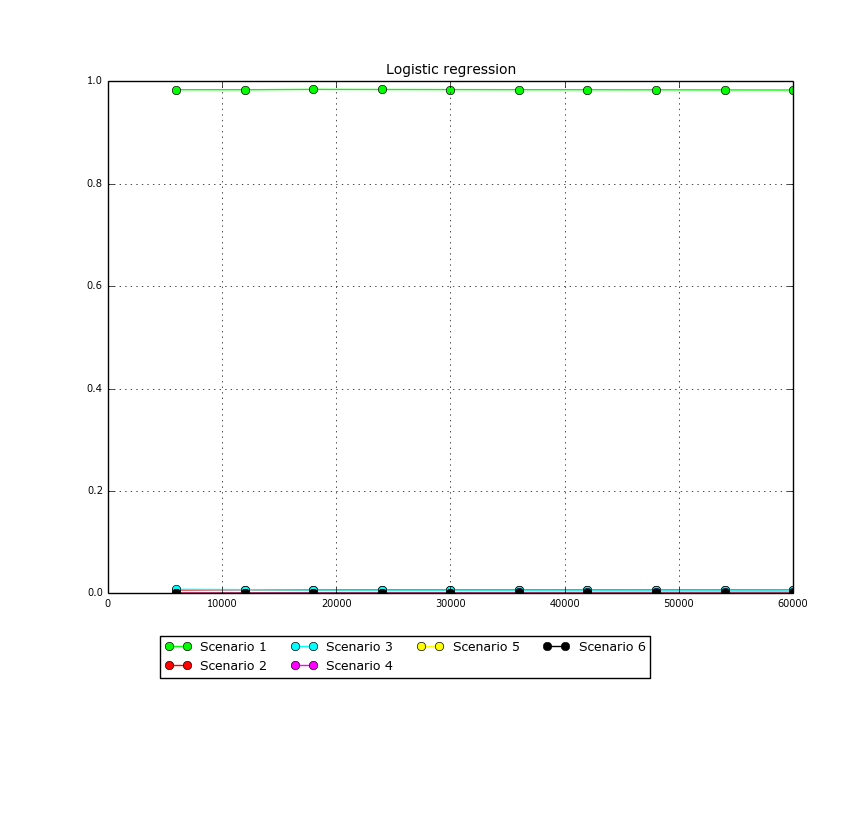

A polychotomic weighted logistic regression is performed on the first 10% data sets with the proportion of the scenario as the dependent variable and the differences between observed and simulated data set summary statistics as the independent variables. The intercept of the regression (corresponding to an identity between simulated and observed summary statistics) is taken as the point estimate. (Cornuet et al., 2008). Seven scenarios are compared

**Figure S2.5:** Model checking for the best scenario of divergence (scenario 1).

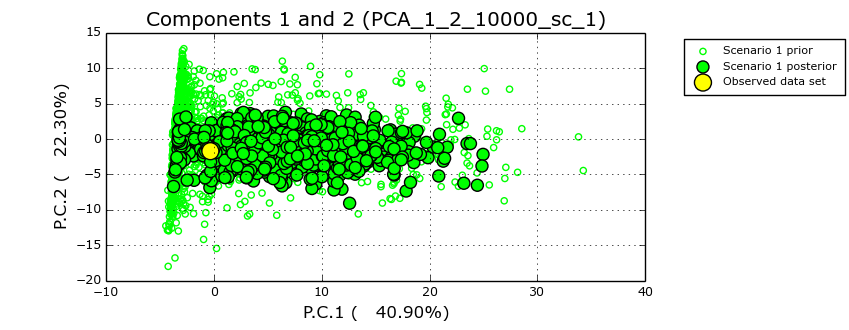

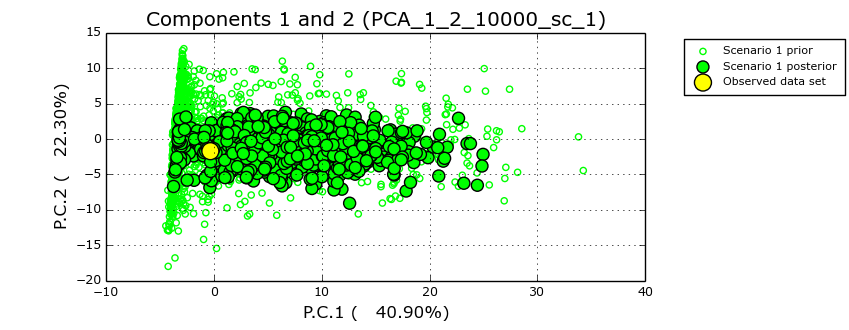

**Figure S2.6:** Comparison of the best scenario of divergence (S1a) with a scenario allowing demographic changes along the branches (S1b)

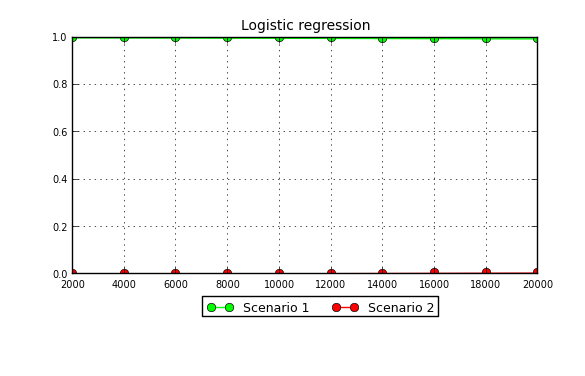

A polychotomic weighted logistic regression is performed on the first 10% data sets with the proportion of the scenario as the dependent variable and the differences between observed and simulated data set summary statistics as the independent variables. The intercept of the regression (corresponding to an identity between simulated and observed summary statistics) is taken as the point estimate. (Cornuet et al., 2008). Scenario 1 corresponds to the best scenario of divergence without demographic changes along the branches whereas scenario 2 allows demographic changes.

**Figure S2.7:** Evaluation of the simulated datasets to infer admixture parameters: Principal component analysis

The purpose of this analysis is to check that at least one combination of scenarios and priors can produce simulated data (green points) sets that are close enough to the observed data set (yellow point). A Principal Component Analysis is performed in the space of summary statistics on at most 100,000 simulated data sets and the observed data are added to each plane (large yellow dot) of the analysis in order to evaluate how the latter is surrounded by simulated data sets.

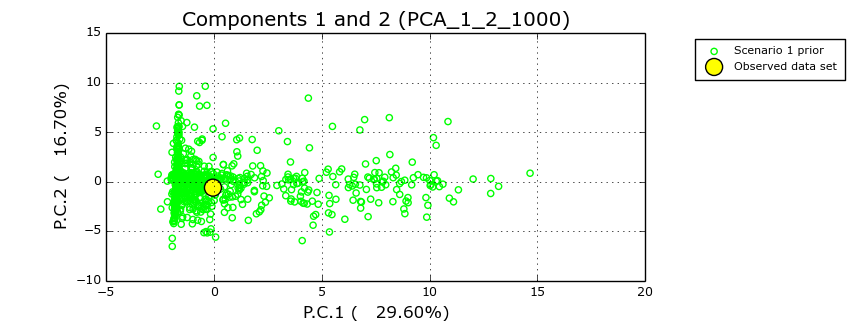

**a.** Population 1 of Cyprus (CYP-01, *P.n. pallasiana)* and population 1 of Crimea *(CRIMEA-01, P.n. pallasiana)* came into contact to give rise to population 1 of Turkey *(TUR-01, P.n. pallasiana).*

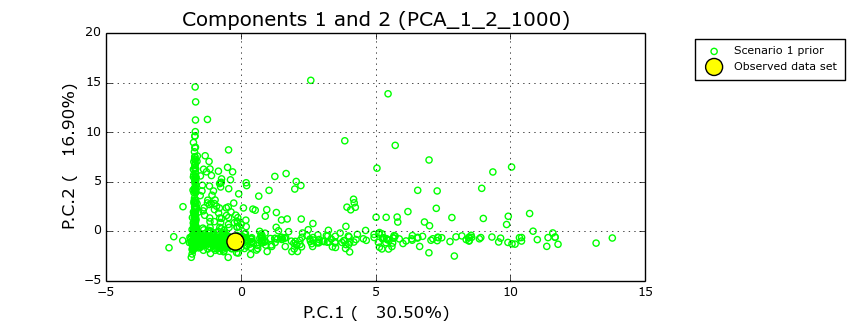

***b****.* Population 1 of France (FRA-01, *P.n. salzmannii)* and population 1 of Algeria *(DZA-01, P.n. salzmannii))* came into contact to give rise to population 1 of Spain *(ESP-01, P.n. salzmannii)*

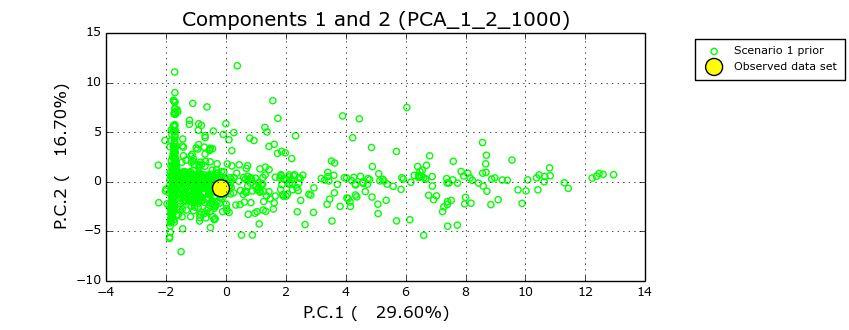

***c.*** Population 2 of Serbia (SCG-02, *P.n. nigra)* and population 1 of Croatia *(HRV-01, P.n. dalmatica)* came into contact to give rise to population 1 of Romania *(ROU-01, P.n. nigra)*

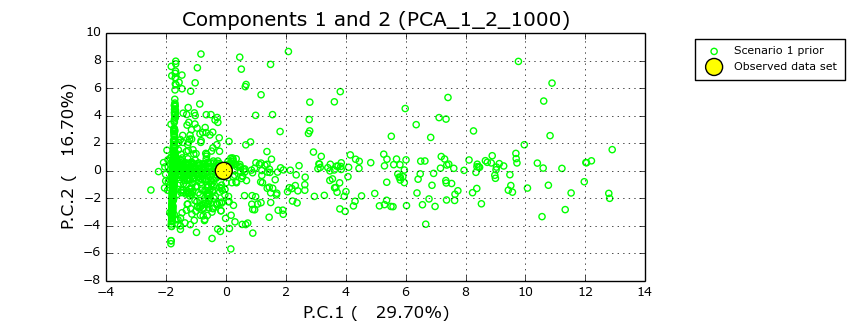

***d****:* Population 2 of Serbia ( SCG-02, *P.n. nigra)* and population 1 of Crimea *(CRIMEA-01, P.n. pallasiana)* came into contact to give rise to population 2 of Croatia *(HRV-02, P.n. dalmatica)*

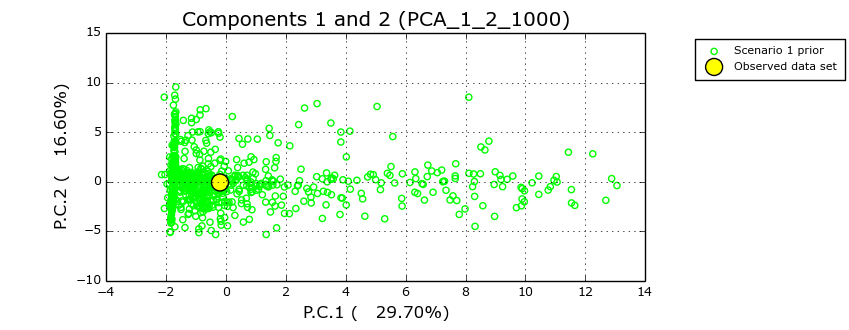

***e****.* Population 1 of Italy (ITA-01, *P.n. laricio)* and population 2 of Serbia *(SCG-02, P.n. pallasiana)* came into contact to give rise to population 2 of Italy *(ITA-02, P.n. laricio)*

**Figure S2.8:** Model checking for the scenarios of admixture

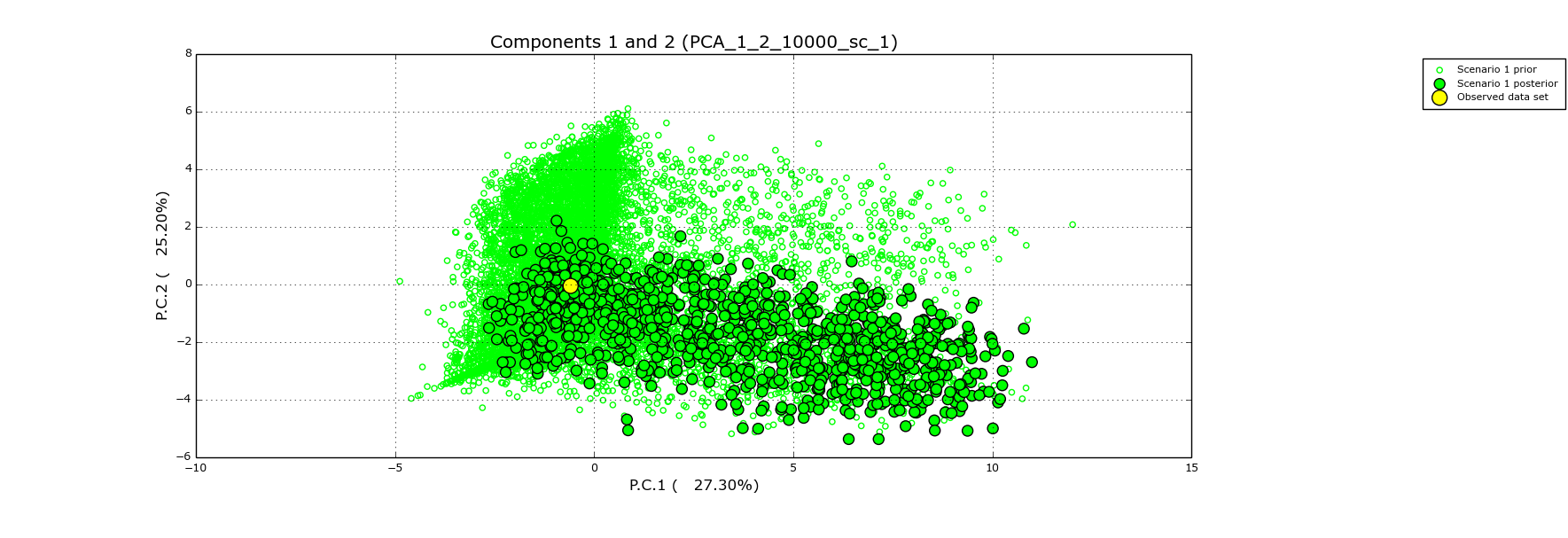

**a**: Population 1 of Cyprus (CYP-01, *P.n. pallasiana*) and population 1 of Crimea (CRIMEA-01, *P.n. pallasiana*) came into contact to give rise to population 1 of Turkey (TUR-01, *P.n. pallasiana*)

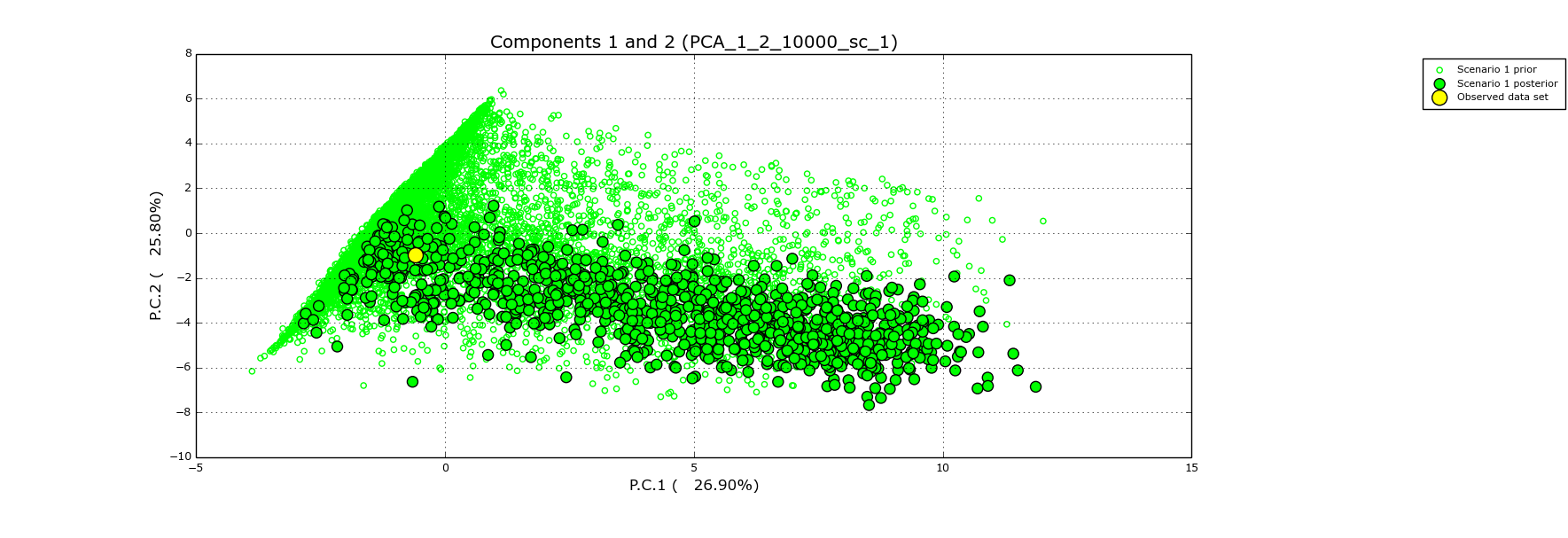

**b**: Population 1 of France (FRA-01, *P.n. salzmannii*) and population 1 of Algeria (DZA-01, *P.n. salzmannii*) came into contact to give rise to population 1 of Spain (ESP-01, *P.n. salzmannii*)

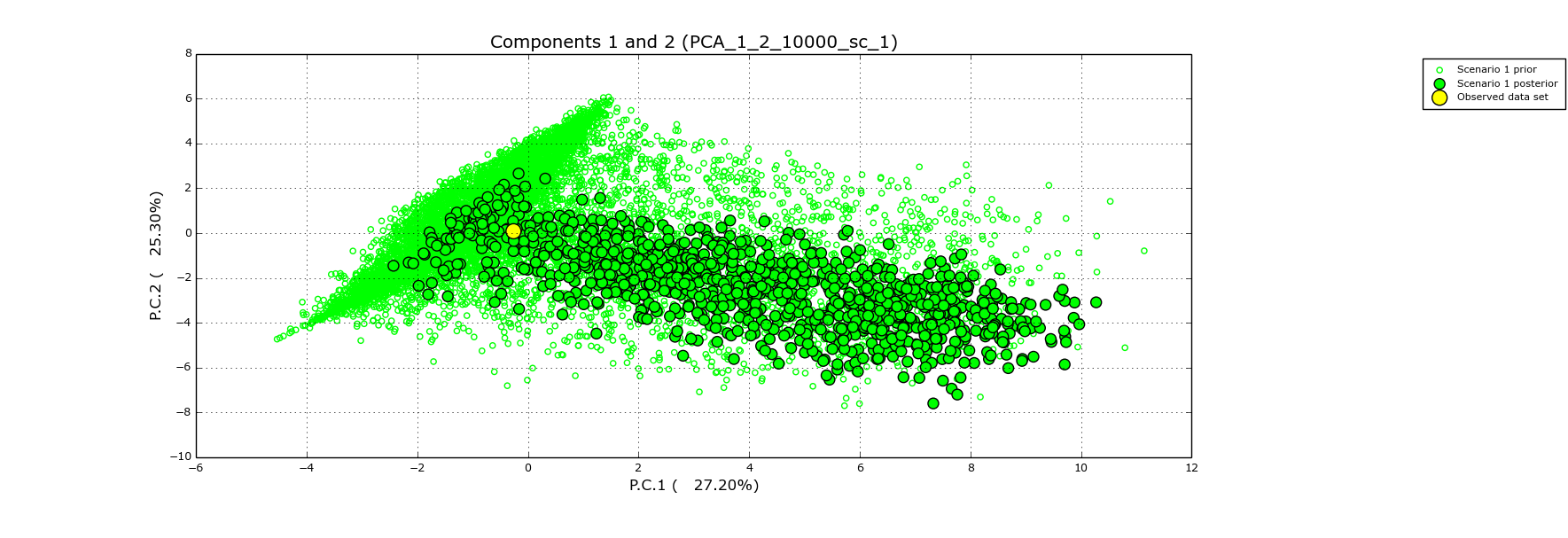

**c**: Population 2 of Serbia (SCG-02, *P.n. nigra*) and population 1 of Croatia (HRV-01, *P.n. dalmatica*) came into contact to give rise to population 1 of Romania (ROU-01, *P.n. nigra*).

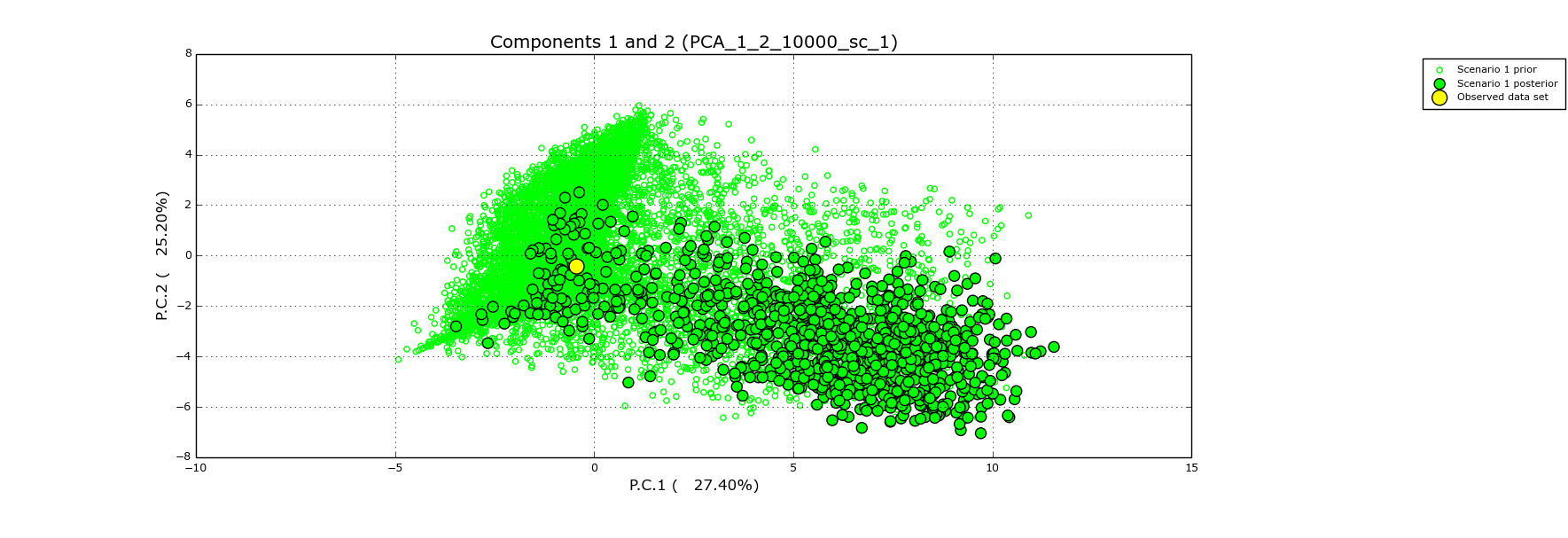

**d**: Population 2 of Serbia (SCG-02, P.n. nigra) and population 1 of Crimea (CRIMEA-01, *P.n. pallasiana*) came into contact to give rise to population 2 of Croatia (HRV-02, *P.n. dalmatic*a).

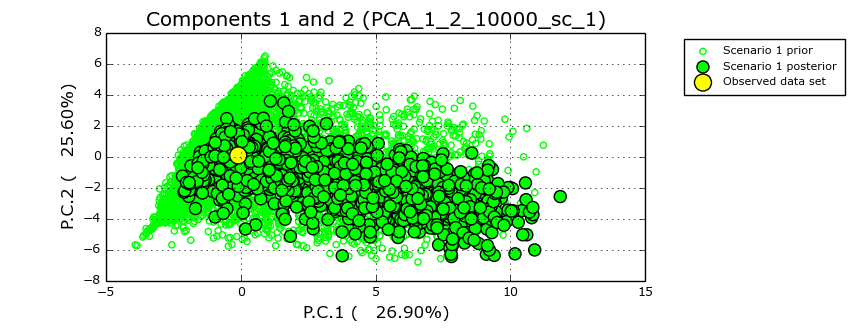

**e**: Population 1 of Italy (ITA-01, *P.n. laricio*) and population 2 of Serbia (SCG-02, *P.n. pallasiana*) came into contact to give rise to population 2 of Italiy(ITA-02, *P.n. laricio*).

**Table S2.1**: Prior distribution of the mutation parameters used in DIYABC and detailed description of the divergence and admixture demographic scenarios.

| **Parameter** | **Distribution** | **Min-Max** | **Mean** | **Shape** |
| --- | --- | --- | --- | --- |
| **nSSR** | | | | |
| Mean mutation rate | Log uniform | 1.10^-4^-1.10^-2^ |  |  |
| Individual mutation rate | gamma | 1.10^-5^-1.10^-2^ | Mean mutation rate | 2 |
| Mean coefficient P | uniform | 0.1-1 |  |  |
| Individual locus coefficient P | Gamma | 0.1-1 | Mean coefficient P | 2 |
| **cpSSR** | | | | |
| Mean mutation rate | Log uniform | 1.10^-6^-1.10^-2^ |  |  |
| Individual mutation rate | gamma | 1.10^-6^-1.10^-2^ | Mean mutation rate | 2 |
| Mean coefficient P | uniform | 0.1-1 |  |  |
| Individual locus coefficient P | Gamma | 0.1-1 | Mean coefficient P | 2 |
| **Nuclear Genes** | | | | |
| Mean mutation rate (per site per generation) | Log uniform | 1.10^-12^-1.10^-5^ |  |  |
| Individual locus mutation rate (Gamma distribution around mean) | Gamma | 1.10^-12^-1.10^-5^ | Mean mutation rate | 2 |
| Mean coefficient k_C/T | Log uniform | 0.05-20 |  |  |
| Individual locus coefficient k_C/T (Gamma distribution around mean) | Gamma | 0.05-20 | Mean coefficient k_C/T | 2 |
| Mutation model | Hasegawa-Kishino-Yano | | | |
| Percentage of invariant sites | Zero | | | |
| **Organelle Genes** | | | | |
| Mean mutation rate (per site per generation) | Log uniform | 1.10^-12^-1.10^-5^ |  |  |
| Individual locus mutation rate (Gamma distribution around mean) | Gamma | 1.10^-12^-1.10^-5^ | Mean mutation rate | 2 |
| Mean coefficient k_C/T | Log uniform | 0.05-20 |  |  |
| Individual locus coefficient k_C/T (Gamma distribution around mean) | Gamma | 0.05-20 | Mean coefficient k_C/T | 2 |
| Mutation model | Hasegawa-Kishino-Yano | | | |
| Percentage of invariant sites | Zero | | | |

a) Divergence scenarios

Given the large number of possible scenarios when seven genetic lineages are present, we reduced our investigation by focusing on how populations were grouped after the STRUCTURE analysis. The most likely number of clusters using nSSRs, cpSSRs and genes was seven. We thus choose, as the simplest scenario, a rake shape tree (Scenario 1) with seven branches. For all other scenarios (Figure 2), tree shapes and branch grouping to obtain the seven tips were chosen according to the grouping of the populations resulting from STRUCTURE analyses based on separate or partial combinations of markers and the literature: scenario 2 (nSSR only, best K=6), scenario 3 (following the taxonomic nomenclature of Table 1, K=5), scenario 4 (nSSR and the most geographically discriminant cpSSR, best K=3), scenarios 5 and 6 (nuclear genes only, best K=2). The results of the STRUCTURE analysis using only nuclear genes puts populations considered as *Pinus nigra laricio* in Table 1 as intermediate between a western and an eastern cluster, hence the two alternative scenarios 5 and 6.

Because population subdivision can leave genetic patterns that could be wrongly attributed to a bottleneck effect (Peter et al., 2010), the analysis was performed in two steps. First without allowing for effective population expansion or contraction. Once the best scenario of divergence identified, we allowed effective population size changes along the branches to check whether black pine populations had undergone size changes after lineage divergence.

Demographic prior parameters were drawn using a truncated normal distribution for both population effective sizes and the timing of demographic event. Values ranged between a minimum of 10 and a maximum of 10,000 with a mean of 5000 and a standard deviation of 2500. In total, 33 loci were used, 14 nSSRs, 1 cpSSR (pt1254, the only significantly differentiated cpSSR where 33% of the total variation was among populations), 14 nuclear genes and 4 organelle genes.

Coalescence simulations of nSSRs and cpSSRs were done using a generalized stepwise mutation model (GSM; Zhivotovsky et al., 1997; Estoup et al., 2002). For both nuclear and organelle genes, we used the DNA evolution sequence model of Hasegawa-Kishino-Yano (Hasegawa et al., 1985). All prior distributions are given in the table above.

We applied the procedure “evaluate scenario-prior combination” (Cornuet et al., 2010) which is based on both a principal component analysis and a test of rank, to choose among the available summary statistics that are available (714 in total), a subset that did not under- or over-estimate genetic distance (in simulated data sets compared to the observed one). A total of 124 statistics were chosen.

The 124 summary statistics were as follow:

- For nSSRs: mean genic diversity within population and *F_ST_* between pairs of populations.
- For cpSSR: mean number of alleles within population and *F_ST_* between five pairs of populations (two pairs: ITA_1/FRA_1 and SCG_2/CRIMEA_1 were excluded because *F_ST_* values were equal to zero).
- For nuclear and organelle genes: number of haplotypes within population and mean of pairwise differences within populations. For nuclear genes: number of segregating sites between pairs of populations and *F_ST_* between pairs of populations.

For each scenario, one million simulations were performed generating a total of six million datasets for the six tested scenarios. The most likely scenario was selected according to the results of the logistic regression analysis and the posterior distribution of demographic parameters was estimated for the best scenario. Bias and precision of parameters estimation were also computed (Mean relative bias (MRB), relative root mean square (RMSE), 50% and 95% coverage and Factor 2). Finally, the best scenario was validated by carrying out an additional analysis using 1,000,000 simulated datasets and eight summary statistics that were not used during the simulation procedure.

The eight summary statistics were:

- For nSSRs: mean number of alleles within population and mean genic diversity between pairs of populations.
- For cpSSR, mean genetic diversity within populations and mean number of alleles between pairs of populations.
- For nuclear genes, mean number of segregating sites within population, mean number of haplotypes between pairs of populations, mean of pairwise differences between pairs of populations.
- For organelle genes, mean number of segregating sites within population.

Once the best scenario of divergence was identified, we compared its probability to an alternative scenario in which each of the seven populations could have had their effective population resized at independent times (Figure S2.1 in Appendix S2). Parameters for the mutation model and the summary statistics selection were the same as described above for the previous demographic inference analysis. A total of two million datasets was produced and the same post analyses as before were performed. All scenario analyses were performed using the R environment (R Core Team, 2015).

b) Admixture scenarios

To minimize the complexity of possible admixture scenarios, we selected populations with a high membership to a maximum of two of parental lineages inferred from the STRUCTURE analysis (scenarios 1, 2 and 3 below). We extended the approach to two other cases where parental lineage contribution was less than 90%, to explore admixture events in other geographical areas not covered by the first three scenarios (scenarios 4 and 5 below).

From the west to the east:

**(1)** FRA-01 (*P.n. salzmannii*, France) + DZA-01 (*P. n. salzmannii*, Algeria) = ESP-01 (*P. n. salzmannii*, Spain);

**(2)** ITA-01 (*P. n. laricio*, Italy) + SCG-02 (*P.n. nigra*, Serbia) = ITA-02 (*P. n. laricio*, Italy);

**(3)** SCG-02 (*P.n. nigra*, Serbia) + HRV-01 (*P. n. dalmatica,* Croatia*) =* ROU-01 *(P.n. nigra,* Romania);

**(4)** SCG-02 (*P.n. nigra*, Serbia) **+** CRIMEA-01 (*P.n. pallasiana*, Crimea) = HRV-02 (*P. n. dalmatica,* Croatia*)*;

**(5)** CYP-01 (*P. n. pallasiana*, Cyprus) + CRIMEA-01 (*P.n. pallasiana*, Crimea) = TUR-01 (*P.n. pallasiana*, Turkey).

The resulting five ABC analyses performed to infer past events of migration between the five pairs of populations are shown in Figure S2.2 of Appendix S2. The priors used were the same as the ones used to test the divergent scenario with, in addition, the rate of admixture drawn in a normal distribution truncated between 0.01 and 0.99, of average 0.5 and standard deviation 0.25. The same categories of summary statistics (SS) were used giving rise to 30 SS in total for the admixture scenarios. Bias and precision of parameter estimations and model checking were done using the same procedures as for divergence scenarios.

Cornuet, J.M., Ravigné, V., Estoup, A. (2010). Inference on population history and model checking using DNA sequence and microsatellite data with the software DIYABC (v1.0). BMC Bioinformatics 11: 401.

Estoup, A., Jarne, P. & Cornuet, J.M. (2002). Homoplasy and mutation model at microsatellite loci and their consequences for population genetics analysis. Molecular Ecology 11: 1591–1604.

Hasegawa, M., Kishino, H., & Yano, T.A. (1985). Dating of the human-ape splitting by a molecular clock of mitochondrial DNA. Journal of molecular evolution 22: 160-174.

Peter, B.M., Wegmann, D. & Excoffier, L. (2010). Distinguishing between population bottleneck and population subdivision by a Bayesian model choice procedure. Molecular Ecology 19: 4648–4660.

R Core Team, (2015) R: A language and environment for statistical computing. R Foundation for Statistical Computing, Vienna, Austria. URL <https://www.R-project.org/>.

Zhivotovsky, L.A., Feldman, M.W. & Grishechkin, S.A. (1997). Biased mutations and microsatellite variation. Molecular Biology and Evolution 14: 926–933.

**Table S2.2:** Evaluation of the simulated data set to test for the scenarios of divergence: test of rank.

Test of rank: Each summary statistic (124 in total) of the observed data set is ranked against those of the simulated data set for each scenario (6 in total). This analysis helps finding which aspects of the model (including prior) have been mistated. For instance, a grossly overestimated genetic distance (in 6000000 simulated data sets compared to the observed one) may suggest a mispecification of the prior distribution of the time of divergence of the two involved populations or of the mean mutation rate of the markers.

The code used for the summary statistics is composed of three parts: the summary statistic (see above for the definitions), the type of marker (1,2,3,4 for nSSR, cpSSR, nuclear gene, organelle gene respectively), the population (1,2,3,4,5,6,7 for HRV_01, ITA_01, SCG_02, CYP_01, UKR_01, DZA_01, CYP_01 respectively)

*HET*:mean genic diversity within population, *FST_:_* genetic differentiation between pairs of populations, *NAL*: mean number of alleles within population, *NHA*: number of haplotypes within population, *MPD*: mean of pairwise differences within populations, *NS*: number of segregating sites between pairs of populations and *HST* genetic differentiation based on sequence haplotypes between pairs of populations.

Values indicate for each summary statistics the proportion of simulated data sets which have a value below the observed one

|  | Summary statistics | Observed value | S1 | S2 | S3 | S4 | S5 | S6 |
| --- | --- | --- | --- | --- | --- | --- | --- | --- |
| 1 | HET_1_1 | 0.7357 | 0.2959 | 0.2939 | 0.2938 | 0.2935 | 0.2931 | 0.2938 |
| 2 | HET_1_2 | 0.77 | 0.3551 | 0.3537 | 0.3541 | 0.3534 | 0.3527 | 0.3603 |
| 3 | HET_1_3 | 0.7566 | 0.3311 | 0.3297 | 0.3362 | 0.3359 | 0.3345 | 0.3356 |
| 4 | HET_1_4 | 0.7496 | 0.3189 | 0.3176 | 0.3173 | 0.3236 | 0.3224 | 0.3241 |
| 5 | HET_1_5 | 0.7469 | 0.3144 | 0.3127 | 0.3123 | 0.3122 | 0.3182 | 0.319 |
| 6 | HET_1_6 | 0.7254 | 0.279 | 0.2771 | 0.2776 | 0.2837 | 0.2828 | 0.2767 |
| 7 | HET_1_7 | 0.742 | 0.3062 | 0.311 | 0.3111 | 0.3104 | 0.3095 | 0.3108 |
| 8 | FST_1_1&2 | 0.0691 | 0.3954 | 0.3546 | 0.3435 | 0.3378 | 0.3438 | 0.4211 |
| 9 | FST_1_1&3 | 0.0454 | 0.237 | 0.2009 | 0.1866 | 0.1822 | 0.2596 | 0.2584 |
| 10 | FST_1_1&4 | 0.0903 | 0.5 | 0.4559 | 0.4428 | 0.5201 | 0.5286 | 0.527 |
| 11 | FST_1_1&5 | 0.0802 | 0.4547 | 0.4119 | 0.3995 | 0.3938 | 0.4826 | 0.4808 |
| 12 | FST_1_1&6 | 0.1561 | 0.6979 | 0.6554 | 0.6403 | 0.6269 | 0.6356 | 0.6407 |
| 13 | FST_1_1&7 | 0.123 | 0.5784 | 0.5316 | 0.5186 | 0.5128 | 0.5197 | 0.5184 |
| 14 | FST_1_2&3 | 0.0537 | 0.2987 | 0.2602 | 0.2463 | 0.241 | 0.2463 | 0.3242 |
| 15 | FST_1_2&4 | 0.1183 | 0.6115 | 0.5658 | 0.5511 | 0.5388 | 0.5468 | 0.6426 |
| 16 | FST_1_2&5 | 0.0816 | 0.4621 | 0.4182 | 0.4061 | 0.3999 | 0.4023 | 0.4905 |
| 17 | FST_1_2&6 | 0.101 | 0.5132 | 0.4717 | 0.4589 | 0.5327 | 0.5415 | 0.455 |
| 18 | FST_1_2&7 | 0.0965 | 0.4719 | 0.4283 | 0.4169 | 0.4909 | 0.4988 | 0.4127 |
| 19 | FST_1_3&4 | 0.0819 | 0.4606 | 0.4171 | 0.401 | 0.3913 | 0.4899 | 0.4887 |
| 20 | FST_1_3&5 | 0.0345 | 0.1469 | 0.1155 | 0.1649 | 0.1594 | 0.1682 | 0.1672 |
| 21 | FST_1_3&6 | 0.1236 | 0.5987 | 0.555 | 0.5358 | 0.5241 | 0.5325 | 0.5361 |
| 22 | FST_1_3&7 | 0.0973 | 0.4734 | 0.4298 | 0.4154 | 0.4095 | 0.4159 | 0.4144 |
| 23 | FST_1_4&5 | 0.0929 | 0.5114 | 0.4669 | 0.4533 | 0.4434 | 0.5424 | 0.5408 |
| 24 | FST_1_4&6 | 0.1351 | 0.6335 | 0.5899 | 0.5752 | 0.5583 | 0.5667 | 0.5705 |
| 25 | FST_1_4&7 | 0.0886 | 0.4325 | 0.3903 | 0.3791 | 0.3703 | 0.3759 | 0.3753 |
| 26 | FST_1_5&6 | 0.1347 | 0.6349 | 0.5913 | 0.5771 | 0.5644 | 0.5675 | 0.5718 |
| 27 | FST_1_5&7 | 0.0994 | 0.485 | 0.4407 | 0.4294 | 0.4238 | 0.4265 | 0.4251 |
| 28 | FST_1_6&7 | 0.1013 | 0.4775 | 0.5177 | 0.5022 | 0.4963 | 0.5037 | 0.5026 |
| 29 | NAL_2_1 | 2 | 0.4991 | 0.4996 | 0.4994 | 0.4989 | 0.4985 | 0.4992 |
| 30 | NAL_2_2 | 5 | 0.771 | 0.7707 | 0.7711 | 0.7714 | 0.7705 | 0.7727 |
| 31 | NAL_2_3 | 3 | 0.6257 | 0.626 | 0.6278 | 0.6277 | 0.6268 | 0.6277 |
| 32 | NAL_2_4 | 3 | 0.6344 | 0.6342 | 0.6346 | 0.6363 | 0.6353 | 0.6369 |
| 33 | NAL_2_5 | 3 | 0.6188 | 0.6188 | 0.6185 | 0.6188 | 0.62 | 0.6205 |
| 34 | NAL_2_6 | 3 | 0.6259 | 0.6257 | 0.6261 | 0.628 | 0.6266 | 0.6264 |
| 35 | NAL_2_7 | 2 | 0.5101 | 0.5123 | 0.5121 | 0.512 | 0.5113 | 0.5126 |
| 36 | FST_2_1&2 | 0.1876 | 0.6573 | 0.628 | 0.6179 | 0.6108 | 0.6167 | 0.6792 |
| 37 | FST_2_1&3 | 0.5707 | 0.8799 | 0.8581 | 0.8478 | 0.8429 | 0.8975 | 0.8972 |
| 38 | FST_2_1&4 | 0.3002 | 0.7502 | 0.7221 | 0.7116 | 0.7681 | 0.772 | 0.7715 |
| 39 | FST_2_1&5 | 0.6288 | 0.8971 | 0.8776 | 0.8692 | 0.8639 | 0.9135 | 0.9136 |
| 40 | FST_2_1&6 | 0.143 | 0.6004 | 0.5722 | 0.5622 | 0.5553 | 0.5612 | 0.5604 |
| 41 | FST_2_1&7 | 0.5319 | 0.8664 | 0.8424 | 0.8329 | 0.8274 | 0.833 | 0.8321 |
| 42 | FST_2_2&3 | 0.2832 | 0.738 | 0.7103 | 0.6987 | 0.6926 | 0.6988 | 0.7595 |
| 43 | FST_2_2&4 | 0.189 | 0.6579 | 0.6288 | 0.6186 | 0.6118 | 0.6178 | 0.6789 |
| 44 | FST_2_2&5 | 0.3569 | 0.7825 | 0.7554 | 0.7454 | 0.7386 | 0.7435 | 0.8029 |
| 45 | FST_2_2&7 | 0.0647 | 0.4757 | 0.4513 | 0.4429 | 0.4917 | 0.4959 | 0.4421 |
| 46 | FST_2_3&4 | 0.397 | 0.8085 | 0.7827 | 0.7708 | 0.7642 | 0.8283 | 0.8277 |
| 47 | FST_2_3&6 | 0.4 | 0.8078 | 0.7818 | 0.7701 | 0.7637 | 0.7702 | 0.7703 |
| 48 | FST_2_3&7 | 0.537 | 0.8668 | 0.8431 | 0.8319 | 0.8275 | 0.8327 | 0.8321 |
| 49 | FST_2_4&5 | 0.4638 | 0.8374 | 0.8126 | 0.8031 | 0.7954 | 0.8552 | 0.8556 |
| 50 | FST_2_4&6 | 0.1713 | 0.6363 | 0.6081 | 0.5972 | 0.5899 | 0.5964 | 0.5952 |
| 51 | FST_2_4&7 | 0.4565 | 0.836 | 0.811 | 0.8011 | 0.7943 | 0.7997 | 0.7993 |
| 52 | FST_2_5&6 | 0.4647 | 0.8394 | 0.8149 | 0.8053 | 0.7976 | 0.8032 | 0.8037 |
| 53 | FST_2_5&7 | 0.5997 | 0.8886 | 0.8672 | 0.8585 | 0.8534 | 0.8569 | 0.8572 |
| 54 | FST_2_6&7 | 0.0873 | 0.5171 | 0.5475 | 0.5384 | 0.5343 | 0.5387 | 0.5382 |
| 55 | NHA_3_1 | 4.5 | 0.7765 | 0.7753 | 0.7755 | 0.7753 | 0.7761 | 0.7756 |
| 56 | NHA_3_2 | 4.5714 | 0.7799 | 0.7782 | 0.7788 | 0.7786 | 0.779 | 0.7804 |
| 57 | NHA_3_3 | 5.7857 | 0.8094 | 0.8082 | 0.8096 | 0.8098 | 0.8102 | 0.81 |
| 58 | NHA_3_4 | 4.4286 | 0.7884 | 0.7871 | 0.7875 | 0.7886 | 0.7891 | 0.7888 |
| 59 | NHA_3_5 | 5.4286 | 0.7946 | 0.7932 | 0.7935 | 0.7934 | 0.7953 | 0.7953 |
| 60 | NHA_3_6 | 4.3571 | 0.7829 | 0.7817 | 0.782 | 0.7835 | 0.784 | 0.782 |
| 61 | NHA_3_7 | 3.8571 | 0.768 | 0.7681 | 0.7686 | 0.7685 | 0.7689 | 0.7691 |
| 62 | MPD_3_1 | 2.1825 | 0.7956 | 0.7949 | 0.7951 | 0.7955 | 0.7957 | 0.7956 |
| 63 | MPD_3_2 | 1.5678 | 0.7747 | 0.7736 | 0.7743 | 0.7744 | 0.7748 | 0.7759 |
| 64 | MPD_3_3 | 2.2618 | 0.798 | 0.797 | 0.7988 | 0.799 | 0.7993 | 0.7992 |
| 65 | MPD_3_4 | 1.697 | 0.7798 | 0.7789 | 0.7798 | 0.7807 | 0.7813 | 0.7811 |
| 66 | MPD_3_5 | 2.6094 | 0.8069 | 0.8061 | 0.8069 | 0.8068 | 0.8084 | 0.8081 |
| 67 | MPD_3_6 | 1.8085 | 0.7838 | 0.7828 | 0.7838 | 0.7852 | 0.7851 | 0.7837 |
| 68 | MPD_3_7 | 1.8642 | 0.7857 | 0.7862 | 0.7869 | 0.7868 | 0.7872 | 0.7873 |
| 69 | NS2_3_1&2 | 7.8571 | 0.7553 | 0.7509 | 0.7502 | 0.7493 | 0.7506 | 0.7585 |
| 70 | NS2_3_1&3 | 7.7857 | 0.755 | 0.7506 | 0.7505 | 0.7496 | 0.7584 | 0.7583 |
| 71 | NS2_3_1&4 | 8 | 0.7591 | 0.7546 | 0.754 | 0.7614 | 0.7626 | 0.7624 |
| 72 | NS2_3_1&5 | 8.2857 | 0.7575 | 0.7531 | 0.7526 | 0.7515 | 0.761 | 0.7608 |
| 73 | NS2_3_1&6 | 7.8571 | 0.7574 | 0.7528 | 0.7523 | 0.7517 | 0.7532 | 0.7525 |
| 74 | NS2_3_1&7 | 7.4286 | 0.7538 | 0.7496 | 0.749 | 0.7483 | 0.7494 | 0.7493 |
| 75 | NS2_3_2&3 | 7.9286 | 0.7565 | 0.752 | 0.7519 | 0.7511 | 0.7522 | 0.7596 |
| 76 | NS2_3_2&4 | 8.2143 | 0.7612 | 0.7567 | 0.756 | 0.7556 | 0.7568 | 0.7643 |
| 77 | NS2_3_2&5 | 8.4286 | 0.7589 | 0.7545 | 0.7538 | 0.753 | 0.7546 | 0.7621 |
| 78 | NS2_3_2&6 | 7.4286 | 0.7541 | 0.7496 | 0.7488 | 0.7567 | 0.7576 | 0.7497 |
| 79 | NS2_3_2&7 | 7.1429 | 0.7515 | 0.7475 | 0.7469 | 0.7541 | 0.755 | 0.7477 |
| 80 | NS2_3_3&4 | 8.7143 | 0.7652 | 0.7606 | 0.7607 | 0.7602 | 0.7686 | 0.7684 |
| 81 | NS2_3_3&5 | 8.3571 | 0.7586 | 0.7542 | 0.7619 | 0.7611 | 0.762 | 0.7619 |
| 82 | NS2_3_3&6 | 8.5 | 0.7628 | 0.7585 | 0.7585 | 0.758 | 0.7592 | 0.7587 |
| 83 | NS2_3_3&7 | 8.2857 | 0.7612 | 0.7574 | 0.7571 | 0.7564 | 0.7576 | 0.7575 |
| 84 | NS2_3_4&5 | 8.6429 | 0.763 | 0.7587 | 0.7579 | 0.7576 | 0.7665 | 0.7662 |
| 85 | NS2_3_4&6 | 8.2857 | 0.764 | 0.7594 | 0.7586 | 0.7587 | 0.7602 | 0.7596 |
| 86 | NS2_3_4&7 | 7.8571 | 0.7606 | 0.7564 | 0.7557 | 0.7554 | 0.7566 | 0.7565 |
| 87 | NS2_3_5&6 | 8.5714 | 0.7619 | 0.7574 | 0.7569 | 0.7565 | 0.7584 | 0.7577 |
| 88 | NS2_3_5&7 | 8.5 | 0.7612 | 0.7573 | 0.7567 | 0.756 | 0.7576 | 0.7576 |
| 89 | NS2_3_6&7 | 6.2857 | 0.7458 | 0.749 | 0.7488 | 0.7483 | 0.7492 | 0.7492 |
| 90 | HST_3_1&2 | 0.1898 | 0.5875 | 0.5296 | 0.5136 | 0.5053 | 0.5131 | 0.6245 |
| 91 | HST_3_1&3 | 0.05 | 0.4808 | 0.463 | 0.46 | 0.4573 | 0.4931 | 0.493 |
| 92 | HST_3_1&4 | 0.2451 | 0.642 | 0.575 | 0.551 | 0.6769 | 0.6885 | 0.6885 |
| 93 | HST_3_1&5 | 0.1251 | 0.531 | 0.493 | 0.4853 | 0.4802 | 0.555 | 0.5552 |
| 94 | HST_3_1&6 | 0.2351 | 0.6331 | 0.567 | 0.5439 | 0.5286 | 0.5397 | 0.5438 |
| 95 | HST_3_1&7 | 0.253 | 0.6542 | 0.5814 | 0.5564 | 0.543 | 0.5561 | 0.556 |
| 96 | HST_3_2&3 | 0.1053 | 0.5147 | 0.4825 | 0.4763 | 0.4724 | 0.4757 | 0.5354 |
| 97 | HST_3_2&4 | 0.2903 | 0.6965 | 0.6263 | 0.5984 | 0.5751 | 0.5914 | 0.7468 |
| 98 | HST_3_2&5 | 0.1405 | 0.5418 | 0.4984 | 0.4892 | 0.4842 | 0.487 | 0.5692 |
| 99 | HST_3_2&6 | 0.1581 | 0.5594 | 0.5107 | 0.4992 | 0.5813 | 0.5886 | 0.4957 |
| 100 | HST_3_2&7 | 0.2932 | 0.7026 | 0.6251 | 0.5969 | 0.7434 | 0.7545 | 0.5854 |
| 101 | HST_3_3&4 | 0.2405 | 0.6416 | 0.575 | 0.5445 | 0.529 | 0.6893 | 0.6893 |
| 102 | HST_3_3&5 | 0.0735 | 0.4918 | 0.4679 | 0.5071 | 0.5024 | 0.5073 | 0.5069 |
| 103 | HST_3_3&6 | 0.1512 | 0.5576 | 0.5115 | 0.4989 | 0.4927 | 0.4967 | 0.4976 |
| 104 | HST_3_3&7 | 0.2366 | 0.637 | 0.5656 | 0.5381 | 0.5267 | 0.5372 | 0.5367 |
| 105 | HST_3_4&5 | 0.1761 | 0.5727 | 0.5196 | 0.5057 | 0.4979 | 0.6074 | 0.6077 |
| 106 | HST_3_4&6 | 0.3876 | 0.8014 | 0.743 | 0.7149 | 0.6739 | 0.6924 | 0.705 |
| 107 | HST_3_4&7 | 0.4071 | 0.8202 | 0.7552 | 0.7273 | 0.6975 | 0.7168 | 0.716 |
| 108 | HST_3_5&6 | 0.1129 | 0.5253 | 0.4898 | 0.4831 | 0.4796 | 0.4809 | 0.4807 |
| 109 | HST_3_5&7 | 0.1354 | 0.5389 | 0.4972 | 0.4884 | 0.4835 | 0.4865 | 0.4859 |
| 110 | HST_3_6&7 | 0.2309 | 0.6364 | 0.7011 | 0.681 | 0.6706 | 0.6813 | 0.6811 |
| 111 | NHA_4_1 | 1 | 0.3142 | 0.3138 | 0.3141 | 0.3139 | 0.314 | 0.3146 |
| 112 | NHA_4_2 | 1 | 0.3141 | 0.314 | 0.3141 | 0.3142 | 0.3139 | 0.315 |
| 113 | NHA_4_3 | 1 | 0.3152 | 0.3151 | 0.3157 | 0.3156 | 0.3157 | 0.3165 |
| 114 | NHA_4_4 | 1 | 0.3142 | 0.3139 | 0.314 | 0.3145 | 0.3145 | 0.3151 |
| 115 | NHA_4_5 | 1 | 0.3141 | 0.3139 | 0.3138 | 0.3139 | 0.3146 | 0.3151 |
| 116 | NHA_4_6 | 1 | 0.3142 | 0.3137 | 0.3139 | 0.3146 | 0.3145 | 0.3143 |
| 117 | NHA_4_7 | 1 | 0.3165 | 0.3168 | 0.3168 | 0.3172 | 0.3172 | 0.3175 |
| 118 | MPD_4_1 | 0 | 0.3142 | 0.3138 | 0.3141 | 0.3139 | 0.314 | 0.3146 |
| 119 | MPD_4_2 | 0 | 0.3141 | 0.314 | 0.3141 | 0.3142 | 0.3139 | 0.315 |
| 120 | MPD_4_3 | 0 | 0.3152 | 0.3151 | 0.3157 | 0.3156 | 0.3157 | 0.3165 |
| 121 | MPD_4_4 | 0 | 0.3142 | 0.3139 | 0.314 | 0.3145 | 0.3145 | 0.3151 |
| 122 | MPD_4_5 | 0 | 0.3141 | 0.3139 | 0.3138 | 0.3139 | 0.3146 | 0.3151 |
| 123 | MPD_4_6 | 0 | 0.3142 | 0.3137 | 0.3139 | 0.3146 | 0.3145 | 0.3143 |
| 124 | MPD_4_7 | 0 | 0.3165 | 0.3168 | 0.3168 | 0.3172 | 0.3172 | 0.3175 |

**Table S2.3:** Evaluation of the simulated data set to test for the scenarios of divergence: Posterior error rate.

| Scenario | Type I error | Type II error |
| --- | --- | --- |
| S1 | 0.092 | 0.398 |
| S2 | 0.672 | 0.556 |
| S3 | 0.472 | 0.196 |
| S4 | 0.325 | 0.129 |
| S5 | 0.202 | 0.121 |
| S6 | 0.237 | 0.174 |

Type I error, the probability that the scenario 1 was not selected although it was the true model, was 0.09. Type II error, the probability that scenario 1 was selected although it was not the true model, was 0.397. One-third of all Type II errors was caused by selecting scenario 1 while scenario 2 (six lineages) was true.

**Table S2.4**: Estimates of the ratio N_current_/N_ancestral_.

Estimates are from the original parameters (one estimate of N_current_/N_ancestral_ per population) and from the composite parameters ( N_current_*µ / N_ancestral_ *µ ) (four estimates, one per marker, per population).

Values in red highlight cases where an event of contraction is support at 95 % by the credible interval (values are strictly lower than 1). Values in green, highlight cases where an event of contraction is support at 90 % by the credible interval. We do not observe event of expansion (values strictly higher than 1 in the credible interval) but rather one population, SCG_02, at the demographic equilibrium ( credible interval encompasses values both lower and higher than 1).

| Data type | genome | Current Population | Country | Median | Mean | 2.5% | 5% | 25% | 75% | 95% | 97.5% | MODE |
| --- | --- | --- | --- | --- | --- | --- | --- | --- | --- | --- | --- | --- |
| Composite | **NuSSR** | **HRV-01** | Croatia | 0.214 | 0.115 | 0.016 | 0.019 | 0.037 | 0.115 | 0.396 | 0.639 | 0.001 |
| Composite | cpSSR | HRV-01 | Croatia | 5.133 | 1.829 | 0.038 | 0.055 | 0.208 | 1.829 | 12.223 | 25.855 | 0.278 |
| Composite | gene | HRV-01 | Croatia | 0.756 | 0.253 | 0.002 | 0.005 | 0.031 | 0.253 | 1.261 | 2.403 | 0.009 |
| Composite | Organelle gene | HRV-01 | Croatia | 40.718 | 0.187 | 0.002 | 0.003 | 0.022 | 0.187 | 1.311 | 4.829 | 0.002 |
| Original | ALL | HRV-01 | Croatia | 0.622 | 0.666 | 0.247 | 0.281 | 0.407 | 0.666 | 1.053 | 1.282 | 0.238 |
| Composite | **NuSSR** | **ITA-01** | Italy | 0.236 | 0.126 | 0.015 | 0.019 | 0.037 | 0.126 | 0.428 | 0.693 | 0.001 |
| Composite | cpSSR | ITA-01 | Italy | 9.907 | 3.285 | 0.047 | 0.078 | 0.334 | 3.285 | 23.801 | 55.160 | 0.921 |
| Composite | gene | ITA-01 | Italy | 9.733 | 1.359 | 0.035 | 0.062 | 0.275 | 1.359 | 6.242 | 12.847 | 0.323 |
| Composite | Organelle gene | ITA-01 | Italy | 412.521 | 0.348 | 0.000 | 0.000 | 0.007 | 0.348 | 12.119 | 40.277 | 0.000 |
| Original | ALL | ITA-01 | Italy | 0.765 | 0.807 | 0.337 | 0.379 | 0.529 | 0.807 | 1.196 | 1.486 | 0.368 |
| Composite | NuSSR | SCG-02 | Serbia | 1.295 | 0.536 | 0.012 | 0.017 | 0.067 | 0.536 | 2.442 | 4.190 | 0.031 |
| Composite | cpSSR | SCG-02 | Serbia | 19.131 | 6.273 | 0.116 | 0.171 | 0.669 | 6.273 | 43.755 | 97.688 | 4.483 |
| Composite | gene | SCG-02 | Serbia | 18.585 | 3.702 | 0.441 | 0.611 | 1.260 | 3.702 | 16.190 | 30.401 | 3.174 |
| Composite | Organelle gene | SCG-02 | Serbia | 448.813 | 0.711 | 0.001 | 0.003 | 0.031 | 0.711 | 10.882 | 31.838 | 0.006 |
| Original | ALL | SCG-02 | Serbia | 1.059 | 1.042 | 0.589 | 0.639 | 0.794 | 1.042 | 1.524 | 1.950 | 0.745 |
| Composite | NuSSR | CYP-01 | Cyprus | 0.483 | 0.236 | 0.032 | 0.040 | 0.081 | 0.236 | 0.719 | 1.115 | 0.006 |
| Composite | cpSSR | CYP-01 | Cyprus | 0.245 | 0.234 | 0.017 | 0.025 | 0.062 | 0.234 | 0.739 | 1.161 | 0.006 |
| Composite | **gene** | **CYP-01** | Cyprus | 0.419 | 0.071 | 0.001 | 0.002 | 0.011 | 0.071 | 0.390 | 0.818 | 0.000 |
| Composite | Organelle gene | CYP-01 | Cyprus | 4.931 | 0.032 | 0.000 | 0.000 | 0.003 | 0.032 | 0.294 | 1.036 | 0.000 |
| Original | ALL | CYP-01 | Cyprus | 0.629 | 0.670 | 0.257 | 0.288 | 0.409 | 0.670 | 1.068 | 1.326 | 0.202 |
| Composite | **NuSSR** | CRIMEA-01 | CRIMEA | 0.210 | 0.130 | 0.014 | 0.018 | 0.036 | 0.130 | 0.458 | 0.830 | 0.001 |
| Composite | cpSSR | CRIMEA-01 | CRIMEA | 23.672 | 5.623 | 0.109 | 0.171 | 0.638 | 5.623 | 37.112 | 84.634 | 3.446 |
| Composite | gene | CRIMEA-01 | CRIMEA | 14.458 | 2.588 | 0.302 | 0.416 | 0.872 | 2.588 | 11.152 | 21.306 | 1.604 |
| Composite | Organelle gene | CRIMEA-01 | CRIMEA | 5049.571 | 0.365 | 0.000 | 0.002 | 0.024 | 0.365 | 4.434 | 10.382 | 0.005 |
| Original | ALL | CRIMEA-01 | CRIMEA | 0.788 | 0.824 | 0.370 | 0.409 | 0.553 | 0.824 | 1.211 | 1.501 | 0.383 |
| Composite | NuSSR | FRA-01 | France | 0.496 | 0.219 | 0.011 | 0.016 | 0.049 | 0.219 | 0.846 | 1.442 | 0.003 |
| Composite | cpSSR | FRA-01 | France | 7.854 | 1.860 | 0.037 | 0.055 | 0.213 | 1.860 | 12.343 | 27.537 | 0.391 |
| Composite | gene | FRA-01 | France | 8.139 | 1.061 | 0.027 | 0.047 | 0.209 | 1.061 | 4.820 | 9.868 | 0.261 |
| Composite | Organelle gene | FRA-01 | France | 22.519 | 0.064 | 0.000 | 0.001 | 0.005 | 0.064 | 0.600 | 2.301 | 0.000 |
| Original | ALL | FRA-01 | France | 0.551 | 0.592 | 0.208 | 0.235 | 0.346 | 0.592 | 0.969 | 1.173 | 0.152 |
| Composite | NuSSR | DZA-01 | Algeria | 0.091 | 0.048 | 0.003 | 0.004 | 0.010 | 0.048 | 0.233 | 0.418 | 0.000 |
| Composite | cpSSR | DZA-01 | Algeria | 8.666 | 2.815 | 0.050 | 0.074 | 0.303 | 2.815 | 19.724 | 44.899 | 0.757 |
| Composite | gene | DZA-01 | Algeria | 1.527 | 0.488 | 0.019 | 0.033 | 0.120 | 0.488 | 1.754 | 3.196 | 0.054 |
| Composite | Organelle gene | DZA-01 | Algeria | 15448.350 | 26.384 | 0.007 | 0.031 | 1.033 | 26.384 | 275.817 | 810.778 | 73.550 |
| Original | ALL | DZA-01 | Algeria | 0.479 | 0.524 | 0.171 | 0.198 | 0.298 | 0.524 | 0.875 | 1.099 | 0.120 |

**Table S2.5**: Accuracy test of parameter estimation for the scenarios of divergence

Accuracy is measured by the mean relative bias (MRB) and the relative root mean square (RMSE) while precision is measured by the coverage values and factor 2.

| **Parameter** |  | **MRB (Median)** | **MRB (Mode)** | **RMSE (Median)** | **RMSE (Mode)** | **50% Coverage** | **95% Coverage** | **Factor 2 (Median)** | **Factor 2 (Mode)** |
| --- | --- | --- | --- | --- | --- | --- | --- | --- | --- |
| Current size N_0_ | HRV-01 | 0.1607 | 0.1284 | 0.338 | 0.322 | 0.534 | 0.974 | 0.99 | 0.984 |
| Current size N_0_ | ITA-01 | 0.0895 | 0.0707 | 0.274 | 0.268 | 0.524 | 0.966 | 0.996 | 0.996 |
| Current size N_0_ | SCG-02 | -0.0663 | -0.0648 | 0.188 | 0.199 | 0.406 | 0.956 | 0.998 | 0.998 |
| Current size N_0_ | CYP-01 | 0.166 | 0.1134 | 0.356 | 0.336 | 0.538 | 0.958 | 0.984 | 0.986 |
| Current size N_0_ | CRIMEA-01 | 0.0669 | 0.0464 | 0.253 | 0.253 | 0.538 | 0.974 | 0.994 | 0.992 |
| Current size N_0_ | FRA-01 | 0.1531 | 0.0692 | 0.36 | 0.327 | 0.578 | 0.966 | 0.98 | 0.978 |
| Current size N_0_ | DZA-01 | 0.1571 | 0.0706 | 0.369 | 0.348 | 0.546 | 0.976 | 0.97 | 0.962 |
| Ancestral size N_1_ | Anc | -0.1527 | -0.1372 | 0.291 | 0.268 | 0.308 | 0.822 | 0.96 | 0.948 |
| Divergence time | T | -0.0316 | -0.1642 | 0.355 | 0.337 | 0.558 | 0.98 | 0.96 | 0.916 |
| mutation rate µ | nSSR | 0.724 | 0.2755 | 1.087 | 0.704 | 0.354 | 0.86 | 0.706 | 0.896 |
| mutation rate µ | cpSSR | 0.0556 | -0.3598 | 2.229 | 2.222 | 0.582 | 0.966 | 0.668 | 0.436 |
| mutation rate µ | Nuclear genes | 0.4605 | -0.1139 | 3.4 | 0.566 | 0.43 | 0.936 | 0.828 | 0.746 |
| mutation rate µ | Organelle genes | 49.2425 | -0.3977 | 307.204 | 0.63 | 0.828 | 0.996 | 0.566 | 0.602 |
| mutation rate's parameter | nSSR (geometric distribution) | 2.567 | -0.8112 | 5.988 | 0.851 | 0.508 | 0.952 | 0.234 | 0.132 |
| mutation rate's parameter | cpSSR (geometric distribution) | 0.5047 | 0.6726 | 1.213 | 1.852 | 0.546 | 0.958 | 0.792 | 0.546 |
| mutation rate's parameter | nuclear genes (HKY) | 0.1609 | 0.2916 | 0.801 | 1.271 | 0.54 | 0.962 | 0.862 | 0.474 |
| mutation rate's parameter | Organelle genes (HKY) | 2.6374 | -0.8113 | 6.09 | 0.855 | 0.476 | 0.952 | 0.24 | 0.136 |

**Table S2.6**: Model checking for the best scenario of divergence (Scenario 1)

The Probability (SSsimulated<SSobserved) is given for each summary statistics. To avoid the risk of over-estimating the quality of the fit we used different summary statistics (8 in total) than the ones used during the inference steps (model discrimination and posterior estimation of parameters). The simulated summary statistics were obtained from 10,000 data sets that were simulated from the posterior distribution of parameters obtained under Scenario 1.

The code used for the summary statistics is composed of three parts: the summary statistic (see above for the definitions), the type of marker (1,2,3,4 for nSSR, cpSSR, nuclear gene, organelle gene respectively), the population (1,2,3,4,5,6,7 for HRV_01, ITA_01, SCG_02, CYP_01, UKR_01, DZA_01, CYP_01 respectively)

*HET*: mean genic diversity within population, *H2P_:_* mean genic diversity between pairs of populations, *NAL*: mean number of alleles within population, *N2P*: mean number of alleles between pairs of populations, *NSS*: mean number of segregating sites within population , *NH2*: mean number of haplotypes between pairs of populations, *MPB*: mean of pairwise differences between pairs of populations

| Summary statistics | Observed value | Probability (SSsimulated<SSobserved) |
| --- | --- | --- |
| NAL_1_1 | 6.4286 | 0.0925 |
| NAL_1_2 | 8.3571 | 0.4855 |
| NAL_1_3 | 7.7857 | 0.282 |
| NAL_1_4 | 7.4286 | 0.3375 |
| NAL_1_5 | 7.3571 | 0.1835 |
| NAL_1_6 | 6.4286 | 0.169 |
| NAL_1_7 | 6.2143 | 0.3015 |
| H2P_1_1&2 | 0.7827 | 0.117 |
| H2P_1_1&3 | 0.7645 | 0.0655 |
| H2P_1_1&4 | 0.7806 | 0.1185 |
| H2P_1_1&5 | 0.7754 | 0.0975 |
| H2P_1_1&6 | 0.7946 | 0.1945 |
| H2P_1_1&7 | 0.7827 | 0.1495 |
| H2P_1_2&3 | 0.7869 | 0.107 |
| H2P_1_2&4 | 0.8124 | 0.28 |
| H2P_1_2&5 | 0.7945 | 0.151 |
| H2P_1_2&6 | 0.7879 | 0.1465 |
| H2P_1_2&7 | 0.7923 | 0.1785 |
| H2P_1_3&4 | 0.7875 | 0.1245 |
| H2P_1_3&5 | 0.7663 | 0.058 |
| H2P_1_3&6 | 0.7891 | 0.149 |
| H2P_1_3&7 | 0.7836 | 0.124 |
| H2P_1_4&5 | 0.788 | 0.1395 |
| H2P_1_4&6 | 0.7913 | 0.183 |
| H2P_1_4&7 | 0.7758 | 0.122 |
| H2P_1_5&6 | 0.7906 | 0.161 |
| H2P_1_5&7 | 0.7796 | 0.1265 |
| H2P_1_6&7 | 0.766 | 0.0985 |
| HET_2_1 | 0.303 | 0.2635 |
| HET_2_2 | 0.7576 | 0.886 |
| HET_2_3 | 0.5636 | 0.474 |
| HET_2_4 | 0.6444 | 0.7025 |
| HET_2_5 | 0.4394 | 0.352 |
| HET_2_6 | 0.6364 | 0.7555 |
| HET_2_7 | 0.3556 | 0.3795 |
| N2P_2_1&2 | 5 | 0.705 |
| N2P_2_1&3 | 5 | 0.686 |
| N2P_2_1&4 | 4 | 0.506 |
| N2P_2_1&5 | 5 | 0.706 |
| N2P_2_1&6 | 3 | 0.309 |
| N2P_2_1&7 | 2 | 0.141 |
| N2P_2_2&3 | 6 | 0.8395 |
| N2P_2_2&4 | 7 | 0.943 |
| N2P_2_2&5 | 6 | 0.8455 |
| N2P_2_2&6 | 6 | 0.8725 |
| N2P_2_2&7 | 5 | 0.769 |
| N2P_2_3&4 | 6 | 0.8555 |
| N2P_2_3&5 | 3 | 0.239 |
| N2P_2_3&6 | 6 | 0.861 |
| N2P_2_3&7 | 5 | 0.747 |
| N2P_2_4&5 | 6 | 0.8645 |
| N2P_2_4&6 | 4 | 0.539 |
| N2P_2_4&7 | 4 | 0.5785 |
| N2P_2_5&6 | 6 | 0.8685 |
| N2P_2_5&7 | 5 | 0.7485 |
| N2P_2_6&7 | 3 | 0.362 |
| NSS_3_1 | 6.5714 | 0.0865 |
| NSS_3_2 | 6.0714 | 0.078 |
| NSS_3_3 | 7.2143 | 0.0835 |
| NSS_3_4 | 5.6429 | 0.084 |
| NSS_3_5 | 7.3571 | 0.088 |
| NSS_3_6 | 5.2857 | 0.082 |
| NSS_3_7 | 4.8571 | 0.081 |
| NH2_3_1&2 | 6.5 | 0.0745 |
| NH2_3_1&3 | 7.4286 | 0.0885 |
| NH2_3_1&4 | 6.5714 | 0.0845 |
| NH2_3_1&5 | 6.8571 | 0.079 |
| NH2_3_1&6 | 6.7143 | 0.0865 |
| NH2_3_1&7 | 6.1429 | 0.085 |
| NH2_3_2&3 | 7.6429 | 0.089 |
| NH2_3_2&4 | 7.1429 | 0.09 |
| NH2_3_2&5 | 7.3571 | 0.084 |
| NH2_3_2&6 | 6.7143 | 0.0865 |
| NH2_3_2&7 | 6.5 | 0.086 |
| NH2_3_3&4 | 8.4286 | 0.111 |
| NH2_3_3&5 | 8.2857 | 0.0905 |
| NH2_3_3&6 | 8.0714 | 0.1085 |
| NH2_3_3&7 | 7.8571 | 0.1035 |
| NH2_3_4&5 | 7.6429 | 0.0945 |
| NH2_3_4&6 | 7.2857 | 0.112 |
| NH2_3_4&7 | 7.0714 | 0.1025 |
| NH2_3_5&6 | 7.5 | 0.0905 |
| NH2_3_5&7 | 7.2857 | 0.0905 |
| NH2_3_6&7 | 6.0714 | 0.0895 |
| MPB_3_1&2 | 2.3038 | 0.087 |
| MPB_3_1&3 | 2.3399 | 0.084 |
| MPB_3_1&4 | 2.7366 | 0.096 |
| MPB_3_1&5 | 2.7234 | 0.098 |
| MPB_3_1&6 | 2.778 | 0.099 |
| MPB_3_1&7 | 2.8899 | 0.1065 |
| MPB_3_2&3 | 2.1475 | 0.084 |
| MPB_3_2&4 | 2.3184 | 0.086 |
| MPB_3_2&5 | 2.4751 | 0.086 |
| MPB_3_2&6 | 1.9978 | 0.081 |
| MPB_3_2&7 | 2.4138 | 0.09 |
| MPB_3_3&4 | 2.7718 | 0.097 |
| MPB_3_3&5 | 2.6615 | 0.093 |
| MPB_3_3&6 | 2.5065 | 0.091 |
| MPB_3_3&7 | 2.8466 | 0.109 |
| MPB_3_4&5 | 2.8497 | 0.105 |
| MPB_3_4&6 | 2.7847 | 0.1 |
| MPB_3_4&7 | 2.9674 | 0.107 |
| MPB_3_5&6 | 2.6441 | 0.096 |
| MPB_3_5&7 | 2.7331 | 0.0975 |
| MPB_3_6&7 | 2.3847 | 0.087 |
| NSS_4_1 | 0 | 0.473 |
| NSS_4_2 | 0 | 0.468 |
| NSS_4_3 | 0 | 0.466 |
| NSS_4_4 | 0 | 0.479 |
| NSS_4_5 | 0 | 0.4675 |
| NSS_4_6 | 0 | 0.4745 |
| NSS_4_7 | 0 | 0.476 |

**Table S2.7:** Evaluation of the simulated datasets to infer admixture parameters: test of rank.

Each summary statistic (30 in total) of the observed data set is ranked against those of the simulated data set. Values indicate for each summary statistics the proportion of simulated data sets which have a value below the observed one.

|  | Three population model: | CYP-01, CRIMEA-01, TUR-01 | | FRA-01, DZA-01, ESP-01 | | SCG-02, HRV-01, ROU-01 | | SCG-02, CRIMEA-01, HRV-02 | | ITA-01, SCG-02, ITA-02 | |
| --- | --- | --- | --- | --- | --- | --- | --- | --- | --- | --- | --- |
|  | Summary statistic | Observed | Simulated | Observed | Simulated | Observed | Simulated | Observed | Simulated | Observed | Simulated |
| 1 | HET_1_1 | 0.7496 | 0.3169 | 0.7254 | 0.2767 | 0.7357 | 0.2923 | 0.7566 | 0.3289 | 0.77 | 0.3529 |
| 2 | HET_1_2 | 0.7469 | 0.3125 | 0.742 | 0.3044 | 0.7566 | 0.3282 | 0.7469 | 0.312 | 0.7566 | 0.329 |
| 3 | HET_1_3 | 0.7542 | 0.3146 | 0.7294 | 0.2724 | 0.6905 | 0.2115 | 0.7675 | 0.3382 | 0.7685 | 0.3396 |
| 4 | FST_1_1&2 | 0.0929 | 0.4664 | 0.1013 | 0.4403 | 0.0454 | 0.2013 | 0.0345 | 0.1148 | 0.0537 | 0.26 |
| 5 | FST_1_1&3 | 0.0822 | 0.4637 | 0.0633 | 0.2313 | 0.103 | 0.5672 | 0.0006 | 0.0003*** | 0.0083 | 0.0013** |
| 6 | FST_1_2&3 | 0.0427 | 0.2139 | 0 | 0.5 | 0.0466 | 0.2513 | 0.0462 | 0.2477 | 0.0457 | 0.217 |
| 7 | NAL_2_1 | 3 | 0.6333 | 3 | 0.6244 | 2 | 0.4986 | 3 | 0.6244 | 5 | 0.7699 |
| 8 | NAL_2_2 | 3 | 0.6178 | 2 | 0.5093 | 3 | 0.6251 | 3 | 0.6178 | 3 | 0.6248 |
| 9 | NAL_2_3 | 4 | 0.6976 | 2 | 0.4915 | 4 | 0.6976 | 4 | 0.7068 | 3 | 0.6205 |
| 10 | FST_2_1&2 | 0.4638 | 0.8121 | 0.0873 | 0.4899 | 0.5707 | 0.858 | -0.0335 | 0.01* | 0.2832 | 0.7098 |
| 11 | FST_2_1&3 | -0.0677 | 0.0044** | 0.418 | 0.8228 | 0.5076 | 0.862 | -0.0507 | 0.0073** | 0.2413 | 0.7084 |
| 12 | FST_2_2&3 | 0.4091 | 0.8181 | 0.5516 | 0.8781 | 0.0089 | 0.3807 | -0.071 | 0.0028** | -0.0838 | 0.0017** |
| 13 | NHA_3_1 | 4.4286 | 0.7875 | 4.3571 | 0.7826 | 4.5 | 0.7763 | 5.7857 | 0.8089 | 4.5714 | 0.7796 |
| 14 | NHA_3_2 | 5.4286 | 0.7939 | 3.8571 | 0.7677 | 5.7857 | 0.809 | 5.4286 | 0.794 | 5.7857 | 0.8095 |
| 15 | NHA_3_3 | 5.0714 | 0.788 | 4.7857 | 0.7817 | 5.5714 | 0.7933 | 5.2857 | 0.7983 | 5 | 0.7852 |
| 16 | MPD_3_1 | 1.697 | 0.7791 | 1.8085 | 0.7832 | 2.1825 | 0.7957 | 2.2618 | 0.7975 | 1.5678 | 0.7741 |
| 17 | MPD_3_2 | 2.6094 | 0.8067 | 1.8642 | 0.7851 | 2.2618 | 0.7977 | 2.6094 | 0.8065 | 2.2618 | 0.7981 |
| 18 | MPD_3_3 | 2.0375 | 0.7861 | 1.819 | 0.7784 | 2.3595 | 0.7955 | 2.1574 | 0.7894 | 2.0606 | 0.7868 |
| 19 | NS2_3_1&2 | 8.6429 | 0.7597 | 6.2857 | 0.7425 | 7.7857 | 0.7517 | 8.3571 | 0.7554 | 7.9286 | 0.7536 |
| 20 | NS2_3_1&3 | 7.7143 | 0.7559 | 6.5 | 0.7446 | 8 | 0.7538 | 7.7143 | 0.7546 | 7.6429 | 0.7526 |
| 21 | NS2_3_2&3 | 8.1429 | 0.7554 | 6.0714 | 0.7402 | 8.2143 | 0.7561 | 7.8571 | 0.7543 | 7.8571 | 0.7547 |
| 22 | HST_3_1&2 | 0.1761 | 0.5206 | 0.2309 | 0.5716 | 0.05 | 0.4633 | 0.0735 | 0.4685 | 0.1053 | 0.4831 |
| 23 | HST_3_1&3 | 0.1335 | 0.5451 | 0.0925 | 0.5116 | 0.0713 | 0.4904 | 0.0628 | 0.4809 | 0.0246 | 0.4661 |
| 24 | HST_3_2&3 | 0.0999 | 0.5067 | 0.1167 | 0.5286 | 0.0515 | 0.4809 | 0.058 | 0.4797 | 0.0597 | 0.4849 |
| 25 | NHA_4_1 | 1 | 0.3144 | 1 | 0.3143 | 1 | 0.3143 | 1 | 0.3151 | 1 | 0.3141 |
| 26 | NHA_4_2 | 1 | 0.3143 | 1 | 0.3169 | 1 | 0.3154 | 1 | 0.3141 | 1 | 0.3154 |
| 27 | NHA_4_3 | 1 | 0.3122 | 1 | 0.3119 | 1 | 0.3118 | 1 | 0.3127 | 1 | 0.3129 |
| 28 | MPD_4_1 | 0 | 0.3144 | 0 | 0.3143 | 0 | 0.3143 | 0 | 0.3151 | 0 | 0.3141 |
| 29 | MPD_4_2 | 0 | 0.3143 | 0 | 0.3169 | 0 | 0.3154 | 0 | 0.3141 | 0 | 0.3154 |
| 30 | MPD_4_3 | 0 | 0.3122 | 0 | 0.3119 | 0 | 0.3118 | 0 | 0.3127 | 0 | 0.3129 |

The code used for the summary statistics is composed of three parts: the summary statistic (see above for the definitions), the type of marker (1,2,3,4 for nSSR, cpSSR, nuclear gene, organelle gene respectively), the population (1,2,3 respectively for CYP-01, CRIMEA-01, TUR-01, respectively for FRA-01, DZA-01, ESP-01 respectively for SCG-02, HRV-01, ROU-01 respectively for SCG-02, CRIMEA-01, HRV-02 respectively for ITA-01, SCG-02, ITA-02.

*HET*: mean genic diversity within population, *NAL*: mean number of alleles within population, *FST*: genetic differentiation between pair of populations, *NHA*: number of haplotypes within population, *MPD*: mean of pairwise differences within populations, *NS2*: number of segregating sites between pairs of populations , *HST* genetic differentiation based on sequence haplotypes between pairs of populations.

**Table S2.8:** Inference of demographic parameters in the scenarios with admixture

| **Country** | **Sub-species** | **Genetic structure within the population** | **Parameter** | **mean** | **median** | **mode** | **q025** | **q050** | **q250** | **q750** | **q950** | **q975** |
| --- | --- | --- | --- | --- | --- | --- | --- | --- | --- | --- | --- | --- |
|  |  |  | **Current demographic size N** |  |  |  |  |  |  |  |  |  |
| Croatia | *P.n.dalmatica* | *homogeneous* | HRV-01 | 4800 | 4640 | 4450 | 2220 | 2510 | 3660 | 5760 | 7750 | 8390 |
| Serbia | *P.n.nigra* | *homogeneous* | SCG-02 | 8070 | 8190 | 8600 | 5740 | 6170 | 7450 | 8830 | 9560 | 9740 |
| Romania | *P.n.nigra* | *admix* | ROU-01 | 3870 | 3580 | 3180 | 1340 | 1620 | 2620 | 4800 | 7260 | 8230 |
|  |  |  | **Ancestral demographic size N** | 7670 | 8130 | 8680 | 2840 | 4040 | 6990 | 8870 | 9570 | 9730 |
|  |  |  | **Merging time, t in generation** | 485 | 391 | 261 | 95.1 | 124 | 251 | 606 | 1160 | 1420 |
|  |  |  | **admixture rate** | 0.29 | 0.257 | 0.224 | 0.0379 | 0.0633 | 0.167 | 0.376 | 0.64 | 0.735 |
|  |  |  | **Divergence time, t in generation** | 687 | 481 | 355 | 146 | 175 | 314 | 769 | 1790 | 2550 |
|  |  |  | **mutation rate µ** |  |  |  |  |  |  |  |  |  |
|  |  |  | nSSR | 0.000316 | 0.000278 | 0.000218 | 0.000152 | 0.000167 | 0.000221 | 0.000362 | 0.000567 | 0.000672 |
|  |  |  | cpSSR | 0.000555 | 0.000434 | 0.000299 | 0.000145 | 0.000173 | 0.000295 | 0.000663 | 0.00129 | 0.00166 |
|  |  |  | nuclear genes | 0.00000358 | 0.00000258 | 2.49E-008 | 3.07E-008 | 6.45E-008 | 0.000000732 | 0.00000617 | 0.00000944 | 0.00000989 |
|  |  |  | cp barcode | 5.73E-010 | 1E-012 | 1E-012 | 1E-012 | 1E-012 | 1E-012 | 4.95E-010 | 3.09E-009 | 4.83E-009 |
|  |  |  | **mutation rate's parameter** |  |  |  |  |  |  |  |  |  |
|  |  |  | NSSR (geometric distribution) | 0.514 | 0.498 | 0.149 | 0.116 | 0.136 | 0.282 | 0.736 | 0.941 | 0.966 |
|  |  |  | cpSSR (geometric distribution) | 0.547 | 0.548 | 0.13 | 0.123 | 0.143 | 0.314 | 0.776 | 0.959 | 0.981 |
|  |  |  | nuclear genes (HKY) | 3.52 | 1.09 | 0.0567 | 0.0599 | 0.0713 | 0.244 | 4.78 | 15.2 | 17.7 |
|  |  |  | cp barcode (HKY) | 4.46 | 1.77 | 0.0589 | 0.0677 | 0.0857 | 0.364 | 6.89 | 17.2 | 18.8 |
|  |  |  | **Current demographic size N** |  |  |  |  |  |  |  |  |  |
| Italy | *P.n.laricio* | *homogeneous* | ITA-01 | 6970 | 7030 | 6840 | 4140 | 4600 | 6050 | 7950 | 9170 | 9500 |
| Serbia | *P.n.nigra* | *homogeneous* | SCG-02 | 7750 | 7860 | 7880 | 5310 | 5790 | 7050 | 8560 | 9400 | 9610 |
| Italy | *P.n.laricio* | *admix* | ITA-02 | 4640 | 4390 | 4060 | 1770 | 2100 | 3350 | 5730 | 7980 | 8730 |
|  |  |  | **Ancestral demographic size N** | 6710 | 7090 | 7780 | 1710 | 2680 | 5640 | 8170 | 9300 | 9540 |
|  |  |  | **Merging time, t in generation** | 137 | 105 | 53.5 | 26.8 | 33.4 | 65.8 | 168 | 343 | 449 |
|  |  |  | **admixture rate** | 0.512 | 0.513 | 0.504 | 0.116 | 0.177 | 0.381 | 0.644 | 0.839 | 0.895 |
|  |  |  | **Divergence time, t in generation** | 741 | 530 | 293 | 152 | 187 | 337 | 842 | 1970 | 2810 |
|  |  |  | **mutation rate µ** |  |  |  |  |  |  |  |  |  |
|  |  |  | nSSR | 0.000513 | 0.000407 | 0.000335 | 0.000205 | 0.000224 | 0.000314 | 0.000555 | 0.00106 | 0.00145 |
|  |  |  | cpSSR | 0.000924 | 0.000693 | 0.000504 | 0.000223 | 0.000269 | 0.000468 | 0.00106 | 0.00224 | 0.00314 |
|  |  |  | nuclear genes | 0.00000297 | 0.00000186 | 6.05E-009 | 1.92E-008 | 4.05E-008 | 0.000000496 | 0.00000494 | 0.00000896 | 0.00000955 |
|  |  |  | cp barcode | 1.57E-009 | 1E-012 | 1E-012 | 1E-012 | 1E-012 | 1E-012 | 1.21E-009 | 7.67E-009 | 0.000000013 |
|  |  |  | **mutation rate's parameter** |  |  |  |  |  |  |  |  |  |
|  |  |  | NSSR (geometric distribution) | 0.565 | 0.573 | 1 | 0.123 | 0.148 | 0.332 | 0.802 | 0.965 | 0.984 |
|  |  |  | cpSSR (geometric distribution) | 0.526 | 0.517 | 0.117 | 0.118 | 0.136 | 0.294 | 0.749 | 0.951 | 0.974 |
|  |  |  | nuclear genes (HKY) | 3 | 0.851 | 0.05 | 0.0564 | 0.0641 | 0.196 | 3.88 | 13.6 | 16.4 |
|  |  |  | cp barcode (HKY) | 2.8 | 0.749 | 0.05 | 0.0549 | 0.0618 | 0.175 | 3.44 | 13.2 | 16.1 |
|  |  |  | **Current demographic size N** |  |  |  |  |  |  |  |  |  |
| Francia | P.n.salzmannii | *homogeneous* | FRA-01 | 7040 | 7090 | 7360 | 4310 | 4780 | 6140 | 7980 | 9160 | 9510 |
| Algeria | P.n.salzmannii | *homogeneous* | DZA-01 | 5850 | 5800 | 5940 | 3090 | 3490 | 4800 | 6820 | 8470 | 8980 |
| Spain | P.n.salzmannii | *admix* | ESP-01 | 5740 | 5680 | 5720 | 2720 | 3140 | 4600 | 6850 | 8590 | 9090 |
|  |  |  | **Ancestral demographic size N** | 7100 | 7470 | 7780 | 2390 | 3400 | 6160 | 8440 | 9420 | 9630 |
|  |  |  | **Merging time, t in generation** | 359 | 285 | 166 | 55.5 | 77.8 | 176 | 455 | 877 | 1070 |
|  |  |  | **admixture rate** | 0.547 | 0.551 | 0.471 | 0.128 | 0.196 | 0.418 | 0.686 | 0.872 | 0.921 |
|  |  |  | **Divergence time, t in generation** | 1540 | 1240 | 895 | 379 | 465 | 839 | 1890 | 3670 | 4690 |
|  |  |  | **mutation rate µ** |  |  |  |  |  |  |  |  |  |
|  |  |  | nSSR | 0.000327 | 0.000289 | 0.000262 | 0.000154 | 0.00017 | 0.00023 | 0.000377 | 0.000575 | 0.000688 |
|  |  |  | cpSSR | 0.000293 | 0.000223 | 0.000171 | 0.0000552 | 0.0000758 | 0.000147 | 0.000346 | 0.000678 | 0.000886 |
|  |  |  | nuclear genes | 0.00000294 | 0.00000227 | 4.18E-009 | 2.18E-008 | 6.05E-008 | 0.000000938 | 0.00000443 | 0.00000805 | 0.00000877 |
|  |  |  | cp barcode | 2.25E-009 | 1E-012 | 1E-012 | 1E-012 | 1E-012 | 1E-012 | 1.73E-009 | 1.07E-008 | 1.68E-008 |
|  |  |  | **mutation rate's parameter** |  |  |  |  |  |  |  |  |  |
|  |  |  | NSSR (geometric distribution) | 0.558 | 0.561 | 1 | 0.124 | 0.147 | 0.328 | 0.791 | 0.963 | 0.982 |
|  |  |  | cpSSR (geometric distribution) | 0.574 | 0.585 | 1 | 0.126 | 0.149 | 0.346 | 0.807 | 0.973 | 0.992 |
|  |  |  | nuclear genes (HKY) | 3.41 | 1.04 | 0.0539 | 0.0589 | 0.0693 | 0.239 | 4.61 | 15 | 17.4 |
|  |  |  | cp barcode (HKY) | 3.44 | 1.11 | 0.0583 | 0.0594 | 0.0698 | 0.242 | 4.76 | 14.9 | 17.4 |
|  |  |  | **Current demographic size N** |  |  |  |  |  |  |  |  |  |
| Serbia | *P.n.nigra* | *homogeneous* | SCG-02 | 7190 | 7280 | 7550 | 4370 | 4840 | 6310 | 8140 | 9280 | 9560 |
| Ukraine | P.n.pallasiana | *homogeneous* | CRIMEA-01 | 5690 | 5590 | 5520 | 2810 | 3180 | 4520 | 6760 | 8580 | 9140 |
| Croatia | *P.n.dalmatica* | *admix* | HRV-02 | 5670 | 5610 | 5650 | 2500 | 2890 | 4460 | 6840 | 8710 | 9200 |
|  |  |  | **Ancestral demographic size N** | 7610 | 8040 | 8800 | 2550 | 3870 | 6910 | 8830 | 9560 | 9720 |
|  |  |  | **Merging time, t in generation** | 171 | 130 | 100 | 28.9 | 38.9 | 80.7 | 208 | 434 | 564 |
|  |  |  | **admixture rate** | 0.622 | 0.64 | 0.634 | 0.197 | 0.285 | 0.513 | 0.751 | 0.895 | 0.934 |
|  |  |  | **Divergence time, t in generation** | 425 | 304 | 223 | 89.5 | 109 | 201 | 481 | 1050 | 1470 |
|  |  |  | **mutation rate µ** |  |  |  |  |  |  |  |  |  |
|  |  |  | nSSR | 0.000561 | 0.000443 | 0.000342 | 0.000215 | 0.000237 | 0.000335 | 0.000619 | 0.0012 | 0.0016 |
|  |  |  | cpSSR | 0.000303 | 0.000107 | 0.0000115 | 0.00000546 | 0.00000843 | 0.0000402 | 0.00026 | 0.00101 | 0.00189 |
|  |  |  | nuclear genes | 0.00000462 | 0.00000439 | 8.1E-009 | 5.92E-008 | 0.000000155 | 0.00000203 | 0.00000721 | 0.00000961 | 0.00000993 |
|  |  |  | cp barcode | 1.17E-009 | 1E-012 | 1E-012 | 1E-012 | 1E-012 | 1E-012 | 9.33E-010 | 5.94E-009 | 9.74E-009 |
|  |  |  | **mutation rate's parameter** |  |  |  |  |  |  |  |  |  |
|  |  |  | NSSR (geometric distribution) | 0.521 | 0.509 | 0.112 | 0.114 | 0.131 | 0.285 | 0.75 | 0.943 | 0.971 |
|  |  |  | cpSSR (geometric distribution) | 0.617 | 0.639 | 1 | 0.13 | 0.157 | 0.382 | 0.877 | 0.998 | 1 |
|  |  |  | nuclear genes (HKY) | 2.89 | 0.778 | 0.05 | 0.056 | 0.0633 | 0.185 | 3.54 | 13.6 | 16.6 |
|  |  |  | cp barcode (HKY) | 3.35 | 1.02 | 0.0533 | 0.058 | 0.0682 | 0.222 | 4.52 | 15 | 17.4 |
|  |  |  | **Current demographic size N** |  |  |  |  |  |  |  |  |  |
| Cyprus | P.n.pallasiana | *homogeneous* | CYP-01 | 6240 | 6220 | 6490 | 3420 | 3880 | 5240 | 7260 | 8720 | 9120 |
| Ukraine | P.n.pallasiana | *homogeneous* | CRIMEA-01 | 6590 | 6610 | 6220 | 3780 | 4230 | 5640 | 7570 | 8890 | 9320 |
| Turkey | P.n.pallasiana | *admix* | TUR-01 | 5460 | 5360 | 4970 | 2330 | 2820 | 4240 | 6560 | 8490 | 9070 |
|  |  |  | **Ancestral demographic size N** | 7240 | 7600 | 8040 | 2560 | 3660 | 6370 | 8520 | 9450 | 9670 |
|  |  |  | **Merging time, t in generation** | 490 | 402 | 269 | 90.2 | 122 | 257 | 614 | 1140 | 1440 |
|  |  |  | **admixture rate** | 0.335 | 0.308 | 0.254 | 0.0561 | 0.0882 | 0.21 | 0.431 | 0.679 | 0.767 |
|  |  |  | **Divergence time, t in generation** | 1870 | 1520 | 1200 | 475 | 582 | 1030 | 2270 | 4380 | 5560 |
|  |  |  | **mutation rate µ** |  |  |  |  |  |  |  |  |  |
|  |  |  | nSSR | 0.000423 | 0.000355 | 0.000291 | 0.000183 | 0.0002 | 0.000276 | 0.000473 | 0.000808 | 0.000991 |
|  |  |  | cpSSR | 0.000273 | 0.000191 | 0.000117 | 0.0000366 | 0.0000531 | 0.000118 | 0.000311 | 0.00065 | 0.000881 |
|  |  |  | nuclear genes | 0.00000302 | 0.00000222 | 5.54E-009 | 3.09E-008 | 7.14E-008 | 0.000000912 | 0.00000461 | 0.00000844 | 0.0000091 |
|  |  |  | cp barcode | 1.76E-009 | 1E-012 | 1E-012 | 1E-012 | 1E-012 | 1E-012 | 1.42E-009 | 8.66E-009 | 1.43E-008 |
|  |  |  | **mutation rate's parameter** |  |  |  |  |  |  |  |  |  |
|  |  |  | NSSR (geometric distribution) | 0.544 | 0.539 | 0.148 | 0.12 | 0.14 | 0.314 | 0.773 | 0.957 | 0.979 |
|  |  |  | cpSSR (geometric distribution) | 0.552 | 0.545 | 0.161 | 0.122 | 0.143 | 0.315 | 0.791 | 0.973 | 0.992 |
|  |  |  | nuclear genes (HKY) | 3.03 | 0.871 | 0.0609 | 0.0566 | 0.0631 | 0.198 | 3.88 | 14 | 16.7 |
|  |  |  | cp barcode (HKY) | 3.62 | 1.18 | 0.0545 | 0.0607 | 0.0716 | 0.251 | 5.05 | 15.8 | 17.9 |

**Table S2.9:** Accuracy test of parameters estimation for the scenarios of admixture

Accuracy is measured by the mean relative bias (MRB) and the relative root mean square (RMSE) while precision is measured by the coverage values and factor 2.

| **Countries**  **sub-species** | **P*** | **Scenario** | **MRB (Median)** | **MRB (Mode)** | **RMSE (Median)** | **RMSE (Mode)** | **50% Coverage** | **95% Coverage** | **Factor 2 (Median)** | **Factor 2 (Mode)** |
| --- | --- | --- | --- | --- | --- | --- | --- | --- | --- | --- |
| Croatia/ Serbia/ Romania  *P.n.dalmatica/ P.n.nigra/ P.n.nigra/* | M | HRV-01+SCG-02 → ROU-01 | 0.7747 | 0.4343 | 1.575 | 1.218 | 0.44 | 0.936 | 0.706 | 0.768 |
|  | A | From HRV-01 to ROU-01 | 2.8935 | 2.8167 | 20.803 | 21.016 | 0.316 | 0.882 | 0.636 | 0.662 |
|  | D | Ancestral → HRV-01+SCG-02 | 0.5836 | 0.2408 | 0.994 | 0.703 | 0.484 | 0.934 | 0.796 | 0.882 |
| Italy/ Serbia/ Italy  *P.n.laricio/ P.n.nigra/ P.n.laricio* | M | ITA-01+ SCG-02 → ITA-02 | 2.144 | 1.4967 | 3.614 | 2.884 | 0.218 | 0.732 | 0.402 | 0.504 |
|  | A | From ITA-01 to ITA-02 | 0.155 | 0.1328 | 0.817 | 0.72 | 0.604 | 0.974 | 0.946 | 0.94 |
|  | D | Ancestral → ITA-01+ SCG-02 | 0.748 | 0.3134 | 1.277 | 0.837 | 0.416 | 0.904 | 0.708 | 0.838 |
| Francia Algeria Spain  *P.n.salzmannii/ P.n.salzmannii/ P.n.salzmannii/* | M | FRA-01+DZA-01 → ESP-01 | 0.9521 | 0.5564 | 1.906 | 1.415 | 0.404 | 0.914 | 0.664 | 0.692 |
|  | A | From DZA-01 to ESP-01 | 0.0493 | 0.0576 | 0.618 | 0.599 | 0.534 | 0.96 | 0.96 | 0.956 |
|  | D | Ancestral → FRA-01+DZA-01 | 0.5143 | 0.2083 | 0.915 | 0.665 | 0.496 | 0.934 | 0.81 | 0.87 |
| Serbia/ Ukraine/ Croatia  P.n.nigra/ P.n.pallasiana/ P.n.dalmatica | M | SCG-02 + CRIMEA-01 → HRV-02 | 3.5277 | 2.854 | 5.603 | 4.76 | 0.122 | 0.538 | 0.254 | 0.338 |
|  | A | From SCG-02 to HRV-02 | -0.0596 | -0.0742 | 0.576 | 0.586 | 0.368 | 0.938 | 0.966 | 0.952 |
|  | D | Ancestral → SCG-02 + CRIMEA-01 | 1.7543 | 1.1509 | 2.586 | 1.866 | 0.17 | 0.662 | 0.416 | 0.584 |
| Cyprus/ Ukraine/ Turkey  *P.n.pallasiana/ P.n.pallasiana/ P.n.pallasiana* | M | CYP-01 + CRIMEA-01 → TUR-01 | 0.5062 | 0.2371 | 1.116 | 0.866 | 0.506 | 0.978 | 0.792 | 0.824 |
|  | A | from CYP to TUR-01 | 0.6967 | 0.6241 | 2.093 | 1.938 | 0.494 | 0.948 | 0.818 | 0.84 |
|  | D | Ancestral →CYP-01 + CRIMEA-01 | 0.2901 | 0.0479 | 0.584 | 0.419 | 0.586 | 0.982 | 0.91 | 0.926 |

P* demographic parameters. M for merging time (in generation), A for admixture rate, D for divergence time (in generations)

**Table S2.10:** Model checking for the scenarios of admixture

The Probability (SSsimulated<SSobserved) is given for each summary statistics. To avoid the risk of over-estimating the quality of the fit we used different summary statistics (7 in total) than the ones used during the inference steps (model discrimination and posterior estimation of parameters). The simulated summary statistics were obtained from 10,000 data sets that were simulated from the posterior distribution of parameters.

**a**: Population 1 of Cyprus (CYP-01 ,*P.n. pallasiana)* and population 1 of Crimea *(CRIMEA-01 , P.n. pallasiana)* came into contact to give rise to the population 1 of Turkey *(TUR-01 , P.n. pallasiana)*

**b**: Population 1 of France (FRA-01 ,*P.n. salzmannii)* and population 1 of Algeria *( DZA-01 , P.n. salzmannii))* came into contact to give rise to the population 1 of Spain *(ESP-01 , P.n. salzmannii)*

**c**: Population 2 of Serbia ( SCG-02 ,*P.n. nigra)* and population 1 of Croatia *( HRV-01 , P.n. dalmatica)* came into contact to give rise to the population 1 of Romania *(ROU-01 , P.n. nigra)*

**d**: Population 2 of Serbia ( SCG-02 ,*P.n. nigra)* and population 1 of Crimea *(CRIMEA-01 , P.n. pallasiana)* came into contact to give rise to the population 2 of Croatia *(HRV-02 , P.n. dalmatica)*

**e**: Population 1 of Italia (ITA-01 ,*P.n. laricio)* and population 2 of Serbia *(SCG-02 , P.n. pallasiana)* came into contact to give rise to the population 2 of Italia *(ITA-02 , P.n. laricio)*

| Summary statistics | **a**  obs | **a**  p.value | **b**  obs | **b**  p.value | **c**  obs | **c**  p.value | **d**  obs | **d**  p.value | **e**  obs | **e**  p.value |
| --- | --- | --- | --- | --- | --- | --- | --- | --- | --- | --- |
| NAL_1_1 | 7.4286 | 0.49 | 6.4286 | 0.312 | 6.4286 | 0.417 | 7.7857 | 0.3875 | 8.3571 | 0.501 |
| NAL_1_2 | 7.3571 | 0.4175 | 6.2143 | 0.438 | 7.7857 | 0.7735 | 7.3571 | 0.2745 | 7.7857 | 0.4205 |
| NAL_1_3 | 8.1429 | 0.5975 | 7.9286 | 0.767 | 7.4286 | 0.7615 | 7.1429 | 0.4485 | 8.5 | 0.6195 |
| H2P_1_1&2 | 0.788 | 0.2925 | 0.766 | 0.3575 | 0.7645 | 0.4245 | 0.7663 | 0.1475 | 0.7869 | 0.26 |
| H2P_1_1&3 | 0.7866 | 0.319 | 0.744 | 0.275 | 0.7553 | 0.408 | 0.7627 | 0.1415 | 0.7706 | 0.2155 |
| H2P_1_2&3 | 0.7688 | 0.2475 | 0.7541 | 0.328 | 0.74 | 0.3325 | 0.7755 | 0.2025 | 0.7776 | 0.2155 |
| HET_2_1 | 0.6444 | 0.6675 | 0.6364 | 0.8475 | 0.303 | 0.3125 | 0.5636 | 0.662 | 0.7576 | 0.8665 |
| HET_2_2 | 0.4394 | 0.3525 | 0.3556 | 0.4965 | 0.5636 | 0.5935 | 0.4394 | 0.524 | 0.5636 | 0.5505 |
| HET_2_3 | 0.7424 | 0.7735 | 0.5303 | 0.666 | 0.6818 | 0.8345 | 0.6 | 0.6915 | 0.6545 | 0.671 |
| N2P_2_1&2 | 6 | 0.778 | 3 | 0.5725 | 5 | 0.7835 | 3 | 0.522 | 6 | 0.84 |
| N2P_2_1&3 | 4 | 0.444 | 5 | 0.908 | 6 | 0.9275 | 4 | 0.69 | 6 | 0.8625 |
| N2P_2_2&3 | 7 | 0.888 | 4 | 0.795 | 4 | 0.652 | 4 | 0.6975 | 3 | 0.3605 |
| NSS_3_1 | 5.6429 | 0.1 | 5.2857 | 0.0925 | 6.5714 | 0.151 | 7.2143 | 0.068 | 6.0714 | 0.1325 |
| NSS_3_2 | 7.3571 | 0.1035 | 4.8571 | 0.089 | 7.2143 | 0.1605 | 7.3571 | 0.0685 | 7.2143 | 0.159 |
| NSS_3_3 | 6.2143 | 0.0955 | 5.5714 | 0.087 | 7.4286 | 0.161 | 6.2143 | 0.064 | 6.6429 | 0.1425 |
| NH2_3_1&2 | 7.6429 | 0.107 | 6.0714 | 0.093 | 7.4286 | 0.1605 | 8.2857 | 0.0765 | 7.6429 | 0.165 |
| NH2_3_1&3 | 7.2143 | 0.1025 | 6.3571 | 0.096 | 6.7857 | 0.148 | 7.7857 | 0.076 | 6.8571 | 0.146 |
| NH2_3_2&3 | 7.0714 | 0.1015 | 6.1429 | 0.093 | 8.2857 | 0.186 | 7.5 | 0.073 | 7.7143 | 0.175 |
| NSS_4_1 | 0 | 0.4765 | 0 | 0.468 | 0 | 0.4845 | 0 | 0.4785 | 0 | 0.4795 |
| NSS_4_2 | 0 | 0.4735 | 0 | 0.47 | 0 | 0.478 | 0 | 0.4845 | 0 | 0.482 |
| NSS_4_3 | 0 | 0.467 | 0 | 0.4675 | 0 | 0.4785 | 0 | 0.4815 | 0 | 0.48 |

The code used for the summary statistics is composed of three parts: the summary statistic (see above for the definitions), the type of marker (1,2,3,4 for nSSR, cpSSR, nuclear gene, organelle gene respectively), the population (1,2,3 respectively for CYP-01, CRIMEA-01, TUR-01 (case a), respectively for FRA-01, DZA-01, ESP-01 (case b) respectively for SCG-02, HRV-01, ROU-01 (case c) respectively for SCG-02, CRIMEA-01, HRV-02 (case d) and respectively for ITA-01, SCG-02, ITA-02 (case e).

*HET*: mean genic diversity within population, *NAL*: mean number of alleles within population, *H2P:* mean genic diversity between pairs of populations , *N2P*: mean number of alleles between pairs of populations, *NSS*: mean number of segregating sites within population, *NH2*: mean number of haplotypes between pairs of populations.

***Appendix S3: Correlations between genetic parameters and environmental variables***

**Table S3.1:** List and definition of the environmental variables (83) used to test for correlations with genetic parameters. Description of how the climate data were processed and the correlations carried out. .

**Table S3.2:** Environmental characteristics for the eighteen populations

**Table S3.3:** Correlations of Pearson

**Figure S3.1:** classification tree of each genetic variable.

**Table S3.1:** List and definition of the environmental variables (83) used to test for correlations with genetic parameters. Description of how the climate data were processed and the correlations carried out.

| **ENVIRONNEMENTAL VARIABLES** | **DEFINITION** |
| --- | --- |
| **19 standard bioclimatic variables from the WorldClim database (version 1.4, Hijmans et al., 2005)** | |
| Bio 1 | Annual Mean Temperature |
| Bio 2 | Mean Diurnal Range |
| Bio 3 | Isothermality |
| Bio 4 | Temperature Seasonality |
| Bio 5 | Max Temperature of Warmest Month |
| Bio 6 | Min Temperature of Coldest Month |
| Bio 7 | Temperature Annual Range |
| Bio 8 | Mean Temperature of Wettest Quarter |
| Bio 9 | Mean Temperature of Driest Quarter |
| Bio 10 | Mean Temperature of Warmest Quarter |
| Bio 11 | Mean Temperature of Coldest Quarter |
| Bio 12 | Annual Precipitation |
| Bio 13 | Precipitation of Wettest Month |
| Bio 14 | Precipitation of Driest Month |
| Bio 15 | Precipitation Seasonality |
| Bio 16 | Precipitation of Wettest Quarter |
| Bio 17 | Precipitation of Driest Quarter |
| Bio 18 | Precipitation of Warmest Quarter |
| Bio19 | Precipitation of Coldest Quarter |
| **Bioclimatic variables at four geological times** | |
| Present | Present at 30 arc-seconds resolution |
| MH | Mid Holocene, approx. 6,000 years before present, at 30 arc-seconds resolution |
| LGM | Last Glacial Maximum (LGM, approx. 22,000 years BP) 2.5 arc-minutes resolution |
| LIG | Last inter-glacial (LIG, approx. between 120,000 – 140,000 years BP), at 30 arc-seconds resolution |
| **Other environmental variables:** | |
| Long | Longitude |
| Lat | Latitude |
| **Custom-made environmental variables:** | |
| FragArea | Area in Km2of the black pine natural non-fragmentated stands (according to the Euforgen Network) around the sampled population .When the sampled population did not fall in any stands the value of 100 km2 was noted |
| FragDistance | Distance in Km between the sampled population and the nearest Euforgen area delimitation |
| Isolation_index | qualitative index of isolation: 1 when Area higher than 10000 km2; 2 when Area between 1000 and 10 000 km2; 3 when Area lower than 1000 km² and the closest distance lower than 100 km; 4 when the Area was less than 1000 km2 and the closest distance higher than 100 km |
| Isolation_binary | binary value of the Isolation_index: 0 if Isolation_index lower than 3, 1 if Isolation_index higher or equal to 3, |

Latitude, longitude and 19 standard bioclimatic variables were downloaded from the WorldClim database (version 1.4, Hijmans et al., 2005) were downloaded in (January 2016) for the present time at 30 arc-seconds resolution, the Mid Holocene (MH, approx. 6,000 years before present) at 30 arc-seconds resolution, the Last Glacial Maximum (LGM, approx. 22,000 years BP) at 2.5 arc-minutes resolution, and last inter-glacial (LIG, approx. between 120,000 – 140,000 years BP) at 30 arc-seconds resolution. Climate data were obtained from the mean of the different WorldClim scenarios using the *stack* function in R (R Core Team, 2015). We also added four custom-made variables representative of the population isolation.

Pearson correlations and partial correlations, *i.e.* the correlation of two variables while controlling for longitude and latitude (“*ppcor*” R package, Kim 2015), were computed between each environmental and genetic variables. Genetic variables were: expected heterozygosity (*H_E_*), observed heterozygosity (*H_O_*), number of alleles (*N_A_*), fixation index (*F_IS_*), linkage disequilibrium (LD), N_0_/N_1_ ratio, and Qmax, which is the highest proportion of membership Q observed among the 7 STRUCTURE clusters. The higher the proportion of Q for one of the clusters, the higher its genetic homogeneity will be.

The environmental variables were highly correlated according to Pearson correlation analyses. Therefore, classifications were performed with the conditional inference version of cforest (Hothorn et al., 2006b; Strobl et al., 2007; 2008) in the party package of Random Forest using the settings suggested for the construction of unbiased random forests by Strobl et al. (2007). The number of input variables randomly sampled as candidate variables at each node was *mtry*= 70, the number of trees to grow was 4000. To test the consistency in the variable importance scores, the classification was repeated five times using different starting points in a set of pseudo-random numbers (seed value).

The five most explicative variables were retained to build the classification tree of each genetic variable. For this, we used the *ctree* function from the package *partykit* (Hothorn et al., 2006a). Significance of the selected variables was obtained from Monte Carlo procedures (9999 resampling); the value of the test statistics that must be exceeded in order to implement a split was 0.99. We conducted all analyses in Rstudio (RStudio Team version 1.0.143, 2016).

Hothorn, T., Buhlmann, P., Dudoit, S., Molinaro, A. & van der Laan, M.J. (2006a). Survival Ensembles. Biostatistics 7: 355–373.

Hothorn, T., Hornik, K. & Zeileis, A. (2006b). Unbiased Recursive Partitioning: A Conditional Inference Framework. J. Comput. Graph. Stat. 15: 651–674.

Kim, S. (2015). ppcor: An R Package for a Fast Calculation to Semi-partial Correlation Coefficients. Communications for Statistical Applications and Methods, 22(6): 665-674.

R Core Team, (2015) R: A language and environment for statistical computing. R Foundation for Statistical Computing, Vienna, Austria. URL <https://www.R-project.org/>.

RStudio Team (2016). RStudio: Integrated Development for R. RStudio, Inc., Boston, MA URL <http://www.rstudio.com/>.

Strobl, C., Boulesteix, A.L., Kneib, T., Augustin, T. & Zeileis A. (2008). Conditional variable importance for random forests. BMC Bioinformatics 9: 1–11. https://doi.org/10.1186/1471-2105-9-307

Strobl, C., Boulesteix, A.-L. Zeileis, A. & Hothorn, T. (2007). Bias in Random Forest Variable Importance Measures: Illustrations, Sources and a Solution. BMC Bioinformatics 8: 25.

**Table S3.2:** Environmental characteristics for the eighteen populations

| **Population** | **DZA-01** | **MAR-01** | **ESP-01** | **FRA-01** | **AUT-01** | **ITA-01** | **ITA-02** | **FRA-02** | **HRV-01** | **HRV-02** | **SCG-02** | **SCG-01** | **ROU-01** | **CRIMEA-01** | **TUR-01** | **TUR-02** | **CYP-01** | **CRIMEA-02** |
| --- | --- | --- | --- | --- | --- | --- | --- | --- | --- | --- | --- | --- | --- | --- | --- | --- | --- | --- |
| Long | 4.1 | -4.0 | -2.9 | 3.6 | 16.2 | 16.0 | 16.3 | 9.1 | 17.6 | 17.2 | 19.6 | 20.5 | 22.4 | 34.1 | 28.4 | 28.6 | 32.8 | 34.3 |
| Lat | 36.5 | 35.0 | 37.9 | 43.8 | 47.8 | 38.2 | 39.3 | 41.9 | 42.9 | 43.4 | 43.8 | 43.5 | 44.9 | 44.4 | 37.3 | 39.4 | 35.0 | 44.6 |
| FragArea | 1148.0 | 100.0 | 8434.0 | 100.0 | 2217.0 | 1446.0 | 2577.0 | 3113.0 | 100.0 | 86624 | 86624 | 6185.0 | 100.0 | 10363 | 100.0 | 33166 | 100.0 | 10363 |
| FragDistance | 0.0 | 276.0 | 0.0 | 179.0 | 0.0 | 0.0 | 0.0 | 0.0 | 57.0 | 0.0 | 0.0 | 0.0 | 215.0 | 0.0 | 361.0 | 0.0 | 172.0 | 0.0 |
| Isolation_index | 2.0 | 4.0 | 2.0 | 4.0 | 2.0 | 2.0 | 2.0 | 2.0 | 3.0 | 1.0 | 1.0 | 2.0 | 4.0 | 1.0 | 4.0 | 1.0 | 4.0 | 1.0 |
| Isolation_binary | 0.0 | 1.0 | 0.0 | 1.0 | 0.0 | 0.0 | 0.0 | 0.0 | 1.0 | 0.0 | 0.0 | 0.0 | 1.0 | 0.0 | 1.0 | 0.0 | 1.0 | 0.0 |
| bio1_Present | 12.8 | 16.7 | 11.6 | 11.7 | 9.0 | 11.2 | 13.1 | 9.2 | 14.6 | 13.9 | 8.5 | 9.2 | 7.4 | 10.0 | 10.5 | 9.4 | 12.5 | 11.9 |
| bio2_Present | 10.4 | 10.7 | 12.1 | 9.9 | 9.6 | 6.5 | 6.3 | 5.0 | 8.2 | 8.7 | 9.1 | 9.4 | 8.4 | 8.6 | 11.2 | 10.4 | 8.5 | 8.6 |
| bio3_Present | 33.0 | 40.0 | 37.0 | 37.0 | 32.0 | 29.0 | 28.0 | 24.0 | 32.0 | 31.0 | 31.0 | 31.0 | 29.0 | 29.0 | 36.0 | 36.0 | 31.0 | 29.0 |
| bio4_Present | 6971.0 | 5148.0 | 6664.0 | 5756.0 | 7263.0 | 5663.0 | 5618.0 | 5376.0 | 6148.0 | 6570.0 | 6850.0 | 7089.0 | 7253.0 | 7205.0 | 7121.0 | 6500.0 | 6468.0 | 7285.0 |
| bio5_Present | 30.8 | 31.1 | 30.6 | 26.4 | 25.1 | 23.8 | 25.5 | 20.6 | 28.9 | 29.1 | 23.3 | 24.7 | 22.5 | 25.5 | 28.0 | 25.2 | 27.1 | 27.5 |
| bio6_Present | 0.2 | 4.9 | -1.4 | -0.1 | -4.9 | 1.5 | 3.6 | 0.4 | 3.4 | 1.8 | -5.4 | -4.9 | -5.9 | -3.4 | -2.9 | -3.2 | 0.5 | -1.5 |
| bio7_Present | 30.6 | 26.2 | 32.0 | 26.5 | 30.0 | 22.3 | 21.9 | 20.2 | 25.5 | 27.3 | 28.7 | 29.6 | 28.4 | 28.9 | 30.9 | 28.4 | 26.6 | 29.0 |
| bio8_Present | 5.8 | 10.5 | 9.4 | 12.5 | 18.1 | 6.1 | 10.7 | 7.0 | 8.3 | 6.9 | 15.2 | 16.1 | 14.8 | 3.0 | 1.7 | 1.2 | 4.4 | 4.8 |
| bio9_Present | 22.0 | 23.5 | 20.7 | 19.2 | 0.9 | 18.5 | 20.4 | 16.2 | 22.6 | 22.4 | 0.8 | 1.3 | -1.0 | 8.5 | 19.7 | 17.5 | 20.6 | 10.3 |
| bio10_Present | 22.4 | 23.7 | 20.7 | 19.2 | 18.1 | 18.6 | 20.5 | 16.5 | 22.6 | 22.4 | 17.0 | 17.9 | 16.4 | 19.4 | 19.8 | 17.7 | 20.9 | 21.4 |
| bio11_Present | 4.6 | 10.5 | 3.8 | 4.5 | -0.6 | 4.5 | 6.4 | 3.0 | 7.1 | 5.6 | -0.7 | -0.3 | -2.1 | 1.0 | 1.7 | 1.2 | 4.4 | 2.9 |
| bio12_Present | 947.0 | 394.0 | 595.0 | 765.0 | 691.0 | 853.0 | 920.0 | 837.0 | 1204.0 | 1118.0 | 945.0 | 869.0 | 822.0 | 636.0 | 942.0 | 920.0 | 976.0 | 561.0 |
| bio13_Present | 135.0 | 61.0 | 75.0 | 89.0 | 88.0 | 120.0 | 134.0 | 108.0 | 167.0 | 151.0 | 100.0 | 96.0 | 121.0 | 83.0 | 190.0 | 161.0 | 219.0 | 74.0 |
| bio14_Present | 7.0 | 1.0 | 14.0 | 36.0 | 32.0 | 20.0 | 18.0 | 19.0 | 38.0 | 46.0 | 60.0 | 56.0 | 48.0 | 40.0 | 12.0 | 19.0 | 7.0 | 34.0 |
| bio15_Present | 57.0 | 64.0 | 40.0 | 20.0 | 33.0 | 50.0 | 50.0 | 39.0 | 40.0 | 34.0 | 16.0 | 15.0 | 33.0 | 23.0 | 76.0 | 57.0 | 92.0 | 24.0 |
| bio16_Present | 395.0 | 169.0 | 207.0 | 229.0 | 256.0 | 343.0 | 371.0 | 305.0 | 461.0 | 411.0 | 290.0 | 263.0 | 310.0 | 212.0 | 503.0 | 407.0 | 589.0 | 189.0 |
| bio17_Present | 48.0 | 13.0 | 61.0 | 148.0 | 111.0 | 73.0 | 79.0 | 92.0 | 147.0 | 169.0 | 187.0 | 179.0 | 147.0 | 132.0 | 44.0 | 70.0 | 27.0 | 115.0 |
| bio18_Present | 64.0 | 18.0 | 61.0 | 148.0 | 256.0 | 104.0 | 105.0 | 108.0 | 147.0 | 169.0 | 265.0 | 234.0 | 291.0 | 151.0 | 53.0 | 81.0 | 33.0 | 132.0 |
| bio19_Present | 367.0 | 169.0 | 178.0 | 195.0 | 111.0 | 284.0 | 343.0 | 259.0 | 411.0 | 362.0 | 204.0 | 191.0 | 158.0 | 210.0 | 503.0 | 407.0 | 589.0 | 187.0 |
| bio1_LGM | 8.1 | 12.1 | 7.4 | 6.8 | 0.0 | 7.6 | 6.4 | 4.3 | 8.0 | 6.4 | 1.7 | 2.5 | 1.5 | 3.0 | 5.5 | 4.7 | 10.1 | 3.4 |
| bio2_LGM | 10.5 | 10.3 | 12.3 | 10.9 | 11.0 | 8.3 | 7.2 | 6.4 | 10.6 | 11.2 | 10.4 | 10.5 | 9.8 | 10.8 | 12.4 | 12.8 | 9.3 | 10.9 |
| bio3_LGM | 39.7 | 48.3 | 43.7 | 41.7 | 31.0 | 36.0 | 33.0 | 31.3 | 35.3 | 34.3 | 32.3 | 32.0 | 28.7 | 30.0 | 37.3 | 36.7 | 33.7 | 29.7 |
| bio4_LGM | 5425.0 | 3550.0 | 5173.7 | 5367.7 | 8768.3 | 5284.0 | 5284.7 | 4847.0 | 6752.7 | 7413.0 | 7622.7 | 7763.7 | 8445.7 | 8611.0 | 7146.3 | 7426.3 | 6373.3 | 8794.3 |
| bio5_LGM | 23.7 | 23.8 | 24.0 | 20.5 | 17.3 | 20.4 | 18.4 | 15.6 | 23.7 | 23.3 | 16.9 | 18.2 | 18.5 | 21.1 | 24.2 | 23.5 | 25.3 | 21.9 |
| bio6_LGM | -2.6 | 2.6 | -3.6 | -5.4 | -17.7 | -2.4 | -3.1 | -4.3 | -5.5 | -8.6 | -14.6 | -13.8 | -14.9 | -14.4 | -8.9 | -10.6 | -1.8 | -14.2 |
| bio7_LGM | 26.3 | 21.2 | 27.6 | 25.9 | 35.0 | 22.9 | 21.5 | 19.9 | 29.2 | 31.8 | 31.5 | 32.0 | 33.5 | 35.5 | 33.1 | 34.1 | 27.1 | 36.1 |
| bio8_LGM | 2.3 | 7.8 | 3.5 | 4.0 | 10.1 | 1.9 | 0.6 | -1.2 | 0.9 | 4.4 | 6.4 | 6.2 | 9.7 | -7.7 | -3.1 | -4.0 | 2.2 | -7.5 |
| bio9_LGM | 15.4 | 16.9 | 14.5 | 13.4 | -11.6 | 13.5 | 12.4 | 10.2 | 15.3 | 14.4 | -6.2 | -0.5 | -7.8 | 9.1 | 14.6 | 13.8 | 18.4 | 9.7 |
| bio10_LGM | 15.5 | 16.9 | 14.5 | 13.6 | 10.4 | 14.6 | 13.3 | 10.7 | 16.5 | 15.4 | 10.6 | 11.5 | 11.5 | 13.7 | 14.9 | 14.2 | 18.6 | 14.3 |
| bio11_LGM | 1.8 | 7.8 | 1.4 | -0.2 | -11.6 | 1.2 | -0.1 | -1.5 | -0.7 | -3.4 | -8.8 | -8.2 | -9.9 | -8.4 | -3.6 | -4.9 | 2.2 | -8.2 |
| bio12_LGM | 1159.3 | 568.0 | 679.0 | 745.0 | 668.7 | 806.3 | 840.0 | 876.3 | 1207.3 | 1185.7 | 974.7 | 898.0 | 753.7 | 587.7 | 977.3 | 888.3 | 953.7 | 546.0 |
| bio13_LGM | 180.0 | 103.0 | 96.0 | 87.7 | 89.7 | 101.0 | 103.3 | 107.3 | 146.3 | 146.0 | 123.0 | 105.3 | 101.7 | 83.0 | 191.3 | 151.3 | 227.3 | 76.0 |
| bio14_LGM | 11.7 | 1.0 | 11.0 | 34.7 | 34.0 | 28.3 | 35.0 | 38.3 | 43.0 | 52.3 | 56.7 | 52.3 | 42.7 | 26.7 | 12.3 | 13.0 | 5.7 | 25.0 |
| bio15_LGM | 62.0 | 76.3 | 52.0 | 25.3 | 32.3 | 36.7 | 31.3 | 31.7 | 32.7 | 28.7 | 22.7 | 21.3 | 29.3 | 37.0 | 69.0 | 56.7 | 93.7 | 35.3 |
| bio16_LGM | 512.0 | 280.3 | 274.0 | 235.3 | 247.0 | 287.3 | 291.7 | 299.3 | 413.3 | 399.7 | 312.7 | 279.0 | 260.7 | 219.7 | 484.0 | 372.7 | 577.3 | 200.7 |
| bio17_LGM | 63.7 | 12.0 | 54.3 | 135.0 | 113.0 | 104.3 | 127.0 | 135.0 | 165.7 | 186.0 | 190.0 | 173.3 | 136.0 | 93.0 | 49.3 | 56.7 | 23.7 | 86.7 |
| bio18_LGM | 67.3 | 12.0 | 54.3 | 136.7 | 244.3 | 109.7 | 132.0 | 146.3 | 188.0 | 227.3 | 264.3 | 224.0 | 242.7 | 106.0 | 56.3 | 69.3 | 25.0 | 99.0 |
| bio19_LGM | 459.7 | 280.3 | 253.0 | 214.0 | 113.0 | 275.7 | 278.7 | 297.3 | 384.7 | 354.0 | 199.0 | 187.3 | 143.3 | 214.7 | 470.3 | 350.0 | 577.3 | 194.7 |
| bio1_MH | 12.7 | 16.6 | 11.4 | 11.9 | 9.5 | 11.4 | 13.3 | 9.4 | 15.1 | 14.4 | 9.0 | 9.7 | 8.1 | 10.6 | 10.8 | 9.9 | 12.7 | 12.5 |
| bio2_MH | 10.5 | 10.3 | 12.3 | 10.9 | 11.0 | 8.3 | 7.2 | 6.4 | 10.6 | 11.2 | 10.4 | 10.5 | 9.8 | 10.8 | 12.4 | 12.8 | 9.3 | 10.9 |
| bio3_MH | 31.2 | 36.7 | 34.8 | 34.6 | 29.8 | 26.3 | 25.3 | 22.0 | 29.0 | 28.7 | 28.8 | 28.8 | 26.6 | 27.1 | 32.6 | 32.8 | 29.3 | 27.1 |
| bio4_MH | 7747.7 | 6046.4 | 7556.7 | 6538.7 | 8114.3 | 6258.4 | 6267.8 | 5987.0 | 7026.1 | 7464.7 | 7737.1 | 7981.1 | 8159.7 | 7952.6 | 7937.9 | 7415.4 | 6992.8 | 8063.2 |
| bio5_MH | 33.1 | 33.7 | 32.6 | 28.3 | 28.0 | 25.6 | 27.6 | 22.4 | 31.6 | 32.0 | 27.1 | 28.6 | 26.5 | 27.9 | 30.9 | 28.7 | 28.7 | 29.9 |
| bio6_MH | -0.2 | 4.3 | -1.8 | -0.3 | -4.7 | 1.3 | 3.5 | 0.2 | 3.2 | 1.7 | -5.4 | -4.9 | -5.7 | -3.4 | -3.1 | -3.5 | 0.2 | -1.5 |
| bio7_MH | 33.2 | 29.3 | 34.4 | 28.7 | 32.7 | 24.3 | 24.2 | 22.2 | 28.3 | 30.4 | 32.5 | 33.4 | 32.2 | 31.3 | 34.0 | 32.2 | 28.5 | 31.4 |
| bio8_MH | 5.3 | 11.3 | 7.7 | 10.4 | 18.4 | 8.0 | 10.2 | 6.6 | 10.3 | 9.8 | 14.5 | 15.0 | 14.7 | 2.6 | 1.5 | 1.5 | 4.1 | 4.4 |
| bio9_MH | 22.9 | 24.4 | 21.9 | 18.8 | -0.3 | 19.4 | 21.4 | 17.2 | 24.4 | 24.3 | 2.9 | 3.4 | 1.4 | 17.0 | 21.6 | 19.8 | 21.8 | 18.1 |
| bio10_MH | 23.8 | 25.3 | 22.3 | 20.7 | 20.2 | 20.0 | 22.0 | 17.8 | 24.6 | 24.3 | 19.0 | 20.0 | 18.7 | 21.2 | 21.6 | 19.8 | 21.8 | 23.2 |
| bio11_MH | 4.1 | 9.8 | 3.2 | 4.1 | -0.6 | 4.2 | 6.2 | 2.8 | 6.9 | 5.4 | -0.7 | -0.4 | -2.0 | 1.0 | 1.5 | 1.0 | 4.1 | 2.8 |
| bio12_MH | 1113.1 | 502.6 | 719.9 | 819.7 | 709.1 | 962.4 | 1036.3 | 940.1 | 1292.4 | 1196.2 | 1020.1 | 938.0 | 866.7 | 643.8 | 1057.9 | 997.1 | 1106.3 | 567.0 |
| bio13_MH | 162.0 | 82.6 | 108.0 | 96.2 | 97.6 | 134.7 | 144.3 | 120.8 | 179.4 | 162.7 | 125.8 | 114.7 | 138.8 | 89.2 | 204.8 | 164.7 | 246.2 | 79.0 |
| bio14_MH | 7.0 | 1.0 | 12.9 | 34.9 | 31.0 | 19.9 | 18.4 | 20.0 | 36.0 | 44.8 | 59.6 | 55.6 | 46.2 | 32.4 | 11.0 | 15.1 | 7.4 | 28.2 |
| bio15_MH | 56.1 | 66.7 | 47.2 | 24.3 | 35.2 | 50.1 | 49.7 | 40.4 | 40.0 | 33.8 | 23.3 | 22.8 | 37.6 | 29.9 | 72.7 | 55.7 | 90.2 | 31.0 |
| bio16_MH | 438.2 | 225.2 | 289.2 | 248.3 | 266.8 | 381.0 | 413.7 | 332.8 | 487.4 | 432.4 | 337.4 | 308.1 | 340.3 | 229.2 | 536.4 | 417.6 | 645.4 | 203.7 |
| bio17_MH | 55.8 | 15.7 | 66.3 | 151.8 | 108.0 | 74.8 | 83.4 | 97.4 | 150.7 | 172.4 | 192.8 | 182.7 | 148.8 | 117.0 | 43.3 | 64.7 | 29.2 | 102.8 |
| bio18_MH | 75.7 | 22.6 | 70.9 | 170.1 | 260.1 | 106.6 | 106.0 | 110.7 | 170.4 | 182.0 | 279.9 | 250.1 | 301.0 | 131.4 | 43.3 | 66.9 | 29.2 | 114.7 |
| bio19_MH | 404.4 | 198.7 | 202.9 | 201.1 | 109.3 | 324.2 | 354.3 | 300.1 | 425.2 | 372.1 | 204.4 | 190.7 | 156.6 | 209.1 | 536.4 | 417.1 | 645.4 | 181.0 |
| bio1_LIG | 11.9 | 16.0 | 10.7 | 11.3 | 8.0 | 11.7 | 13.6 | 9.2 | 14.3 | 13.4 | 8.0 | 8.8 | 6.5 | 10.0 | 10.5 | 9.5 | 12.1 | 11.9 |
| bio2_LIG | 9.6 | 10.3 | 11.4 | 9.9 | 11.3 | 7.1 | 6.7 | 5.6 | 9.0 | 9.6 | 10.1 | 10.4 | 9.6 | 9.5 | 11.8 | 10.7 | 9.1 | 9.5 |
| bio3_LIG | 26.0 | 32.0 | 29.0 | 30.0 | 25.0 | 26.0 | 24.0 | 21.0 | 26.0 | 26.0 | 26.0 | 26.0 | 22.0 | 24.0 | 31.0 | 29.0 | 27.0 | 24.0 |
| bio4_LIG | 9275 | 7095 | 9093 | 7827 | 10732 | 7210 | 7193 | 7018 | 8287 | 8842 | 9451 | 9711 | 10684 | 9377 | 9045 | 8559 | 8436 | 9497 |
| bio5_LIG | 35.1 | 35.0 | 35.7 | 31.5 | 30.5 | 27.9 | 29.6 | 24.8 | 33.7 | 33.9 | 28.5 | 30.1 | 28.7 | 30.9 | 33.0 | 30.8 | 30.9 | 33.0 |
| bio6_LIG | -1.1 | 3.6 | -3.2 | -1.5 | -13.7 | 0.7 | 2.7 | -0.8 | 0.3 | -1.8 | -10.3 | -9.7 | -14.0 | -7.5 | -4.5 | -5.0 | -1.8 | -5.8 |
| bio7_LIG | 36.2 | 31.4 | 38.9 | 33.0 | 44.2 | 27.2 | 26.9 | 25.6 | 33.4 | 35.7 | 38.8 | 39.8 | 42.7 | 38.4 | 37.5 | 35.8 | 32.7 | 38.8 |
| bio8_LIG | 2.3 | 8.1 | 4.3 | 5.7 | 21.1 | 5.3 | 9.9 | 5.4 | 9.4 | 8.0 | 13.8 | 15.0 | 19.4 | 0.9 | 1.6 | 1.1 | 2.1 | 2.7 |
| bio9_LIG | 25.3 | 26.4 | 24.3 | 22.6 | -4.0 | 19.2 | 21.3 | 19.2 | 26.0 | 25.7 | -2.4 | -1.9 | -5.7 | 3.2 | 23.1 | 21.5 | 20.7 | 17.9 |
| bio10_LIG | 25.3 | 26.4 | 24.3 | 22.6 | 21.9 | 21.9 | 23.8 | 19.2 | 26.0 | 25.7 | 20.7 | 21.9 | 20.6 | 22.9 | 23.1 | 21.5 | 23.8 | 25.0 |
| bio11_LIG | 1.5 | 8.1 | 1.0 | 2.6 | -6.0 | 3.7 | 5.6 | 1.6 | 4.6 | 2.8 | -4.0 | -3.5 | -7.3 | -1.3 | 0.1 | -0.5 | 2.1 | 0.5 |
| bio12_LIG | 1005.0 | 507.0 | 699.0 | 802.0 | 722.0 | 1026.0 | 1091.0 | 928.0 | 1234.0 | 1141.0 | 953.0 | 880.0 | 844.0 | 629.0 | 1044.0 | 1018.0 | 1069.0 | 556.0 |
| bio13_LIG | 145.0 | 87.0 | 97.0 | 98.0 | 106.0 | 139.0 | 156.0 | 126.0 | 181.0 | 166.0 | 122.0 | 119.0 | 128.0 | 85.0 | 196.0 | 179.0 | 229.0 | 78.0 |
| bio14_LIG | 2.0 | 0.0 | 10.0 | 33.0 | 24.0 | 13.0 | 16.0 | 19.0 | 31.0 | 38.0 | 53.0 | 51.0 | 40.0 | 35.0 | 12.0 | 23.0 | 12.0 | 29.0 |
| bio15_LIG | 57.0 | 61.0 | 45.0 | 30.0 | 41.0 | 52.0 | 54.0 | 43.0 | 44.0 | 38.0 | 24.0 | 27.0 | 38.0 | 35.0 | 72.0 | 61.0 | 82.0 | 37.0 |
| bio16_LIG | 411.0 | 218.0 | 244.0 | 262.0 | 278.0 | 410.0 | 449.0 | 347.0 | 490.0 | 437.0 | 286.0 | 264.0 | 328.0 | 249.0 | 529.0 | 477.0 | 600.0 | 226.0 |
| bio17_LIG | 43.0 | 29.0 | 58.0 | 116.0 | 86.0 | 78.0 | 83.0 | 89.0 | 137.0 | 159.0 | 165.0 | 156.0 | 123.0 | 120.0 | 66.0 | 74.0 | 64.0 | 103.0 |
| bio18_LIG | 43.0 | 29.0 | 58.0 | 116.0 | 260.0 | 100.0 | 98.0 | 89.0 | 137.0 | 159.0 | 243.0 | 210.0 | 281.0 | 125.0 | 66.0 | 74.0 | 73.0 | 107.0 |
| bio19_LIG | 395.0 | 218.0 | 227.0 | 211.0 | 99.0 | 348.0 | 413.0 | 317.0 | 427.0 | 373.0 | 206.0 | 194.0 | 153.0 | 209.0 | 528.0 | 443.0 | 600.0 | 187.0 |

**Table S3.3:** Pearson correlations with a False Discovery Rate lower than 0.1

| **VAR1** | **VAR2** | **ddl** | **Pearson correlation** | **p.value** | **FDR** |
| --- | --- | --- | --- | --- | --- |
| He_Genes | bio4_LGM | 15 | 0.77 | 0.0003 | 0.061 |
| He_Genes | bio6_LGM | 15 | -0.77 | 0.0003 | 0.062 |
| He_Genes | bio7_LGM | 15 | 0.72 | 0.0010 | 0.086 |
| He_Genes | bio11_LGM | 15 | -0.73 | 0.0009 | 0.083 |
| Na_Genes | bio4_LGM | 15 | 0.76 | 0.0003 | 0.063 |
| Na_Genes | bio6_LGM | 15 | -0.79 | 0.0002 | 0.051 |
| Na_Genes | bio11_LGM | 15 | -0.77 | 0.0003 | 0.059 |
| Na_Genes | bio18_LIG | 15 | 0.73 | 0.0009 | 0.084 |
| N_current_N_past_ | bio10_Present | 5 | -0.94 | 0.0016 | 0.098 |
| N_current_N_past_ | bio10_MH | 5 | -0.95 | 0.0008 | 0.082 |
| N_current_N_past_ | bio10_LIG | 5 | -0.95 | 0.0009 | 0.084 |

**Figure S3.1:** Classification tree of each genetic variable. See Table S3.1 for the list and definition of the environmental variables and time-periods used to test for correlations with genetic parameters.

In a first analysis, classifications were performed with the conditional inference version of *cforest* (Hothorn et al. 2006b; Strobl et al. 2007, 2008) in the *party* package of Random Forest using the settings suggested for the construction of unbiased random forests by Strobl et al. (2007). The number of input variables randomly sampled as candidates at each node was mtry= 70, the number of trees to grow was 4000. To test the consistency in the variable importance scores, the classification was repeated five times using different seed value.

The five most explicative variables were used to build the classification tree of each genetic variable presented in the figure S3.1. For this, we used the *ctree* function from the package *partykit* (Hothorn *et al.* 2006a). Significance of the selected variables was obtained from Monte Carlo procedures (9999 resampling); the value of the test statistics that must be exceeded in order to implement a split was 0.99. We conducted all analyses in Rstudio (RStudio Team 2016, version 1.0.143).

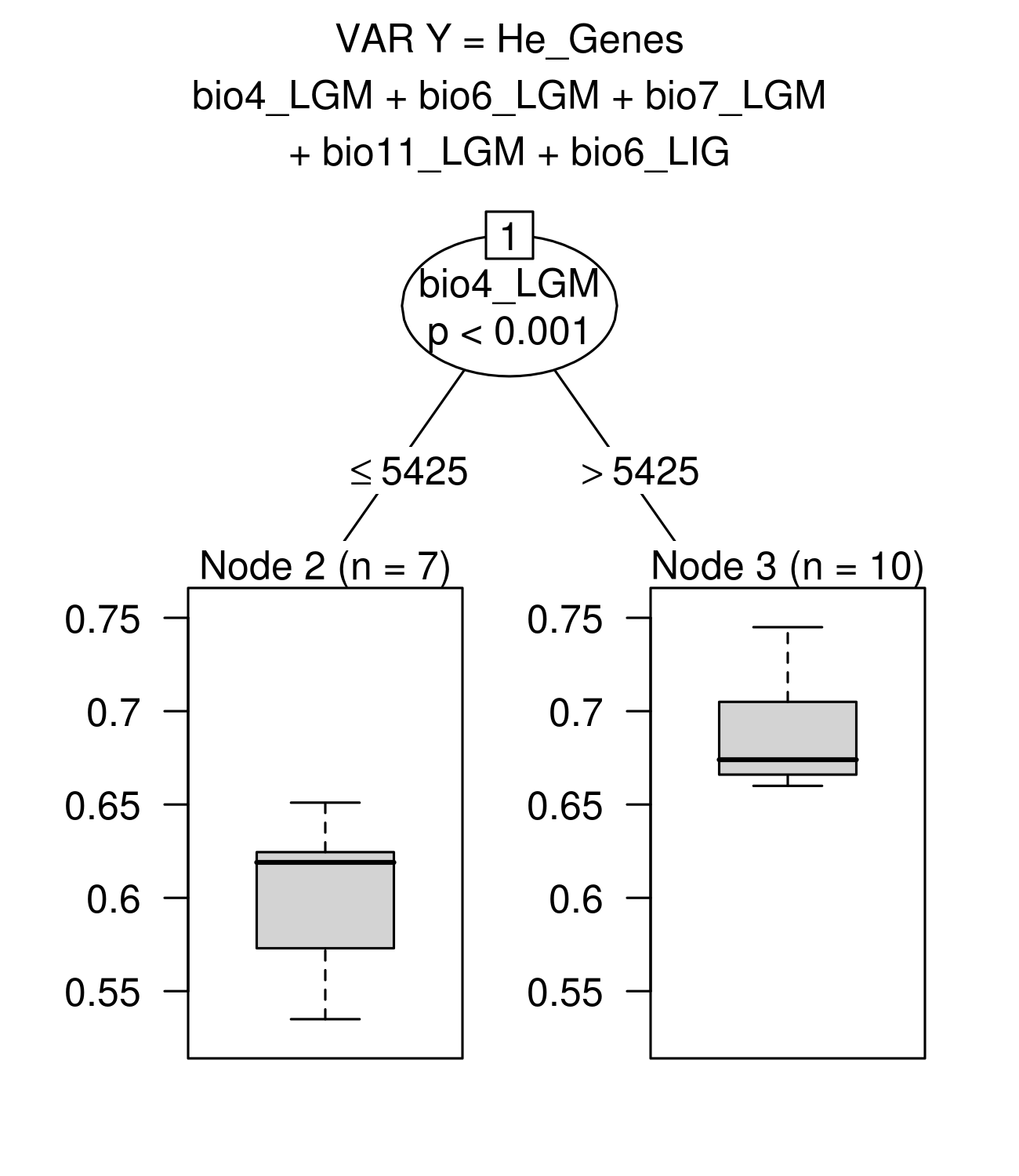

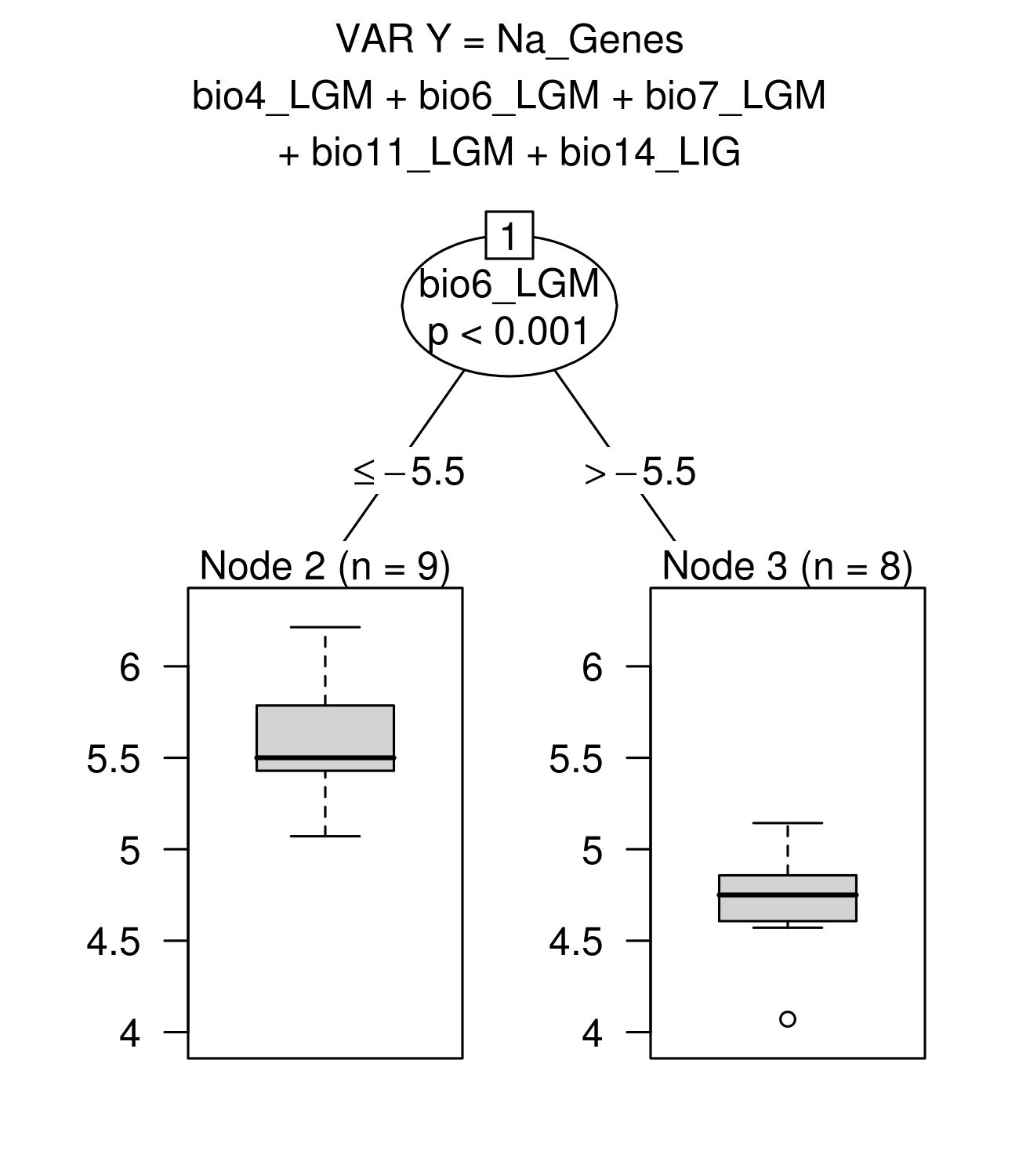

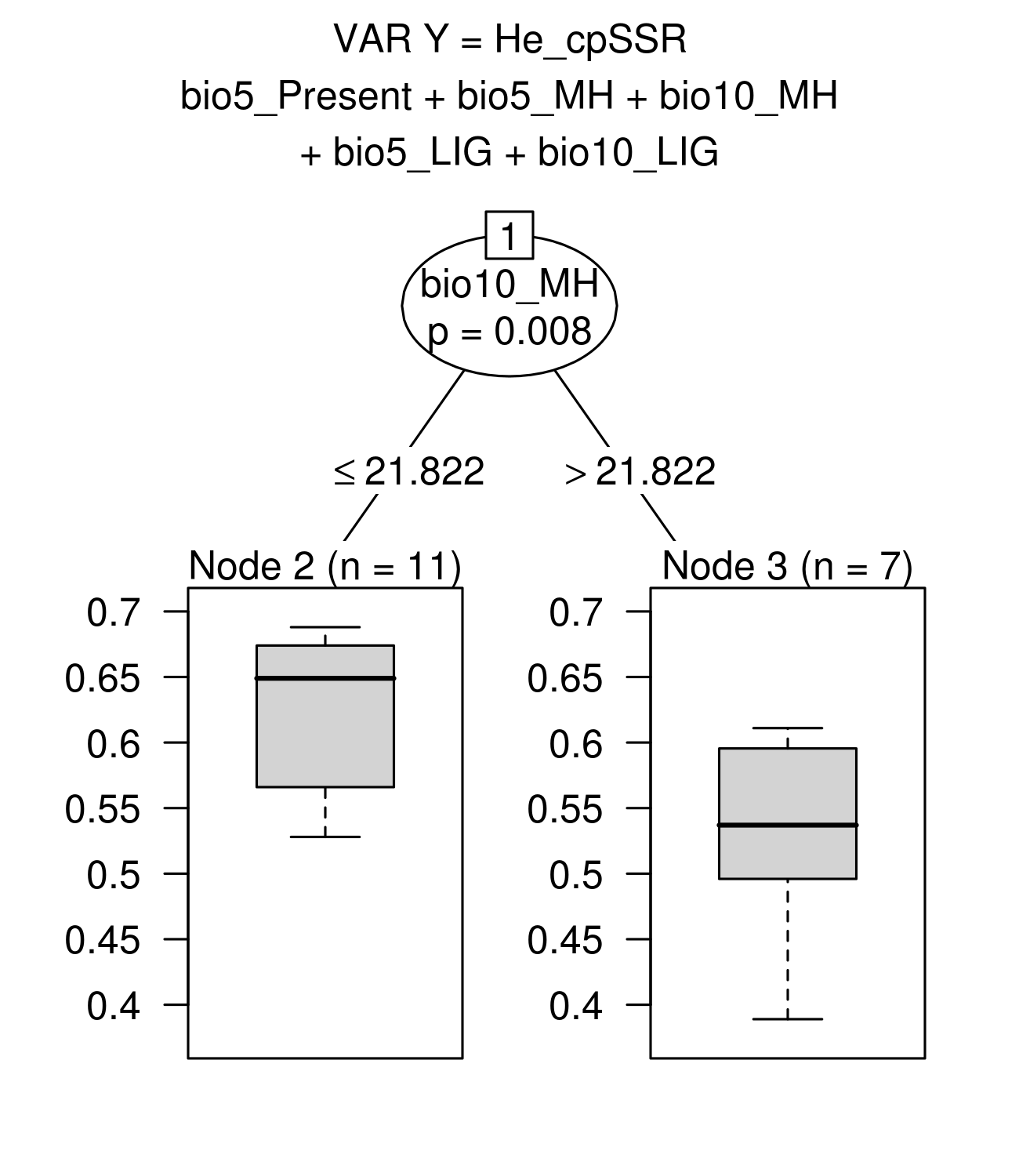

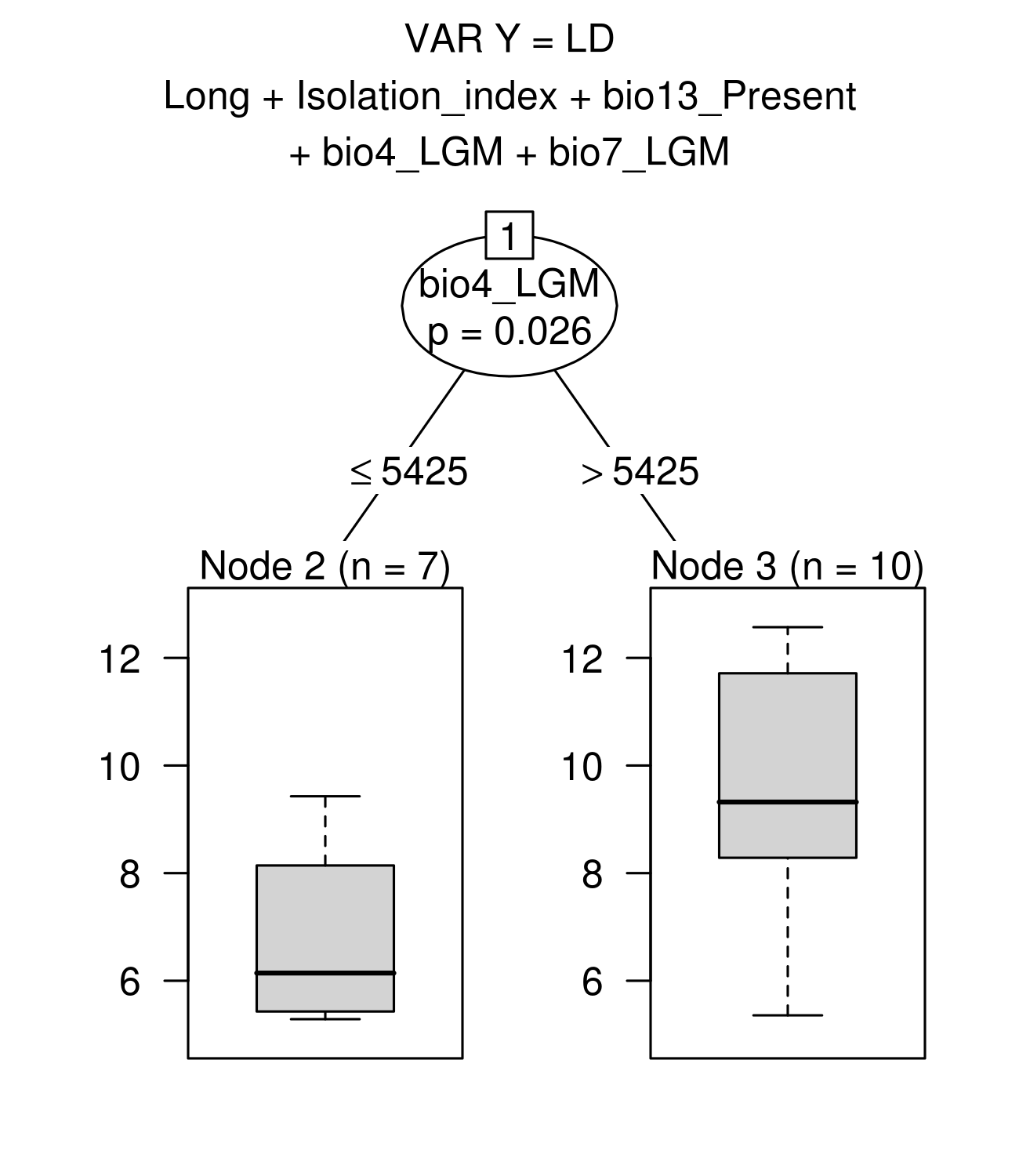

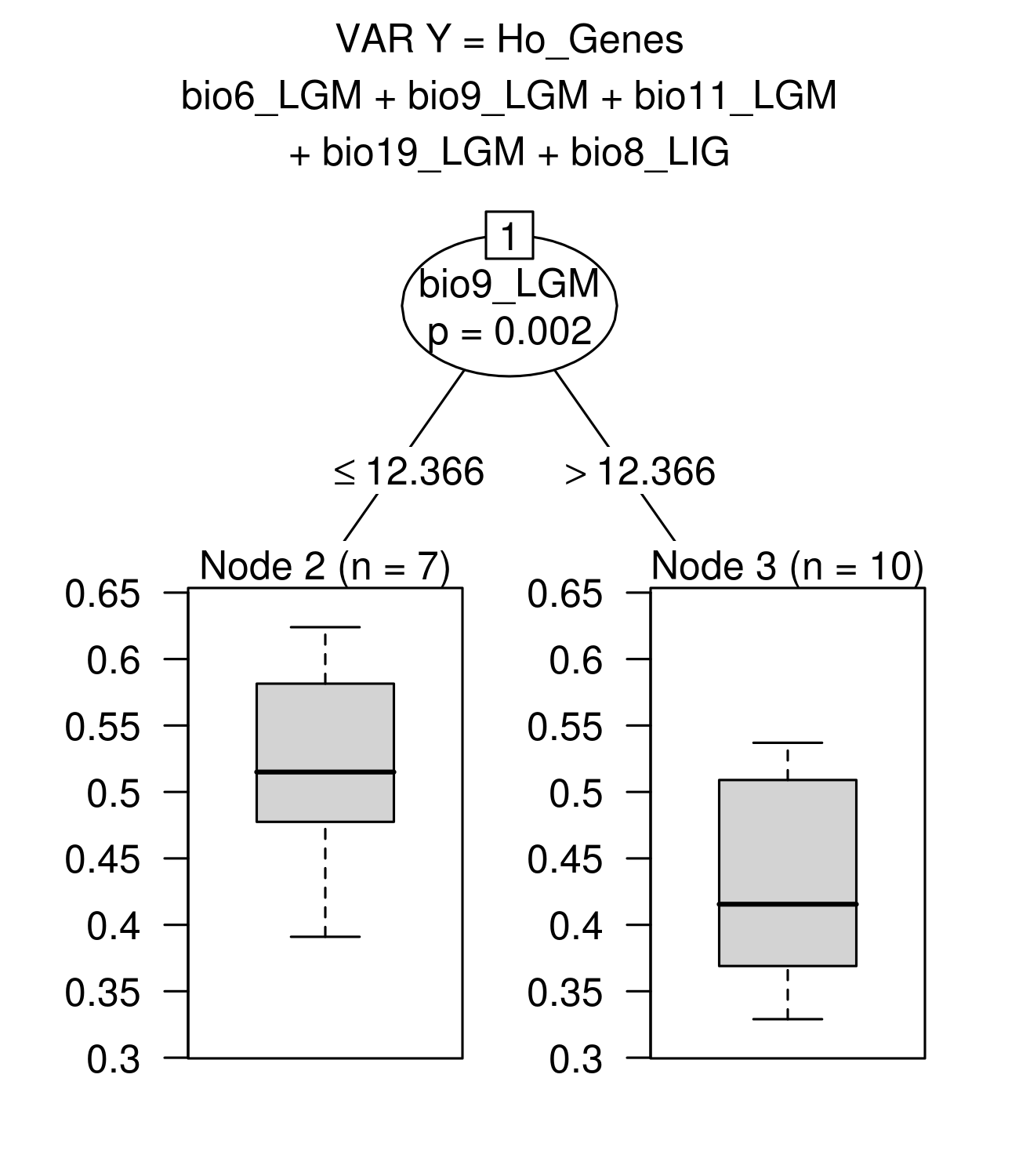

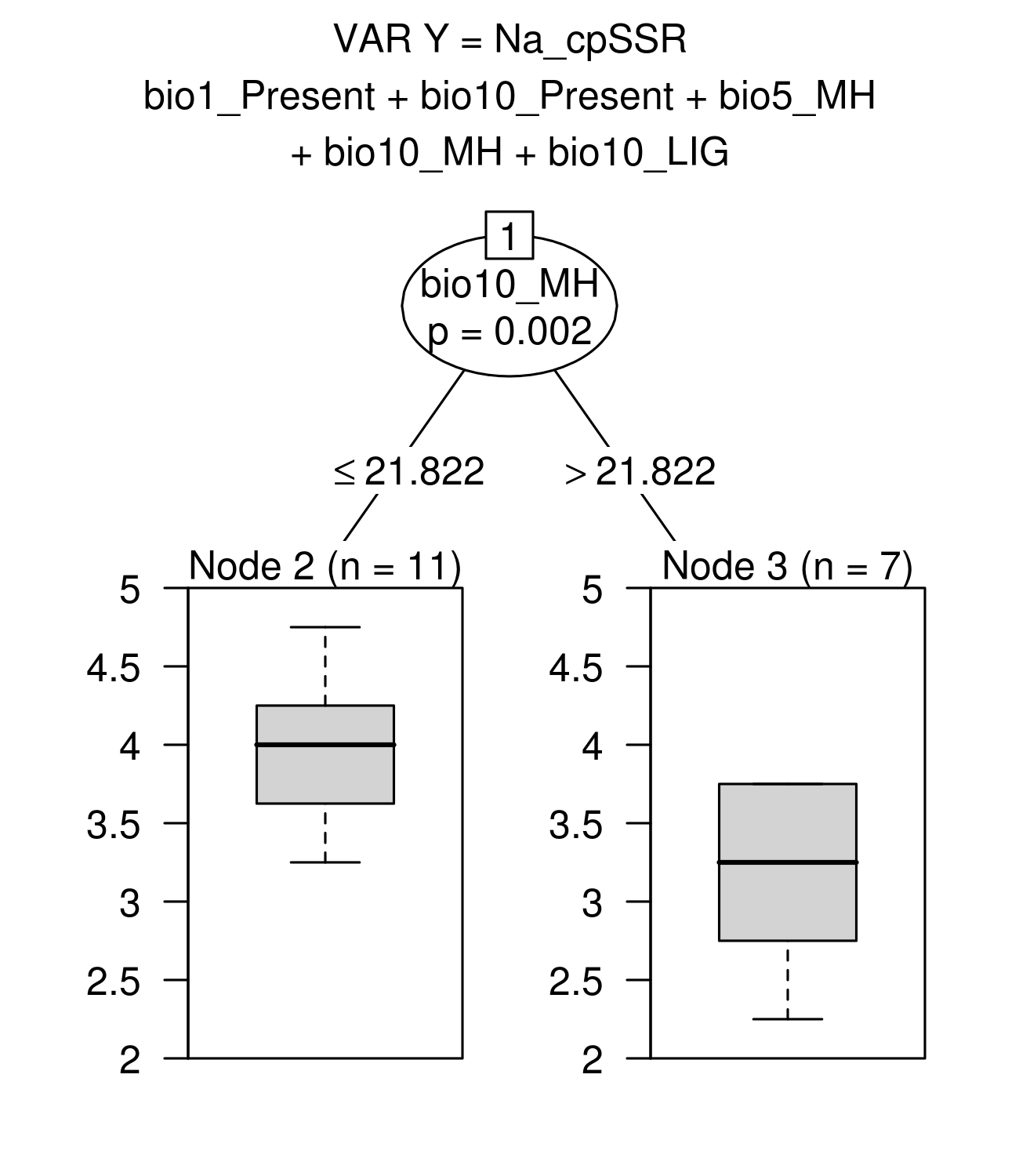

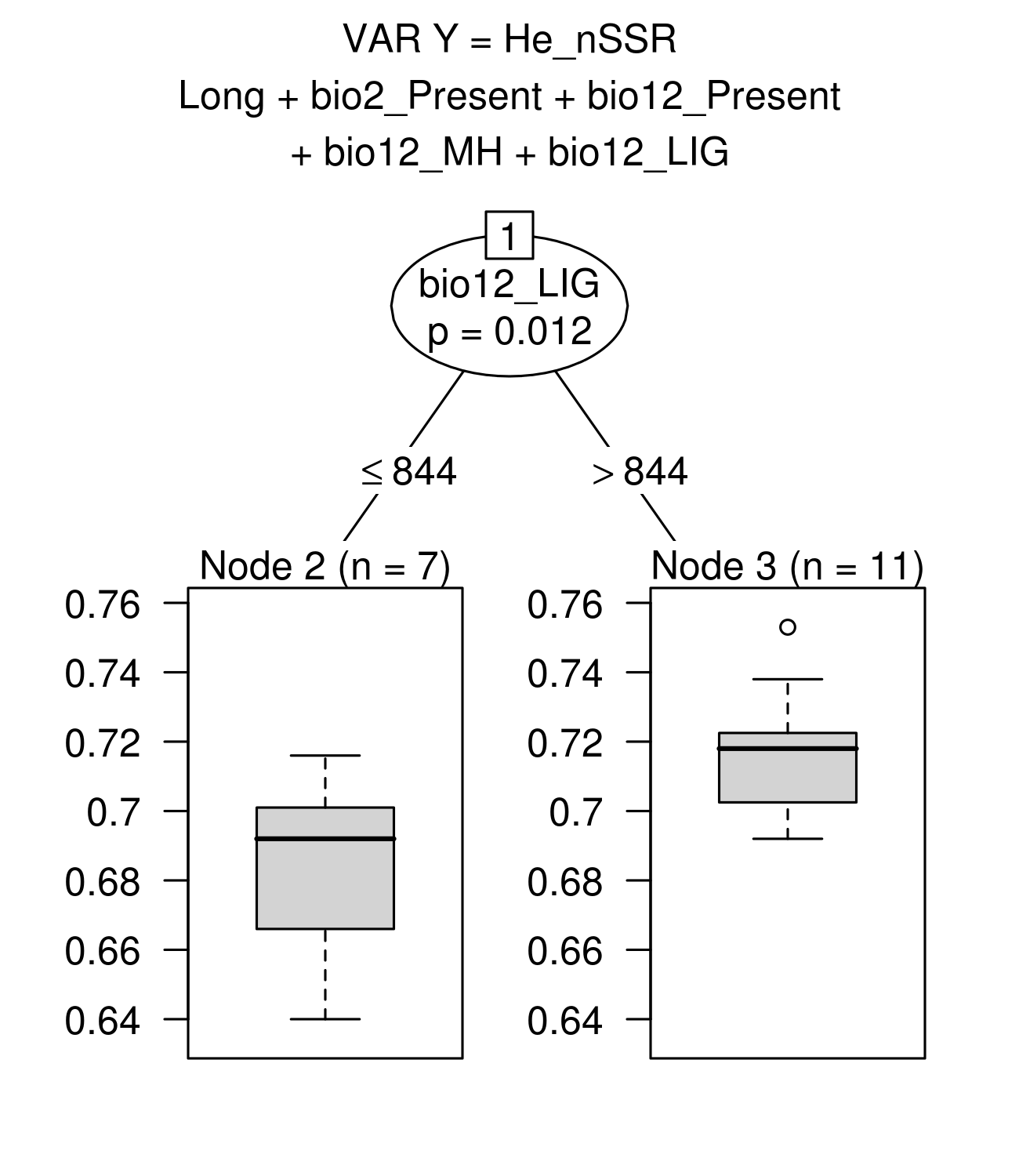

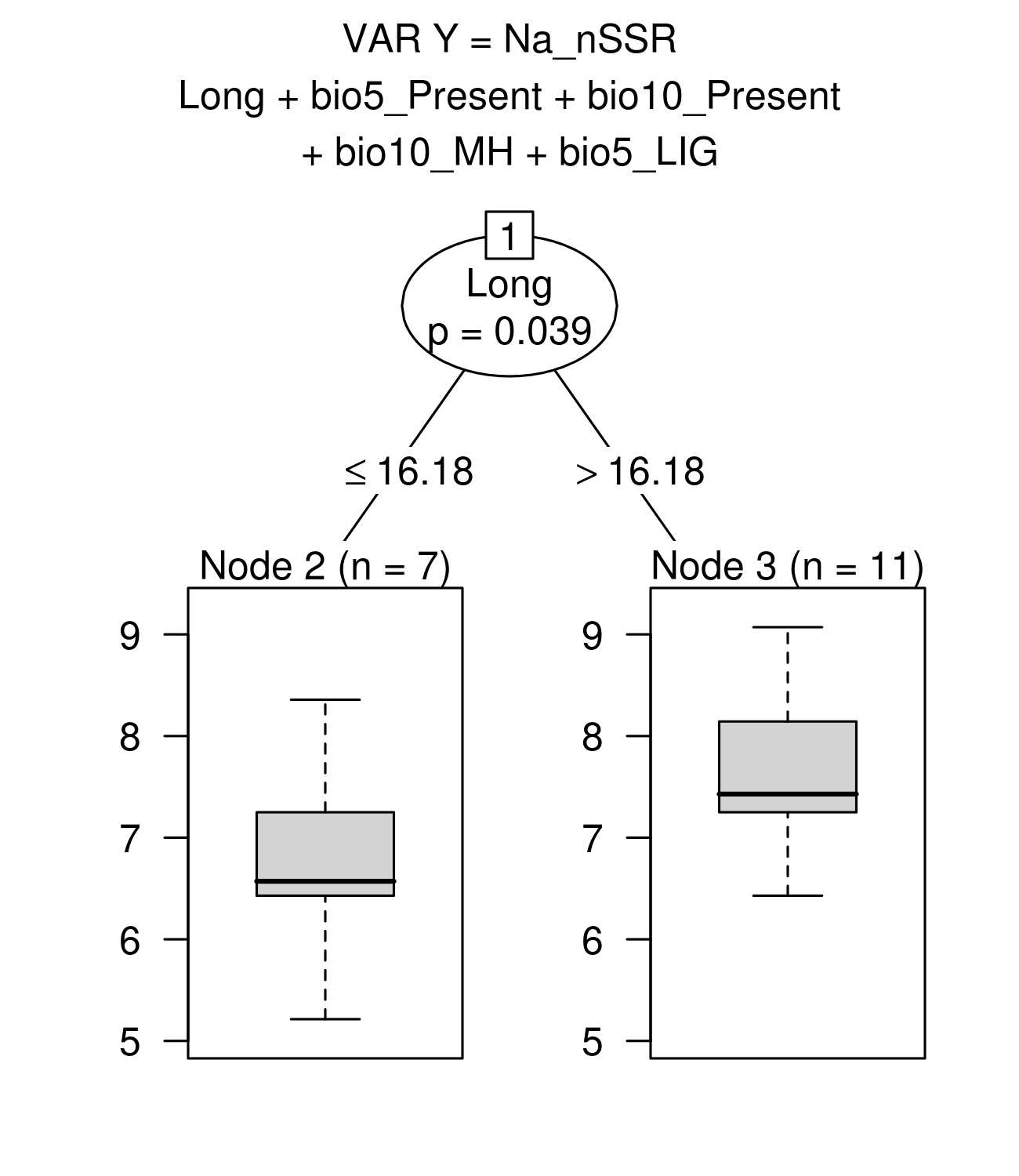

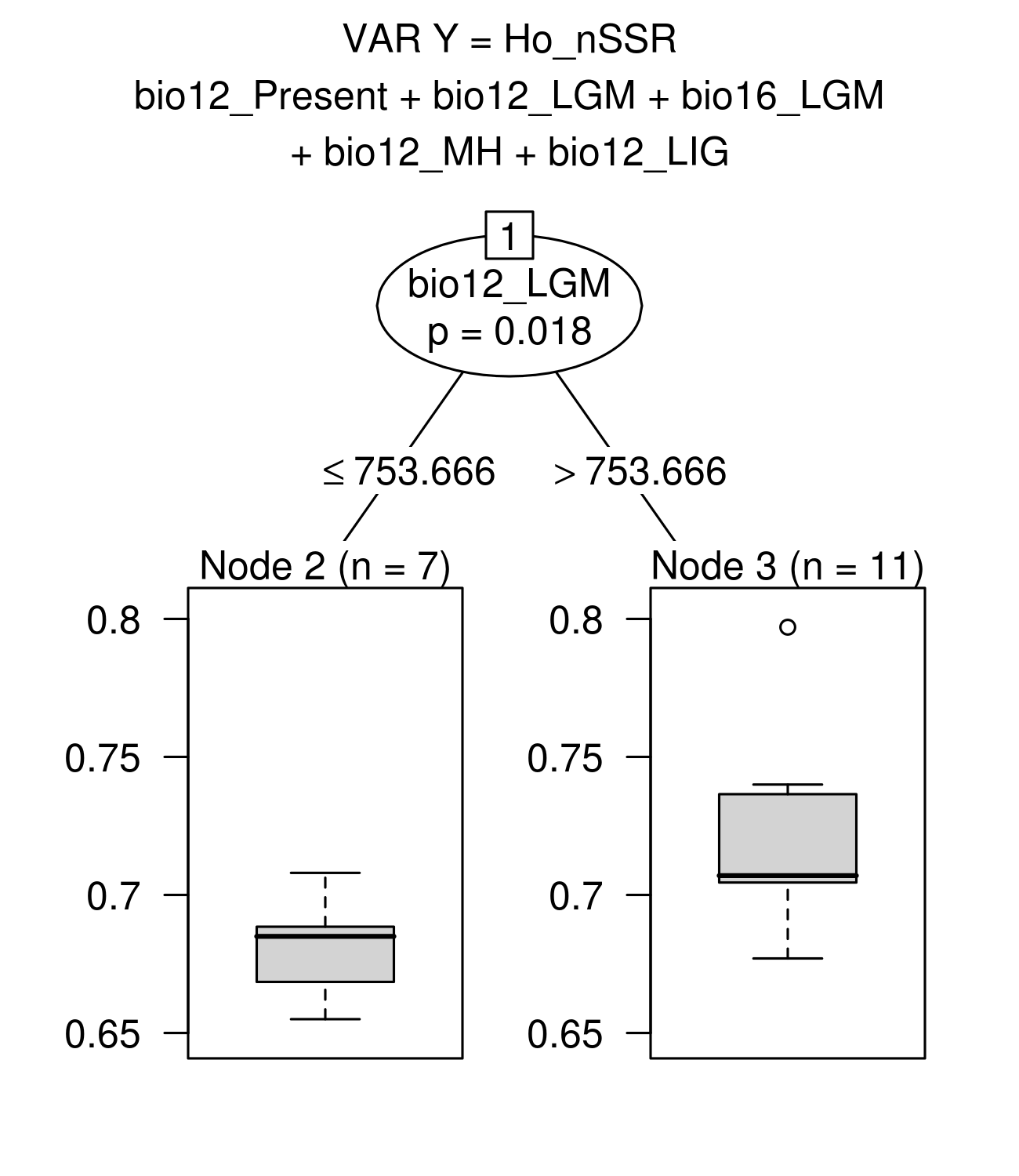

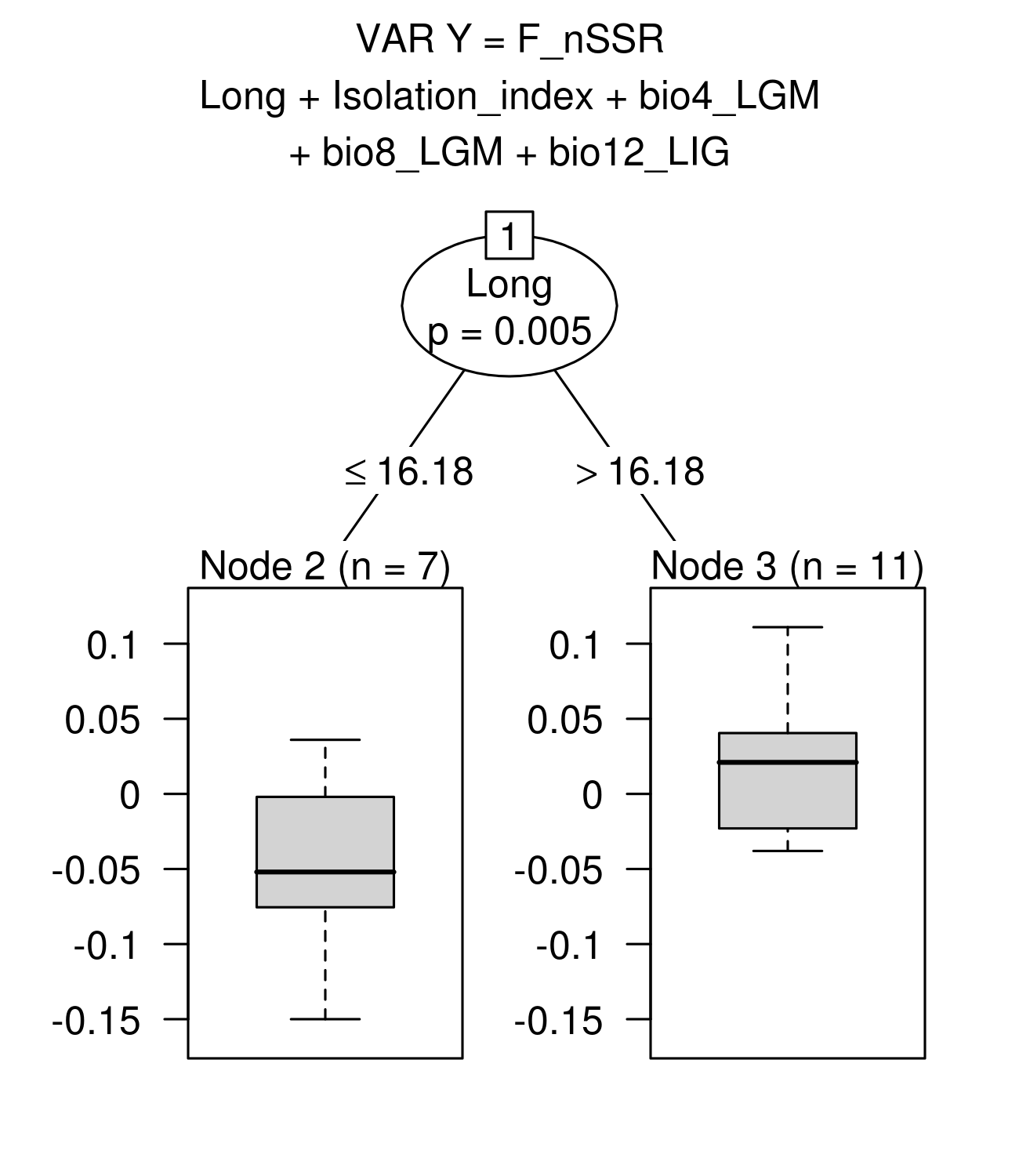

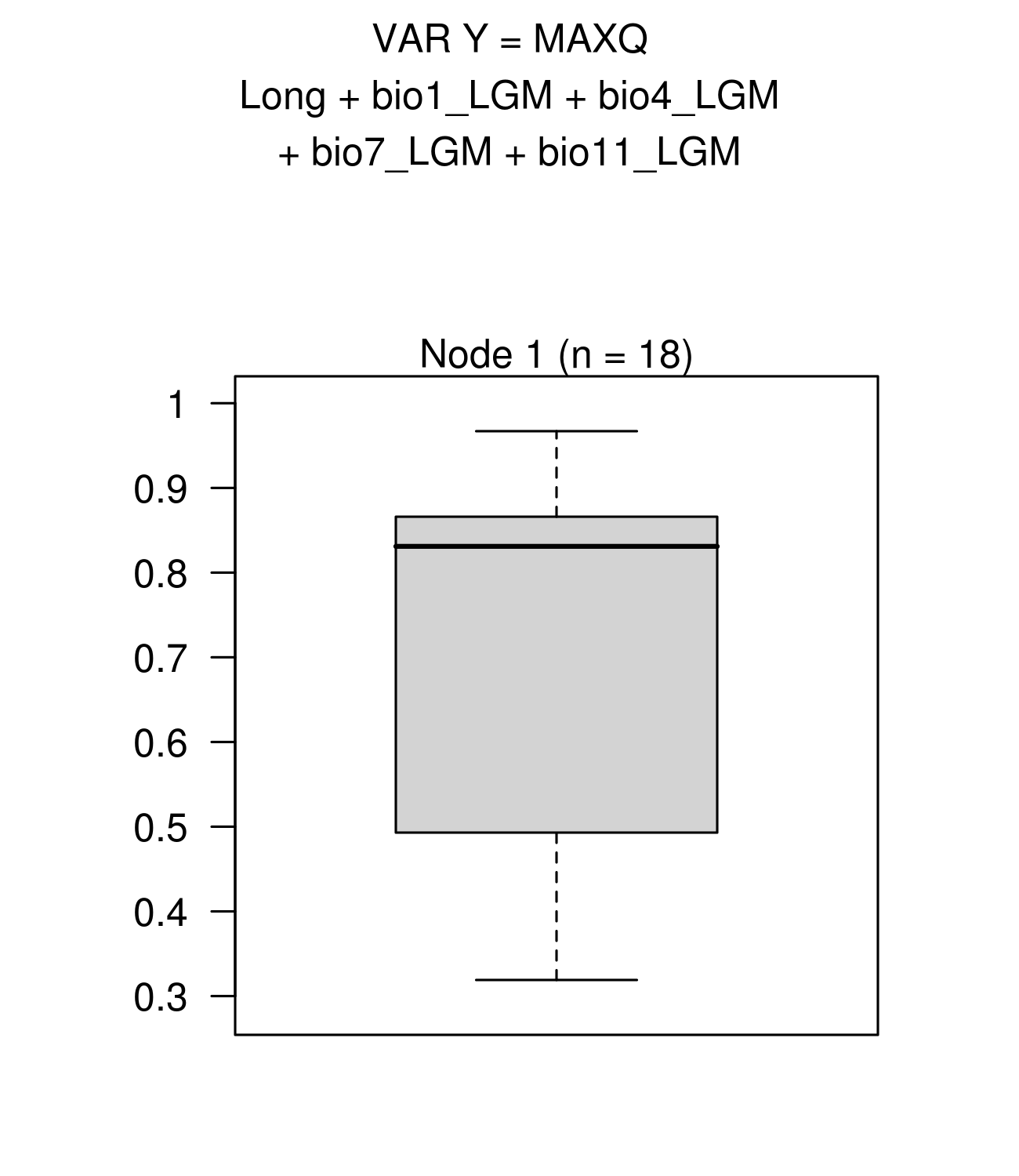

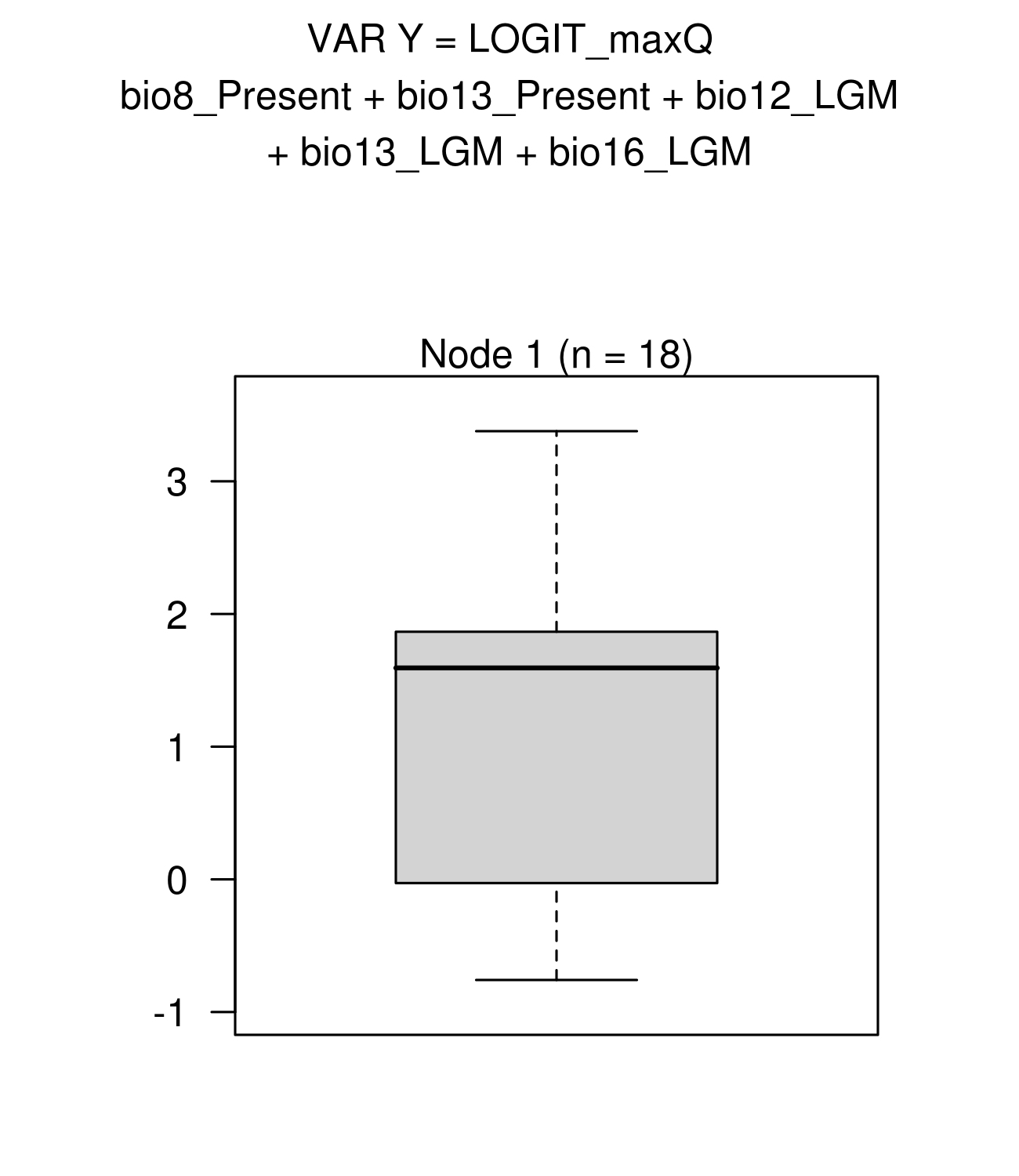

Hothorn, T., Buhlmann, P., Dudoit, S., Molinaro, A. & van der Laan, M.J. (2006a). Survival Ensembles. Biostatistics 7: 355–373.

Hothorn, T., Hornik, K. & Zeileis, A. (2006b). Unbiased Recursive Partitioning: A Conditional Inference Framework. J. Comput. Graph. Stat. 15: 651–674.

Strobl, C., Boulesteix, A.L., Kneib, T., Augustin, T. & Zeileis A. (2008). Conditional variable importance for random forests. BMC Bioinformatics 9: 1–11. https://doi.org/10.1186/1471-2105-9-307

Strobl, C., Boulesteix, A.-L. Zeileis, A. & Hothorn, T. (2007). Bias in Random Forest Variable Importance Measures: Illustrations, Sources and a Solution. BMC Bioinformatics 8: 25.

**Appendix *S4:*** genetic structure among populations

**Figure S4.1***: T*est of genetic structuring at nSSR.

**Figure S4.2**: Test of genetic structuring at cpSSR.

**Figure S4.3:** STRUCTURE analysis. Estimation of the best number of genetic clusters.

**Figure S4.4**: STRUCTURE analysis. Barplots of STRUCTURE ancestry proportions for genetic clusters averaged over 10 runs from K = 2 to K = 7

**Table S4.1:** Test of genetic structuring and mutation effect on genetic structure at nuclear genes.

***Figure S4.1****: T*est of genetic structuring at nSSR.

***Figure S4.1* a:** Linear regressions of pairwise F_ST_ on geographical distances (Km) - 95 % confidence intervals were obtained from 10000 permutations of population locations among all populations. The regression slope was positive and significant (slope ln(dist)= 0.023, pvalue=0)). This test is equivalent to a Mantel test between a matrix of genetic distances and a matrix of geographic distances.

14 nSSR

|  | Fst | CI- | CI+ |
| --- | --- | --- | --- |
| 148.383 | 0.0276479 | 0.0481788 | 0.0872437 |
| 362.032 | 0.0489189 | 0.0453264 | 0.0909443 |
| 542.744 | 0.0409734 | 0.046893 | 0.0898372 |
| 651.299 | 0.0494752 | 0.0486463 | 0.0876722 |
| 726.996 | 0.0529443 | 0.0471177 | 0.090414 |
| 867.548 | 0.0659131 | 0.0481025 | 0.0889405 |
| 965.018 | 0.0526636 | 0.0469379 | 0.090083 |
| 1065.76 | 0.0646096 | 0.0486404 | 0.087456 |
| 1113.99 | 0.0760615 | 0.0485912 | 0.0877414 |
| 1188.83 | 0.0626148 | 0.0467609 | 0.0906788 |
| 1353.27 | 0.0710618 | 0.0442982 | 0.0917146 |
| 1440.44 | 0.0680409 | 0.0478147 | 0.0899653 |
| 1561.56 | 0.0763089 | 0.0463595 | 0.0904587 |
| 1668.28 | 0.0713133 | 0.0497076 | 0.0870848 |
| 1896.29 | 0.0871405 | 0.0492256 | 0.0881251 |
| 2112.77 | 0.0905453 | 0.0493953 | 0.0881114 |
| 2288.42 | 0.103489 | 0.0493717 | 0.088226 |
| 2665.68 | 0.0895387 | 0.0447601 | 0.0911193 |
| 3164.95 | 0.099651 | 0.0403088 | 0.0966475 |

***Figure S4.1* b***:* Test of mutation effect on genetic structure (test of phylogeographic signal) at nSSR loci. Linear regressions of pairwise R_ST_ on geographical distances (Km). 95 % Confidence intervals were obtained after 10000 permutation of allele sizes among alleles within locus. The regression slope was not significant (slope(ln(distance) =0.025, pvalue=0.19)

14 nSSR

|  | Rst | CI-AllelesPermut | CI+ AllelesPermut |
| --- | --- | --- | --- |
| 148.383 | 0.0425596 | -0.00438917 | 0.051568 |
| 362.032 | 0.0587741 | 0.00557938 | 0.0952456 |
| 542.744 | 0.0973403 | 0.00388511 | 0.0787073 |
| 651.299 | 0.0650428 | 0.00533968 | 0.0813923 |
| 726.996 | 0.023703 | 0.00535354 | 0.0771097 |
| 867.548 | 0.088495 | 0.0194516 | 0.100866 |
| 965.018 | 0.0540279 | 0.0124755 | 0.0906508 |
| 1065.76 | 0.0891913 | 0.0131662 | 0.08666 |
| 1113.99 | 0.0876462 | 0.0203254 | 0.0993339 |
| 1188.83 | 0.122314 | 0.00722246 | 0.100424 |
| 1353.27 | 0.0714768 | 0.00967456 | 0.0964401 |
| 1440.44 | 0.13456 | 0.0141532 | 0.101012 |
| 1561.56 | 0.142121 | 0.010927 | 0.0915204 |
| 1668.28 | 0.119415 | 0.012424 | 0.107451 |
| 1896.29 | 0.135402 | 0.0180636 | 0.136257 |
| 2112.77 | 0.0753078 | 0.0102176 | 0.107928 |
| 2288.42 | 0.129429 | 0.0207387 | 0.139134 |
| 2665.68 | 0.0968482 | 0.0133563 | 0.128119 |
| 3164.95 | 0.0775039 | 0.00163624 | 0.162954 |

**Figure S4.2**: Test of genetic structuring at cpSSR.

**a.** Test of genetic structuring at cpSSR loci. Linear regressions of pairwise F_ST_ on geographical distances (Km) - 95 % confidence intervals were obtained from 10000 permutations of population locations among all populations. The regression slope was positive but not significant (slope ln(dist)= 0.008, pvalue=0.16).

cpSSR

|  | Fst | 95%CI-inf | 95%CI-sup |
| --- | --- | --- | --- |
| 233.751 | 0.0501139 | 0.0316874 | 0.0786353 |
| 478.106 | 0.0435093 | 0.0251115 | 0.0867513 |
| 581.663 | 0.0428294 | 0.028501 | 0.0854545 |
| 690.966 | 0.0397046 | 0.0312099 | 0.0785164 |
| 789.291 | 0.0645602 | 0.0277207 | 0.0841079 |
| 920.381 | 0.0332355 | 0.0293036 | 0.0808493 |
| 1035.82 | 0.0400955 | 0.0279806 | 0.0853505 |
| 1085.91 | 0.0342121 | 0.0301078 | 0.0775653 |
| 1144.78 | 0.0472542 | 0.0315879 | 0.0784424 |
| 1283.92 | 0.0430438 | 0.0273478 | 0.0877764 |
| 1363.58 | 0.0830757 | 0.0248506 | 0.0895351 |
| 1528.43 | 0.0447096 | 0.0284679 | 0.0828659 |
| 1599.88 | 0.0560327 | 0.0273368 | 0.0838918 |
| 1743.58 | 0.0446201 | 0.0334272 | 0.0773969 |
| 2036.6 | 0.0657051 | 0.0312712 | 0.0790521 |
| 2189.84 | 0.0827445 | 0.0310833 | 0.0788309 |
| 2437.36 | 0.0448039 | 0.0311137 | 0.0791893 |
| 2793.42 | 0.0832109 | 0.0250807 | 0.0880762 |
| 3404.41 | 0.0795727 | 0.0187492 | 0.100327 |

***b.*** Test of mutation effect on genetic structure (test of phylogeographic signal) at cpSSR loci. Linear regressions of pairwise N_ST_ on geographical distances (Km) -95 % Confidence intervals are obtained after 10000 permutation of permutation of rows and columns of distance matrices between alleles. The regression slope was positive but not significant (slope ln(dist)= 0.009, pvalue=0.84).

cpSSR

|  | Nst | 95%CI-inf | 95%CI-sup |
| --- | --- | --- | --- |
| 233.751 | 0.15812 | 0.0190859 | 0.0831757 |
| 478.106 | 0.0897162 | 0.0103208 | 0.0776169 |
| 581.663 | 0.0704333 | 0.0114442 | 0.0765266 |
| 690.966 | 0.151277 | 0.0109044 | 0.0708622 |
| 789.291 | 0.0842371 | 0.0297728 | 0.101922 |
| 920.381 | 0.117719 | 0.00461489 | 0.0639382 |
| 1035.82 | 0.108089 | 0.00911795 | 0.0750608 |
| 1085.91 | 0.131971 | 0.00669171 | 0.0639728 |
| 1144.78 | 0.133541 | 0.0183421 | 0.0778366 |
| 1283.92 | 0.0856237 | 0.0115868 | 0.0792138 |
| 1363.58 | 0.139969 | 0.0415853 | 0.126998 |
| 1528.43 | 0.134594 | 0.0128663 | 0.0809535 |
| 1599.88 | 0.113519 | 0.022923 | 0.0919116 |
| 1743.58 | 0.14058 | 0.015335 | 0.0759809 |
| 2036.6 | 0.140289 | 0.0314246 | 0.10188 |
| 2189.84 | 0.189688 | 0.0442906 | 0.124226 |
| 2437.36 | 0.129873 | 0.0145665 | 0.0786451 |
| 2793.42 | 0.160756 | 0.0421378 | 0.127749 |
| 3404.41 | 0.14054 | 0.0313441 | 0.134409 |

**Figure S4.3*:*** STRUCTURE analysis. Estimation of the best number of genetic clusters. Characteristics of the black pine STRUCTURE analysis at 14 nSSR, 14 nuclear genes and 1 cpSSR. The genetic structure at 233 individuals sampled in 19 populations was assessed using an admixture model with correlated allele frequencies with five independent runs for each K value ranging from 1 to 10 , a burn-in period of 5x10^5^ steps followed by 1x10^6^ Markov chain Monte Carlo replicates.

**Figure S4.3 *a:*** Posterior log-likelihood of data for K clusters

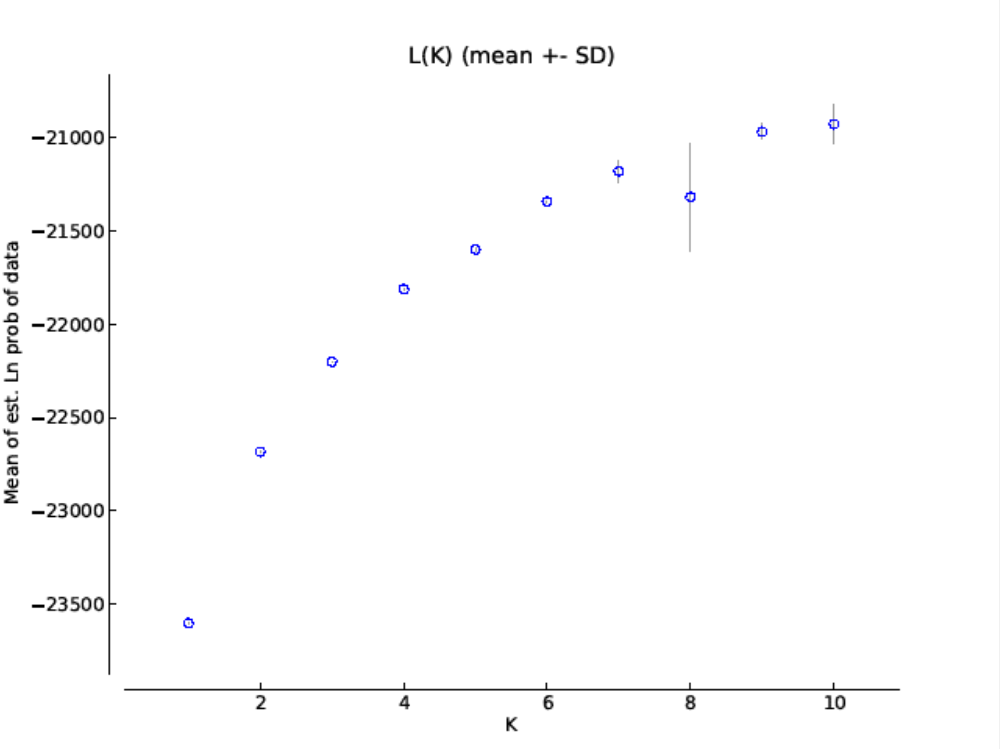

**Figure S4.3 b**: Application of the ad hoc method of Evanno et al. (2005) to identify the best K.

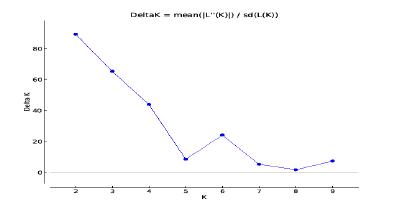

**Figure S4.4**: Structure analysis in the 19 populations of *Pinus nigra*. Barplots of STRUCTURE ancestry proportions for genetic clusters averaged over 10 runs from K = 2 to K = 7. Each individual is represented by a vertical bar divided into colour segments representing the gene pools identified by STRUCTURE. The 7 most homogeneous populations used in the ABC analysis are indicated in bold.

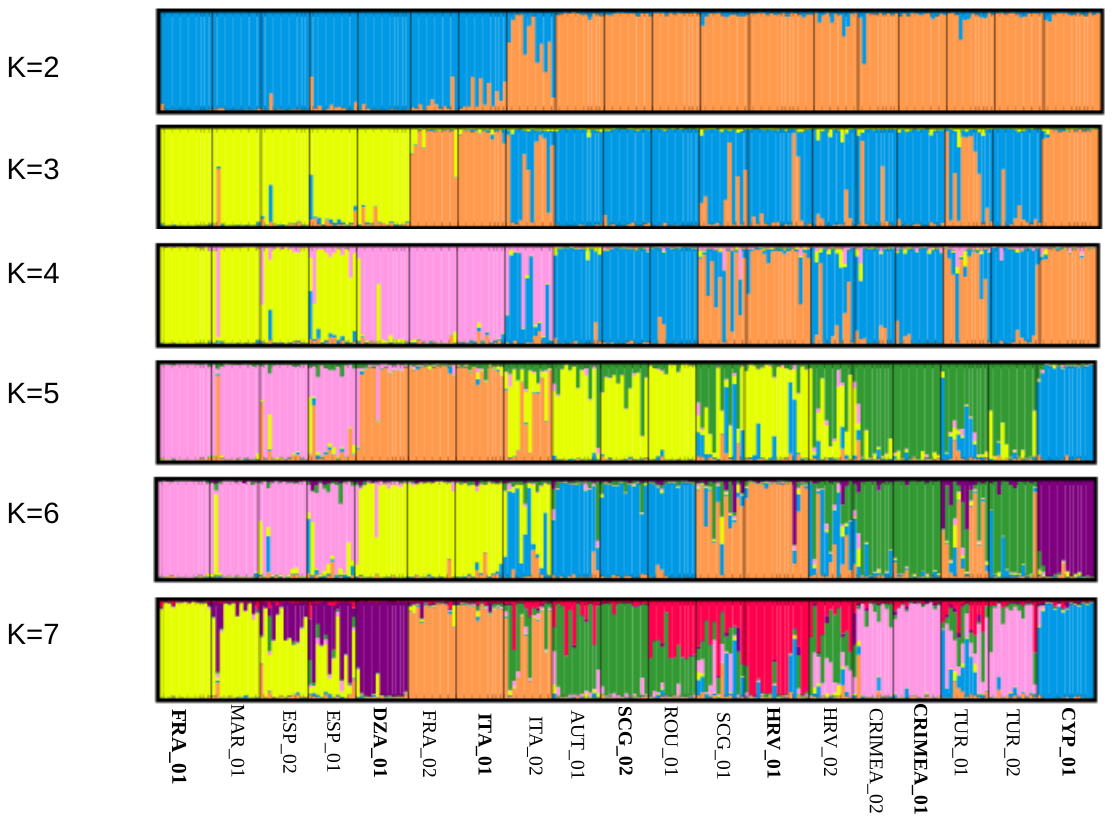

***Table S4.1:*** Test of genetic structuring and mutation effect on genetic structure at nuclear genes.

Linear regressions slope of pairwise F_ST_ on geographical distances (Km) and Pvalue obtained after 10000 permutations of population locations among all populations. Linear regressions slope of pairwise N_ST_ on geographical distances (Km) and Pvalue obtained after 10000 permutation of rows and columns of distance matrices between alleles.

|  | **F_ST_** | | **N_ST_** | |
| --- | --- | --- | --- | --- |
|  | **slope** | **Pvalue** | **slope** | **Pvalue** |
| Gene_1405 | 0.032 | 0.52 | 0.033 | 0.99 |
| Gene_2078 | 0.005 | 0.80 | -0.002 | 0.80 |
| Gene_6293 | 0.035 | **0.018** | 0.018 | 0.46 |
| Gene_7916 | -0.01 | 0.58 | -0.02 | 0.51 |
| Gene_8479 | 0.00007 | 0.76 | -0.01 | 0.18 |
| Gene_10162 | 0.047 | **0.0065** | 0.05 | 0.70 |
| Gene_10384 | 0.14 | **0.0049** | 0.125 | 0.52 |
| Gene_10667 | 0.043 | 0.11 | 0.01 | 0.12 |
| Gene_13484 | - | 1 | 0.12 | 0.31 |
| Gene_13957 | 0.02 | 0.30 | 0.03 | 0.56 |
| Gene_14221 | 0.05 | **0.03** | 0.03 | 0.23 |
| Gene_16810 | 0.057 | **0.0099** | 0.057 | 0.71 |
| Gene_18101 | 0.029 | 0.109 | 0.023 | 0.75 |
| Gene_CL4470 | -0.047 | 0.27 | -0.046 | 0.61 |
